# Centimeter-scale quorum sensing dictates collective survival of differentiating embryonic stem cells

**DOI:** 10.1101/2020.12.20.423651

**Authors:** Hirad Daneshpour, Pim van den Bersselaar, Hyun Youk

## Abstract

Cells can help each other replicate by communicating with diffusible molecules. In cell cultures, molecules may diffuse within a cell colony or between adjacent or distant colonies. Determining which cell helps which cell’s replication is challenging. We developed a systematic approach, integrating modeling and experiments, for determining the length-scales of cell-cell communication (from microns to centimeters). With this approach, we discovered that differentiating murine ES cells, scattered across centimeters on a dish, communicate over millimeters to form one macroscopic entity that survives if and only if its centimeter-scale population-density is above a threshold value. Single-cell-level measurements, transcriptomics, and modeling revealed that this “macroscopic quorum sensing” arises from differentiating ES cells secreting and sensing survival-promoting FGF4 that diffuses over millimeters and activates YAP1-induced survival mechanisms. Through the same mechanism, a lone macroscopic, but not microscopic, colony survives differentiation. Our work rigorously establishes that *in vitro* ES-cell differentiation relies on macroscopic cooperation.

## INTRODUCTION

A cell can autonomously replicate, without interacting with other cells. But autonomously replicating does not guarantee that a cell population grows. A cell population becomes extinct when more cells die than replicate. Cells can prevent a population extinction by helping each other’s replication (i.e., increase the success rate of cell replication) (*1,2*). Cells typically do this by secreting and sensing the same, diffusible molecule (*3–8*). In such “autocrine signaling” with molecules (growth factors) such as FGF and EGF (*9,10*), a cell senses the molecule and then tunes its intracellular processes (e.g., gene expression) to facilitate its growth and replication (*6*). Since a cell can secrete and capture a molecule before it diffuses to another cell, autocrine signaling enables a cell to communicate with itself (self-communicate) (*4,6,7*). A cell can also capture the same molecule, but from another cell, and thereby communicate with another cell (neighbor-communication). Given this complication, several conceptual and technical challenges exist in quantitatively understanding autocrine-based communication among cells in complex, physical architectures (e.g., colonies of various sizes distributed over a vast area) (*6,11*). These challenges include resolving the following questions: which cell secretes a particular molecule, which cell captures that molecule, how many copies of the molecule is diffusing between cells, how many different types of molecules are used for communication, how far the molecules diffuse, how much time elapses between the molecule secretion and gene-expression changes in the molecule-sensing cell, whether two adjacent cells communicate with each other given that both cells may only self-communicate (*4,6,7*), and which intracellular pathways are regulated by the autocrine molecules. Even when we know that cells communicate, it is unclear whether they do so only in the confines of their microscopic environment (within a colony or cell aggregate) (*1,12,13*) or over macroscopic distances that span millimeters to centimeters on a cell-culture dish (*14,15*). Conventionally, one transfers conditioned media (*16*) or uses microfluidics (*17,18*) to determine whether autocrine signals exist in cell cultures. In a medium-transfer experiment, one pools together molecules from everywhere on the dish, uniformly mixes them, and then transfers this mixture to another dish of cells to determine whether secreted molecules exist in the original medium. With microfluidics, one constantly flows away any cell-secreted molecules along one direction, thereby accumulating any secreted molecules with a gradient along the flow direction. Hence both the medium-transfer and microfluidics methods destroy the crucial, spatial information regarding cell-cell communication. To address the questions mentioned above, we need an approach that leaves cell-cell communication and the paths of diffusing molecules undisturbed.

In this article, we developed a systematic approach that addresses the challenges mentioned above for cell cultures by integrating quantitative experiments with mathematical analyses of various communication distances. As a demonstration, we applied this integrated approach to a culture of mouse Embryonic Stem (ES) cells. This led us to discover that ES cells quorum sense at a near-centimeter scale as they exit pluripotency and begin differentiating into all lineages. This macroscopic quorum-sensing dictates whether the entire population, spanning many centimeters on a dish, survives or becomes extinct. Despite their inability to capture many *in vivo* phenomena, ES-cell cultures are ideal testbeds for investigating autocrine signaling because these cells are known to depend on myriad, secreted molecules (e.g., FGF and Wnt ligands) (*19–28*). ES cells also exhibit cell-density dependent effects (*29,30*). In ES-cell cultures, quantifying the spatial range of communication and determining how communication affects a cell’s gene expression are important for building artificial tissues (*31–34*) and embryo-like structures (*35–40*) and for synthetic biology (*41–46*). More generally, given that ES-cell cultures are frequently used to reveal basic principles of cell signaling, it is important to determine which molecules from which cell are received by an ES cell on a dish. By developing a systematic, quantitative approach that is applicable to any cell cultures, we quantified the degree to which an ES cell is autonomous and determined, for each length-scale, how much cell-cell communication occurs and its impact on differentiation.

### Outline of this article

We begin with mathematical models showing two possible scenarios of collective growth: (1) population grows faster as more cells help each other, and (2) population only grows, instead of becoming extinct, if and only if the initial population-density is beyond a threshold value (Fig. 1). We then experimentally show that ES cells collectively survive with a density threshold while differentiating towards either neural ectoderm or mesendoderm lineages. In contrast, pluripotent ES cells do not show signs of collective growth with a density threshold (Fig. 2). We find that, near the threshold density, whether the population grows or becomes extinct is stochastically determined, as predicted by a stochastic version of our model (Fig. 3). We then demonstrate that secreted molecules are responsible for the observed phenomenon (Fig. 3) and that these molecules, while allowing for local communication (e.g., communication within a colony or between colonies within 1-mm range), local communication alone is insufficient to yield the collective growth (Fig. 4). We performed a systematic analysis in which we manipulated the volume of the liquid cell-culture medium, measured the stability of secreted survival factors, determined the range of molecular weights of the secreted survival factors, and compared the results of these measurements with a mathematical model. These established that the secreted survival factors diffuse over nearly a centimeter scale (Fig. 5). Thus, for practical consideration, we can consider these molecules to be uniformly mixed. We then reveal a mechanistic picture that underlies the centimeter-scale quorum sensing: FGF4 is secreted by differentiating ES cells and FGF4 is both sufficient and necessary for the collective growth with the observed density-threshold (Fig. 6). Our article ends with a practical implication for tissue engineering and cell cultures: seeding a macroscopic cluster of cells (cells dropped into a millimeter-scale area) leads to everyone surviving whereas dispersing the same number of cells over the centimeter-scale dish causes the population to become extinct (Fig. 7). The macroscopic clustering also leads to the highest fraction (more than 90%) of cells successfully differentiating among all the culturing methods that we examined. We expect that this result will inform the engineering of macroscopic (millimeter-to-centimeter scale), artificial tissues and multicellular entities with ES cells in cell cultures. From the perspective of basic science (physical biology and statistical physics), our work showcases a spatially extended system, composed of microscopic cells and a macroscopic environment that feedback on each other, that uses molecular diffusion to bridge vast length scales and keeps itself out of thermal equilibrium (death). We expect that further studies of this macroscopically quorum-sensing system will help us reveal physical principles that govern out-of-equilibrium dynamics of macroscopic, reaction-diffusion systems that are composed of living cells (*41,42,47–50*).

**Figure 1.**
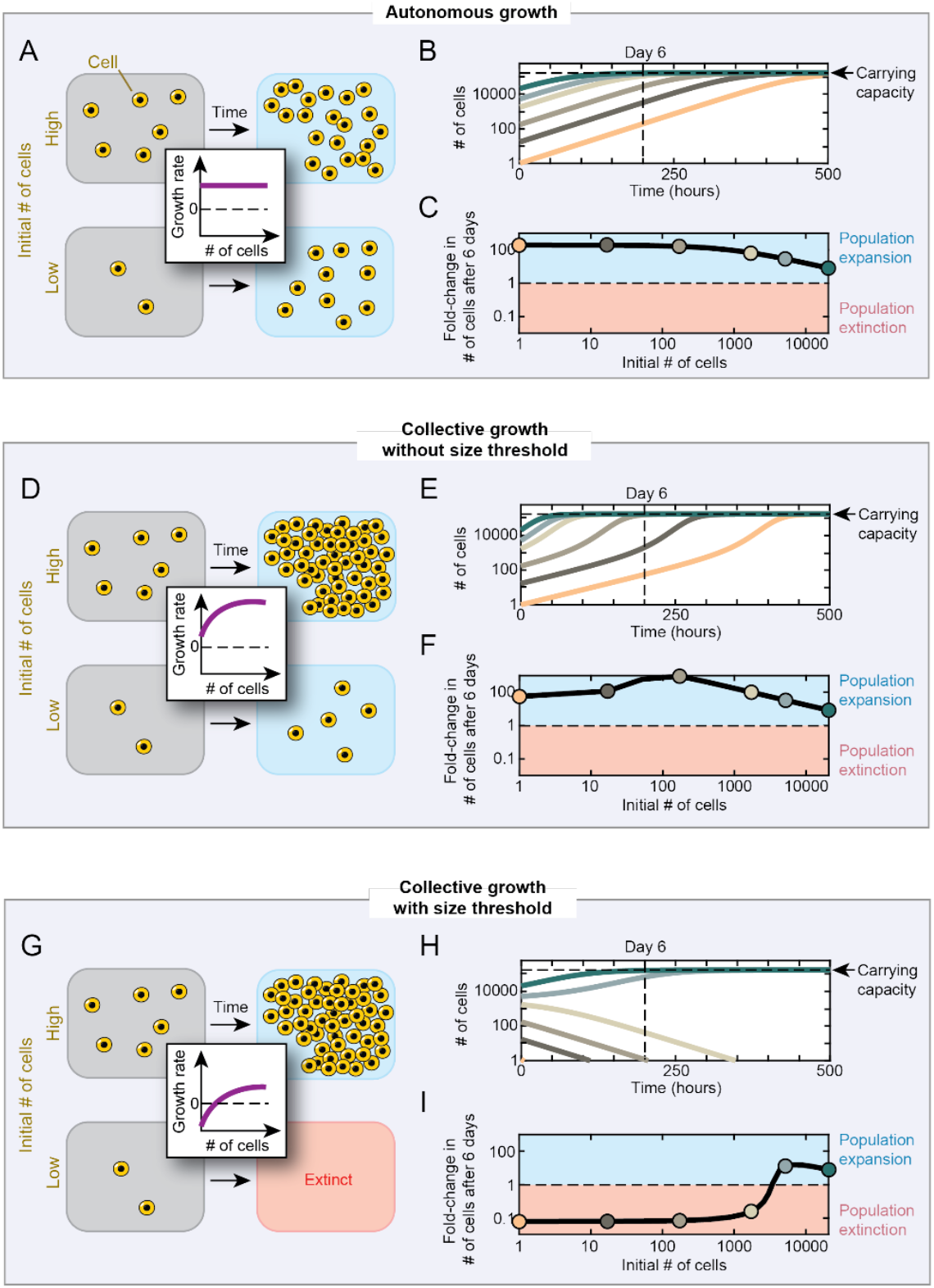
Modeling autonomous and collective growths of cells. Cell populations grow according to the logistic equation and the net growth rate of the cells follows a Michaelis-Menten relation depending on the instantaneous number of cells (see STAR Methods). (**A**) Cartoon showing cells at any initial number autonomously growing toward the carrying capacity with the same growth rate (independent of initial number of cells). (**B**) Model-produced curves for autonomous growth. Populations that start with 1 cell (light orange) or 20,690 cells (dark green) all grow toward the carrying capacity (vertical dashed line). (**C**) Model-produced fold-change in initial number of cells after 6 days for autonomous growth. Any initial number of cells results in expansion of the population (blue shade). (**D**) Cartoon showing cells at any initial number collectively growing toward the carrying capacity, albeit with a higher net growth rate for populations that start at higher initial numbers. (**E**) Model-produced curves for collective growth without density threshold. Populations that start with 1 cell (light orange) or 20,690 cells (dark green) all eventually grow toward the carrying capacity (vertical dashed line). (**F**) Model-produced fold-change in initial number of cells after 6 days for collective growth without density threshold. Any initial number of cells results in expansion of the population (blue shade). (**G**) Cartoon showing cells that start with sufficiently many of them survive and expand, otherwise they become extinct. (**H**) Model-produced curves for collective growth with density threshold. Populations that start with at least ~3400 cells survive and expand toward the carrying capacity. (**I**) Model-produced fold-change in initial number of cells after 6 days for collective growth with density threshold. Populations that start with at least ~3400 cells survive and expand toward the carrying capacity (blue shade), otherwise they become extinct after 6 days (red shade).

## RESULTS

### Modeling autonomous and collective growths

We consider three scenarios: (1) cells autonomously replicate and die, (2) cells help each other replicate and a lone cell is more likely to replicate than die, and (3) cells help each other replicate and a lone cell is more likely to die than to replicate. In scenarios (2) and (3), cells collectively grow. We built a mathematical model for each scenario. The only difference among the models is how the net growth rate, which is the growth rate minus the death rate, changes as a function of the number of cells in a population (“population size”).

We modeled the autonomous growth (Scenario (1)) by letting the net growth rate to be a positive constant (Fig 1A - purple line) (see STAR Methods). Here, regardless of its initial value, the population size exponentially increases over time until it reaches the carrying capacity (Fig. 1B). The fold-change in the population size, relative to the initial population size, after a fixed time (6 days) is the same for a range of initial population sizes (Fig. 1C - first three points) and is less for populations that were initially large enough to already reach the carrying capacity within the 6 days (Fig. 1C - last three points).

In scenario (2), the net growth rate is positive when there is just one cell because the cell has a higher probability of autonomously replicating than dying. The net growth rate increases as the population size increases (Fig. 1D - purple curve). In this scenario, populations grow faster if they start with more cells and all populations eventually reach the carrying capacity regardless of their initial sizes (Fig. 1E and STAR Methods). Hence, the fold-change in the population size, relative to the initial population size, after a fixed time (6 days) is larger than one for all initial population sizes. For this reason, we call this scenario to exhibit a “collective growth without size threshold”.

In scenario (3), the net growth rate increases as the population size increases but it is negative (more deaths than replications) when the population size is below a “size threshold” (Fig. 1G - purple curve). For population sizes above the size threshold, the net growth rate is positive because there are enough cells helping each other (Fig. 1G - purple curve above dashed line). Here, simulations show that a population becomes extinct if its initial size is below the size threshold (Fig. 1H - bottom three curves) whereas it grows to reach the carrying capacity if its initial size is above the size threshold (Fig. 1H - top three curves). Hence, the fold-change in the population size, relative to the initial population size, after a fixed time (6 days) sharply increases in a switch-like manner, from nearly zero to above one, as the initial population size crosses the size threshold (Fig. 1I).

### Setup of ES-cell cultures and differentiation protocols

To determine which of the three growth scenarios best describes murine ES cells, we counted the number of ES cells in suspension and then randomly scattered them across a large (10-cm diameter), gelatin-coated dish without any feeder cells. The resulting, initial population-density (# of cells / cm^2^) ranged from ~5 cells / cm^2^ to ~15,000 cells / cm^2^ (Fig. 2A – dashed box). To count the cells, we used a flow cytometer as well as a hemocytometer with trypan-blue staining (for counting only the viable cells). Both methods yielded similar cell counts (Fig. S1).

**Figure 2.**
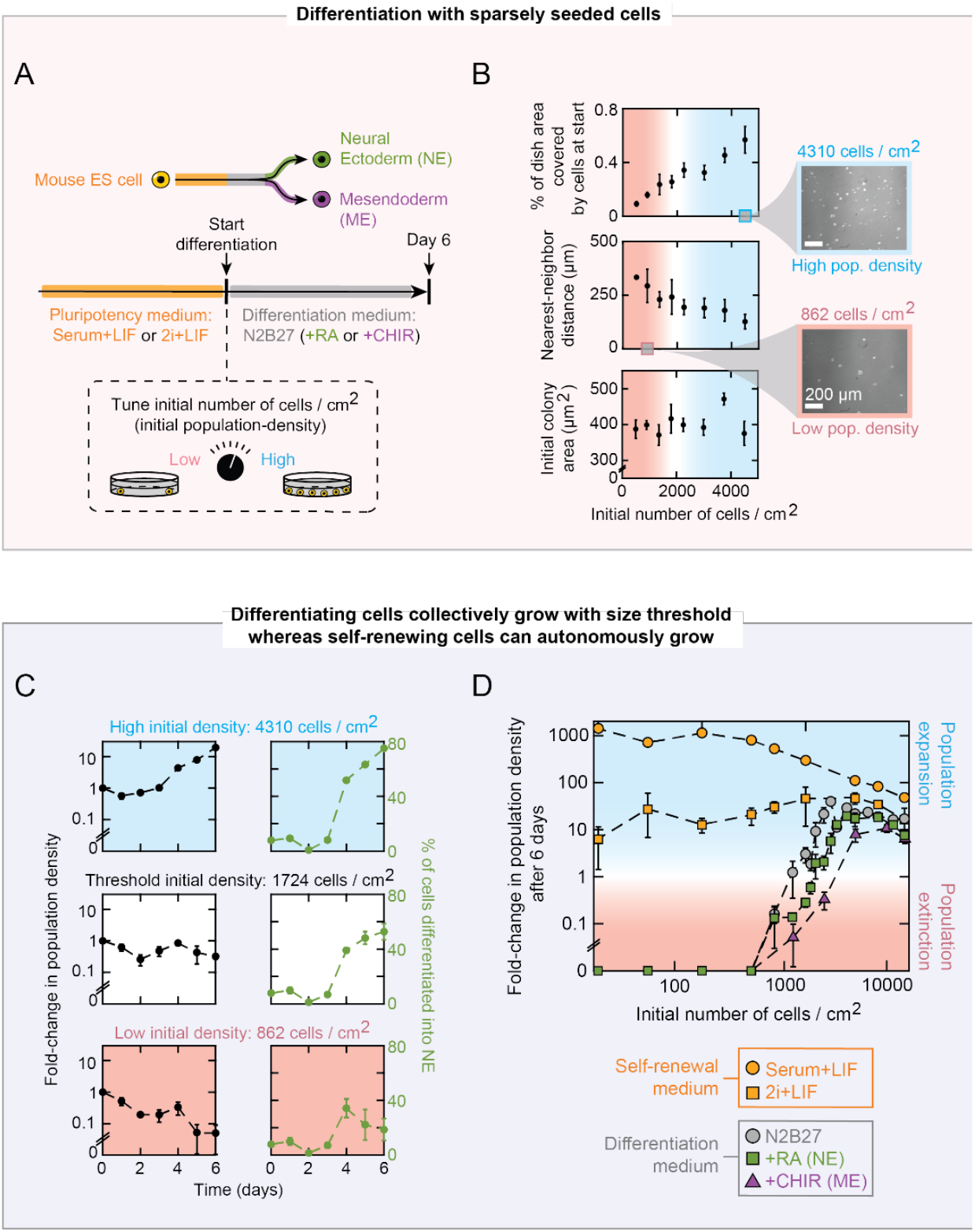
ES cells collectively grow with a size threshold during differentiation but not during self-renewal. (**A**) Differentiation protocol used in this study (see STAR Methods). (**B**) As a function of initial number of E14 cells seeded on a 10-cm diameter dish when cultured in N2B27 to trigger differentiation: percentage of the plate area initially covered by microcolonies (top graph), initial distance to a nearest neighbor colony on average (middle graph), initial area of a microcolony on average (bottom graph). Microscope used to measure all three parameters. Blue shade indicates population expansion. Red shade indicates population extinction. Two example microscope images shown. Scale bar = 200 μm. *n* = 3; Error bars are s.e.m. (**C**) Fold-change in population density (black) and percentage of cells entering Neural Ectoderm (NE) lineage (i.e., percentage of cells expressing Sox1-GFP) (green) as a function of time after RA-induced differentiation starts. Indicated above each row is the initial population density for the population shown in that row (see also Figs. S3–6). 46C cells are used for differentiation in N2B27 + Retinoic Acid (RA), previously self-renewing in serum with LIF. Also see Fig. S2 for confirmation of differentiation into NE lineage regardless of the use of inducers or serum. *n* is at least 3. Error bars are s.e.m. (**D**) As a function of initial population density, fold change in population density after 6 days in pluripotency medium (orange) or one of three differentiation media (grey - N2B27 without any inducers; green - N2B27+RA that induces NE-lineage differentiation; purple - N2B27+CHIR that induces Mesendoderm (ME) lineage differentiation). Cell lines used: E14 (orange and grey), 46C (green), and Brachyury-GFP (purple). See Fig. S6 for all cell lines and media types (with or without any inducer or prior use of serum). Blue shade indicates population expansion. Red shade indicates population extinction. *n* is at least 3; Error bars are s.e.m.

We cultured ES cells in two conditions: self-renewing (i.e., remaining pluripotent) or differentiating (i.e., exiting pluripotency). For self-renewal, we cultured ES cells in either a medium with a serum (FBS) or without serum (2i), both with Leukemia Inhibitory Factor (LIF) (Fig. 2A - yellow). As we will show, our paper’s main conclusions hold for both media. For differentiation, we cultured the cells in three kinds of media, all lacking serum and LIF: N2B27, N2B27 with Retinoic Acid (RA), and N2B27 with CHIR (a small molecule). As we will show, our paper’s main results on differentiation hold for all these media. Absence of LIF causes the cells to exit pluripotency in all three media (Fig. 2A - grey). In accordance with literature (*51,52*), we verified that N2B27 without any inducers (RA or CHIR) caused a large majority of ES cells (more than 80%) to differentiate into the Neural Ectoderm (NE) lineage, regardless of which medium (2i or serum) the ES cells were previously self-renewing in (Fig. S2). N2B27 with RA also induces NE-lineage differentiation (*53,54*) (Fig. 2A - green). N2B27 with CHIR induces ME-lineage differentiation (*56*) (Fig. 2A – purple). Following existing protocols (*53*), whenever we used either RA or CHIR, we first cultured the ES cells in N2B27 without any inducers for two days (Fig. 2A – “Start of differentiation”) and then added either RA or CHIR at the end of the two days. The two days are required for most of the ES cells to degrade their pluripotency factors (e.g., Oct4) to exit pluripotency and be responsive to RA or CHIR (*53*). To not disturb the paths of any diffusing factors that cells might be secreting, we did not shake or stir the cell-culture medium or move the dishes during the entire incubation period. Only when we were ready to measure the population density, we moved the dish, detached all the cells from it, then counted the number of cells. We thereby sacrificed a dish for each measurement of population density.

Our cell-seeding method yielded sparsely distributed, nearly single cells on a dish for every population. For every initial population-density, imaging a dish with a microscope revealed that cells covered less than 1% of the dish area (Fig. 2B - top). From the microscope images, we obtained the distance between a colony and its nearest-neighboring colony, for multiple colonies and fields of view in every initial population-density. The distance between a colony and its nearest-neighboring colony decreased as the initial population-density increased but it was, on average, ~100 μm or more for every population density (Fig. 2B - middle). The initial colony-area, averaged over many colonies, was virtually identical for every initial population-density (Fig. 2B – bottom). Hence, larger initial population-density did not yield larger colonies.

### ES cells collectively grow with a size threshold during differentiation but not during self-renewal

We first examined the differentiation into the NE lineage with RA. Here, we used a “46C” cell line which has a Sox1 promoter controlling GFP expression (*51*). Sox1 is a marker of the NE lineage. Hence ES cells that entered the NE lineage would express GFP. With a flow cytometer, in each population and on different days, we counted and discriminated GFP-expressing cells (NE cells) from those that did not (Fig. 2C – green curves). A higher initial population-density led to a higher fraction of cells becoming an NE cell (Fig. 2C - right column; also Fig. S3–S4). By counting the number of cells, we also obtained a fold-change in population density relative to the initial density on different days (Fig. 2C – black curves). Populations that began with a sufficiently high density (above ~1700 cells / cm^2^) grew towards the carrying capacity (Fig. 2C - top row) whereas populations that began with a sufficiently low density (below ~1700 cells / cm^2^) approached extinction over six days (Fig. 2C - bottom row). A population that nearly began with a “threshold density” (~1700 cells / cm^2^) neither noticeably grew nor shrank during the first six days (Fig. 2C - middle row). But, days later, it either grew towards the carrying capacity or shrank towards extinction (Fig. S5). Specifically, by using an ensemble of many dishes that all started with the same density, we discovered that two populations that start with the same, near-threshold density can have different outcomes: one expanding and one becoming extinct.

We found the same, population-level phenomenon (collective growth with a density threshold) arising in three different, widely used cell lines: E14 cell line (Fig. 2D - grey), 46C cell line (Fig. 2D - green), and a cell line with the promoter of Brachyury, a marker of ME lineage (*57,58*), controlling GFP expression (Fig. 2D – purple). The same phenomenon arises in every (three) differentiation media: N2B27 (Fig. 2D - grey), N2B27 with RA (Fig. 2D - green), and N2B27 with CHIR (Fig. 2D - purple). The same phenomenon also arises whether the ES cells were self-renewing in 2i or serum before differentiating (Fig. S6). In every cell-line and medium, a differentiating population’s survival-versus-extinction fate can hinge on a mere two-fold difference in the initial density. For example, with 46C cells, ~1500 cells / cm^2^ leads to an extinction whereas ~3000 cells / cm^2^ leads to an expansion towards the carrying capacity. The threshold density differs by at most two-fold among three cell-lines and differentiation media (Fig. 2D). From here on, unless stated otherwise, we will focus on the 46C cell-line differentiating in N2B27 with RA after self-renewing in the serum medium. This is because the main phenomenon, collective growth with a size (density) threshold, arises for every cell line, for both (NE and ME) lineage differentiations, for every differentiation media, and regardless of the self-renewing condition (either in serum or 2i) prior to the differentiation.

Self-renewing ES cells did not show signs of collective growth with a density threshold in the wide range of population densities that we examined (~5 to ~15,000 cells / cm^2^). Self-renewing populations of all initial densities expanded towards the carrying capacity, both in serum (Fig. 2D – orange circles) and in 2i (Fig. 2D – orange squares). Notably, cells scattered around the dish at densities as low as ~5 cells/cm^2^ grew by ~1000 fold in six days (Fig. 2D – leftmost data points in orange). Therefore, we conclude that differentiating ES cells collectively grow with a size threshold (matches Fig. 1I) whereas self-renewing ES cells do not.

### Secreted factors dictate collective growth with a density threshold

We hypothesized that differentiating cells secrete at least one factor (“survival factor”) and that cells sensing this factor have an increased chance of replicating. This would lead to a collective growth. If this hypothesis is true, then collecting medium from a high-density population (5172 cells / cm^2^) would pool together the survival factor from everywhere in the dish. Then, giving this medium to a low-density population (862 cells / cm^2^) that was bound for extinction should rescue that population from extinction. We did this medium-transfer experiment in two ways (Fig. 3A). In one method, we collected the high-density population’s medium after X days of differentiation and then, in it, initiated differentiation of low-density population (Fig. 3A - labeled “1”). In this “day X to day 0” transfer, the low-density population was rescued from extinction if and only if X was 2, 3, or 4 (Fig. 3B - left column: black bars show ~4-fold change in population density for X=2, 3, and 4). If X was 5, then the low-density population was barely saved. If X was 1 or 6, the low-density population still went extinct. These results are consistent with the survival factors needing ~2 days to accumulate to a sufficient concentration and degrading over time such that there is not enough of them to rescue populations after ~5 days.

**Figure 3.**
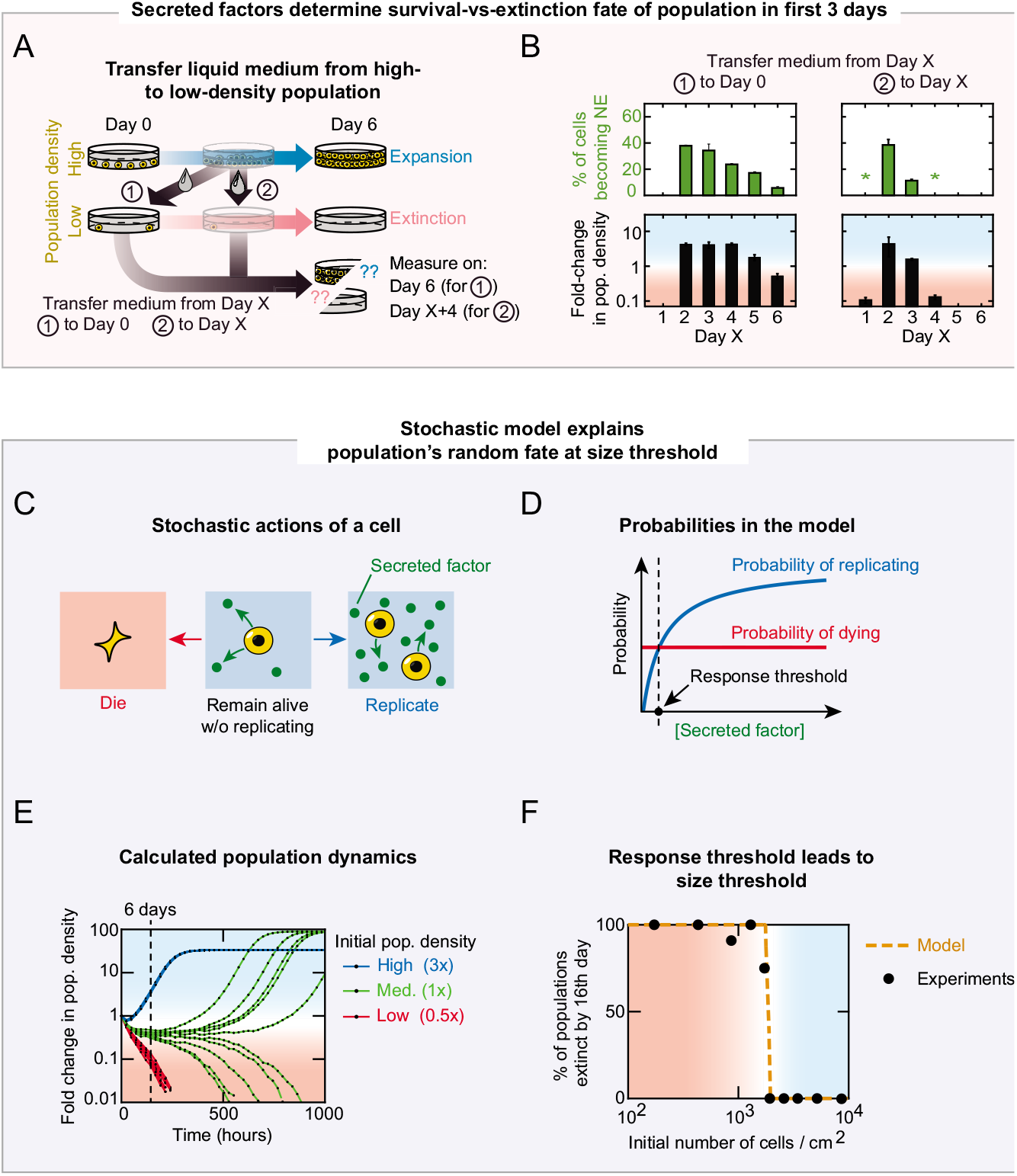
Stochastic model recapitulates stochastic survival-vs-extinction fates of populations at size threshold. (**A**) Over 6 days, high-density population (5172 cells / cm^2^) expands (top: blue arrow) and low-density population (862 cells / cm^2^) approaches extinction (middle: pink arrow). Two methods shown for replacing medium of the low-density population by that of the high-density population. Method (labeled “1”): replace initial medium of low-density population with the medium from X-days-old high-density population. Method 2 (labeled “2”): replace medium of X-days-old low-density population with that of X-days-old high-density population. (**B**) Results of two experiments described in (A). Left column for method 1 and right column for method 2. Data for serum-grown 46C cells induced with RA to enter Neural Ectoderm (NE) lineage. As a function of Day X, green bars show percentage of cells in population becoming NE (expressing Sox1-GFP) and black bars show fold change in population density. Blue shade indicates population expansion and red shade indicates population extinction. Asterisks indicate too few cells for reliable measurement on flow cytometer (nearly or less than 0.1-fold change in population density). Error bars are s.e.m.; *n* = 3. (**C**) Mathematical model with three possible, stochastic actions of a cell (see STAR Methods). (**D**) Probabilities used in the model: probability of a cell dying (red) and probability of cell replicating (blue), both as functions of extracellular concentration of a secreted factor. Probability of dying is constant (see Fig. S8 for experimental validation). The probability of replicating nonlinearly increases with the concentration of extracellular the secreted factor and is based on growth rates of various populations that we measured (see Fig. S7 and S9 and STAR Methods). Vertical dashed line (“response threshold”) indicates the concentration above which cells have a larger probability of replicating relative to the probability of dying. (**E**) Fold-change in population density, simulated by the model. Red: low-density population (862 cells/cm^2^ initially). Green: near-threshold (medium) density population (1931 cells/cm^2^ initially). Blue: high-density population (5172 cells/cm^2^ initially). Vertical, dashed line marks 6 days after differentiation began (for comparison with Fig. S3 and S5). Blue shade indicates population expansion and red shade indicates population extinction. Each density is shown with 10 replicates (simulations). (**F**) Percentage of populations, in an ensemble of populations that start with the same, near-threshold density (1931 cells/cm^2^), that becomes extinct either before or on the 16th day of differentiation. Black data points are from measurements and orange curve is from running the stochastic model multiple times (10 simulations for each population in an ensemble). Subset of *n* = 86 populations shown as black points (see Fig. S5 for full data). Population was considered to have reached extinction if its fold-change decreased to below 0.1, relative to initial density. Blue shade indicates population expansion and red shade indicates population extinction.

As another method, we collected the medium of the high-density population X days after beginning differentiation and then transplanted in it a low-density population that was already differentiating, before being transplanted, for the same number of days (X days) (Fig. 3A - labeled “2”). In this “day X to day X” experiment, the low-density population was rescued from extinction if and only if the X was 2 or 3 (Fig. 3B - right column: black bars show fold-change in population density which is ~4-fold for X = 2 and just above 1-fold for X = 3). These results are consistent with differentiating cells no longer responding to the survival factors starting on the third day of differentiation.

### Stochastic model recapitulates population’s fate at threshold density

The medium-transfer experiment established the existence of at least one secreted, survival factor. But it does not explain the density threshold involved in the collective growth. We reasoned that the density threshold is due to a “response threshold”: a concentration above which a cell’s growth rate exceeds its death rate (Fig. 1G – where the purple curve crosses the dashed line). Before experimentally establishing a response threshold, we mathematically tested this idea. We built a stochastic (single-cell) version of the deterministic, population-level model for collective growth with a size threshold (Fig. 1G). The stochastic model has just one free parameter and is a modification of another model that we previously built to explain collective growth of yeasts at high temperatures (*2*) (see STAR Methods). In this model, a cell randomly chooses to either replicate, or die, or remain alive without dividing (Fig. 3C). Each of these actions occurs with a probability that is determined by one survival factor that every living cell secretes at the same, constant rate. The model assumes that the concentration of the secreted factor is uniform throughout the liquid medium immediately after their secretion. The probability of a cell replicating nonlinearly increases as the concentration of a secreted factor increases (Fig 3D - blue). The probability dying is constant (unaffected by the secreted factor) (Fig. 3D - red). To verify these probability functions, we tracked many differentiating cells with a microscope for up to 4 days and for multiple initial population-densities. From the resulting time-lapse movies, we extracted the rates (probabilities) of cell replications and deaths as a function of the initial population-density (Figs. S7–S9). These measurements confirmed the probability functions in our model (Fig 3D) with which we ran stochastic simulations. The simulations recapitulated the random outcome that we saw in our experiments at the threshold density: two populations both start with this density, but one goes extinct whereas the other one grows (Fig 3E – green curves). Moreover, the simulations yielded a value of the threshold density that closely matched the value seen in our experiments (Fig. 3F).

### Communication occurring below a millimeter scale cannot account for the collective growth

The stochastic model recapitulated the data by assuming that all cells sensed the same concentration of the secreted, survival factor. But this simplification does not exclude another model in which the survival factor forms a concentration gradient. Indeed, since we did not shake or disturb the culture dishes during incubation, the secreted molecule should form some concentration gradient whose shape depends on its diffusion length. Perhaps the survival factor’s diffusion length is sufficiently large that the factor can essentially be uniformly mixed. But this requires a proof. Considering the colonies in our microscope’s field of view (1.40 mm x 0.99 mm), we can consider two kinds of cell-cell communication that occur through the diffusing survival factor: An “intra-colony communication” in which cells within a colony communicate with each other (Fig. 4A - orange) and a “local colony-to-colony communication” in which cells between colonies in the same field of view communicate with each other (Fig. 4A - purple).

**Figure 4.**
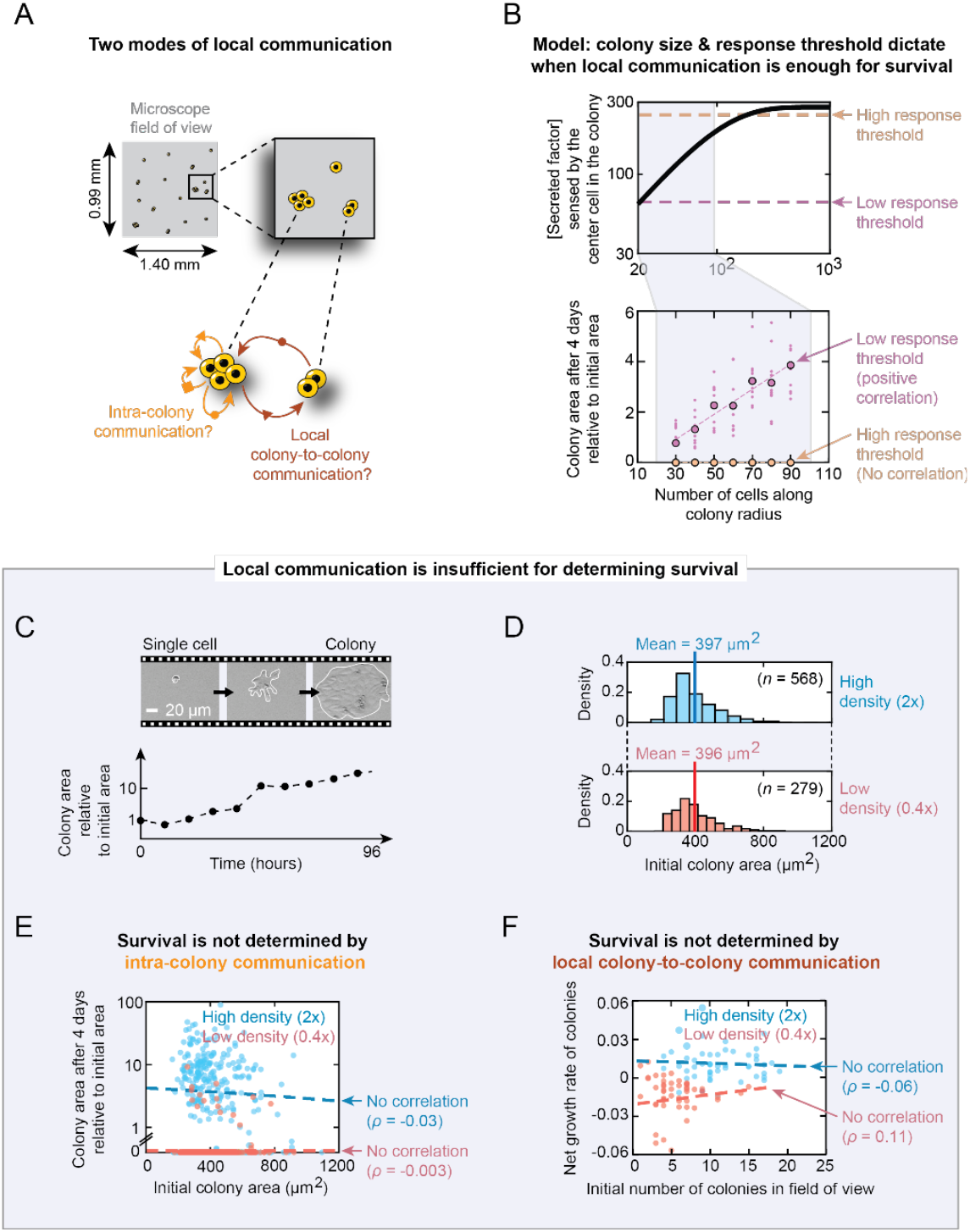
Communication occurring below a millimeter scale cannot account for the collective growth. (**A**) Two modes of local communication to exclude as possible mechanisms that control population’s survival-versus-extinction fate by using time-lapse microscopy. Each field of view in microscope is 1.40 mm x 0.99 mm. (**B**) Model-produced predictions for local communication to determine survival (see STAR Methods). To survive a colony must secrete enough concentration of a diffusible secreted factor to surpass the response threshold of the center cell in the colony. For that, the initial colony must be sufficiently large in size to yield a positive correlation between number of cells along the colony radius (colony size) and colony growth, or else there would be no correlation between the two parameters. Only a sufficiently low response threshold (indicated with purple arrow) yields a positive correlation between colony size and colony growth (shown here: Pearson correlation coefficient is 0.98), because a high response threshold (indicated with yellow arrow) results in no correlation (Pearson correlation coefficient is 0). (**C**) Example of 96-hour time-lapse movie that tracks growth of a microcolony (subregion of a field-of-view shown) (see also Fig. S10). Scale bar = 20 μm. (**D**) Two histograms (sample sizes indicated as *n*) of initial colony areas (in μm^2^) for a high-density population (2727 cells / cm^2^) that survives and expands toward carrying capacity (shown in blue) and a low-density population (818 cells / cm^2^) that goes extinct (shown in red). Data from multiple fields of view are shown for E14 cells differentiating in N2B27 while previously self-renewing in serum with LIF. Means and sample sizes (*n*) are indicated for each histogram. (**E**) Experimental confirmation of model prediction shown in (B). Each blue dot represents a microcolony from high-density population (2727 cells / cm^2^ initially) and each red dot represents a microcolony from low-density population (818 cells / cm^2^ initially). Same data as shown in (D). Area of each microcolony after 4 days of differentiation normalized by its initial area (vertical axis) versus its initial area (horizontal axis). Blue line: linear regression with *ρ* = −0.03. Red line: linear regression with *ρ* = −0.003. See Fig. S12 for all population densities. (**F**) Blue and red data points represent the exact same microcolonies as in (E) but now showing microcolony’s net growth rate (vertical axis) versus the initial number of microcolonies in the ~1-mm^2^ field-of-view that contains the colony (horizontal axis). Blue line: linear regression with *ρ* = −0.06. Red line: linear regression with *ρ* = 0.11. See Figs. S13 for all population densities. Also see Fig. S14 for experiments that show that the distance between one colony and its nearest-neighboring colony does not determine the colony’s survival and its growth rate.

To reveal the effects of intra-colony communication, we revised the stochastic model to account for concentration gradients of the survival factor. We considered a circular colony of a defined radius (i.e., defined colony area). We computed the maximum possible number of cells, in two dimensions, that fit in the circle. We then calculated the steady-state concentration of the survival factor that is sensed by the cell in the middle of the colony (“center cell”) due to every cell in the colony secreting the survival factor at the same, constant rate (see STAR Methods). For this, we solved the three-dimensional reaction-diffusion equation to obtain the steady-state diffusion profile created by a single cell and then summed it over all the cells inside the circular colony. The calculations showed that a larger colony area leads to a larger concentration of the secreted factor at the center cell due to more cells contributing the secreted factor (Fig. 4B - increasing part in top graph). But this increase stops and the concentration at the center cell plateaus as the colony radius increases above the survival factor’s diffusion length (Fig. 4B - flattening in the top graph). This is because the cells that are further than the diffusion length from the center cell cannot send the survival factor to the center cell (see STAR Methods). Hence, there is a maximum (upper-bound) concentration that a colony can create at its center. Having a colony that is any larger does not yield any higher concentration at the center cell. A geometric reasoning (symmetry argument) shows that the center cell senses the highest concentration of the survival factor in a circular colony (see STAR Methods). So, if the center cell senses a concentration that is less than the response threshold, then every cell in the colony also senses less than the response threshold and thus, according to the stochastic model (Fig. 3D), the entire colony would die.

We simulated the stochastic model again but now with above reasoning and considering cells to be in a colony. We restricted the range of initial colony radii (sizes) to small values (Fig. 4B – shaded region in the top graph) because we observed a restricted range of initial colony-sizes in our experiments (Fig. 2B – bottom graph). If the response threshold is sufficiently low, meaning that it is a concentration that can be created by the initial colony-areas achievable by our cell-seeding method, then the simulations showed that all sufficiently large colonies grew (Fig. 4B – bottom graph). On the other hand, if the response threshold is sufficiently high, meaning that our cell-seeding method cannot generate sufficiently large colony areas, then the simulations showed that every colony formed by our cell-seeding method became extinct (Fig. 4B – bottom graph: orange points). The simulations show that the colony area after several days does not correlate with initial colony-area if the range of initial colony-areas is too small to achieve the response threshold at the colony center. The prediction of this model is that if the diffusion length is macroscopic (millimeters), then a colony that is initially millimeters in radius should survive. At the end of our article, we experimentally confirm this prediction.

To experimentally probe intra-colony and local colony-to-colony communication, we used a microscope to continuously track and measure the area of each colony in multiple, millimeter-scale field-of-views (1.40 mm x 0.99 mm) over four days of differentiation and for multiple initial population-densities (Fig. 4C and Fig. S10). Here, we used E14 cells that differentiated in N2B27 without any inducers after the cells were self-renewing in the serum medium with LIF. Every field of view started with colonies that were, on average, hundreds of microns apart from one another (Fig. S11). The range of initial colony-areas was virtually identical for both high population density (2727 cells / cm^2^) and low population density (818 cells / cm^2^) (i.e., Fig. 4D shows nearly identical histograms). The range had an upper bound of ~1000 μm^2^, which corresponds to ~40 μm in diameter if we assume the colonies to be perfect circles. We found that a colony’s area after 4 days was uncorrelated with its initial area, for both high and low initial-density (Fig. 4E; see Fig. S12 for all other population densities). Specifically, a relatively small colony in a population of a low initial density would almost certainly go extinct (Fig. 4E - red points) but a colony of the same, small size in a population of a high initial density would almost certainly keep on growing (Fig. 4E - blue points). According to our model (Fig. 4B) and the range of initial colony-areas (Fig. 4D), the lack of correlation seen here means that the response threshold is higher than can be achieved by a colony whose area is below ~1000 μm^2^ (characteristic length of ~40 μm).

Since intra-colony communication is insufficient for determining a colony’s survival, each colony must be receiving survival factors from other colonies (i.e., there exists colony-to-colony communication). The microscope movies showed that a colony’s growth rate (how colony area changes over time) was independent of the number of colonies in a field of view, for both the high and low population-density (Fig. 4F and Fig. S13). Specifically, everyone would almost certainly go extinct in a field of view with ~10 colonies for a population of a low initial density (Fig. 4F - red points) but everyone would almost certainly grow in field of view with the same, ~10 colonies for a population of a high initial density (Fig. 4F - blue points). Moreover, whether a colony dies during the four days was also uncorrelated with the distance between a colony and its nearest-neighbor colony (Fig. S14). Hence, the threshold concentration must be too high to be generated by local colony-to-colony communication within the ~1 mm^2^ field of view. This means two things. First, there must be a nonlocal (beyond 1-mm scale) communication between cells through the diffusing survival factor. Second, to enable the nonlocal communication, the survival factor’s diffusion length must be larger than 1 mm. We devote the rest of our article to confirming these two claims and identifying the survival factor.

### Survival factor is stable for days and is light enough to enable nonlocal communication

Solving the reaction-diffusion equation reveals that a molecule’s diffusion length is 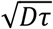, where *D* is the diffusion constant and *τ* is the half-life of the molecule (see STAR methods). *D* is proportional to the molecular weight. To place lower and upper bounds on the molecular weight, we flowed the differentiation medium (supernatant) of a 2-days-old high-density population (initially 5172 cells / cm^2^) through a membrane filter that captures all molecules that are larger (heavier) than the membrane’s “filter size” which is specified in kDa (*16*). The supernatant that passed through the membrane contains only the molecules that are smaller than the filter size (Fig. S15). Into this filtered supernatant, we transplanted a 2-days-old, low-density population (862 cells / cm^2^) (Fig. 5A – top). The filtered supernatant, containing all molecules that are smaller than the filter size, could only rescue the low-density population if and only if the filter size was 50 kDa or larger (Fig. 5B – top graph). This puts a lower bound on the molecular weight. In a second experiment, we filtered the supernatant, took all the molecules that were captured in the membrane, and then dissolved these into a fresh N2B27 medium. This medium, containing all molecules that are larger than the filter size, rescued the low-density population if and only if the filter size was 100 kDa or less (Fig. 5B - bottom graph). With control experiments, we determined that the filters make some errors in filtering molecules such that molecules that were lighter or heavier by ~50% of the filter-size value were passed through or captured by the filter membrane (Fig. S15) (see STAR Methods). Hence, the lowest possible molecular weight of survival factor(s) is ~25 kDa (i.e., 50% of the 50-kDa minimum mentioned above).

**Figure 5.**
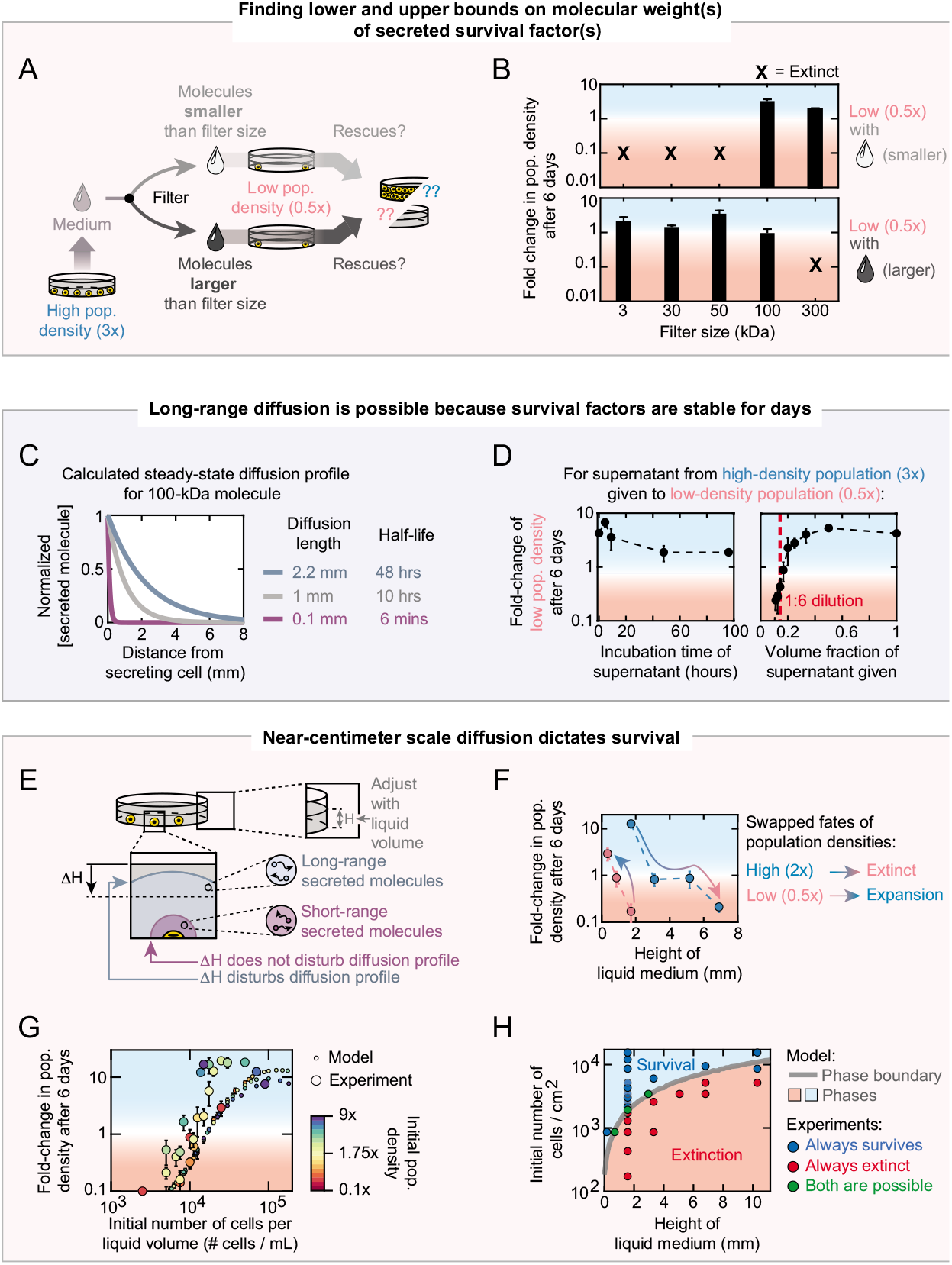
Survival factor is stable for days and is light enough to enable nonlocal communication. (**A**) Protocol for separating secreted molecules based on their molecular weights, through membrane-based filters (see STAR Methods). For all experiments we used 46C cells differentiating in N2B27+RA that were previously self-renewing in serum with LIF. Grey arrow: 2-days-old N2B27 medium is first taken from a high-density population (5172 cells / cm^2^ initially). We then run the medium through a membrane filter of a certain filter (pore) size, resulting in splitting of medium in two parts (splitting arrows): one part (light grey) contains all molecules that are smaller than the filter size (in kDa) and a low-density population (862 cells / cm^2^ initially) is then incubated in this medium to see if it expands or becomes extinct (light grey arrow). The other part of the filtered medium (dark grey) contains all molecules that are larger than the filter size (in kDa). Same procedure is carried out with this medium (dark grey arrow) as with the medium containing all the lighter molecules. See also Fig. S15 for experimental validation of filter’s error (~50%), hence allowing the lowest possible molecular weight to be ~25 kDa for secreted molecules. (**B**) Results of experiment described in (A). Fold-change in initial population density of the low-density population (vertical axis) that was incubated in the medium which contained either molecules smaller than the filter size (top graph) or larger than the filter size (bottom graph). Filter size is indicated on the horizontal axis. Cross (“X”) indicates that the population went extinct (i.e., fold change <0.1). Blue shade indicates population expansion and red shade indicates population extinction. Error bars are s.e.m.; *n* = 3. (**C**) Model-produced diffusion profile for secreted molecule with three different half-lives (*t*_1/2_) and diffusion lengths (*λ*) We used the (normalized) steady-state solution of the 3D reaction-diffusion equation and Stokes-Einstein relation to estimate diffusion constant of a ~100 kDa secreted molecule (nearly the maximum possible value from the filter experiments shown in (B)) (see STAR Methods for full details). (**D**) Experimental results showing that secreted molecules have a combined, effective half-life of at least 2 days. We used 46C cells differentiating in N2B27+RA that were previously self-renewing in serum with LIF. We transferred a 2-days-old high-density population’s (5172 cells / cm^2^) medium to an extinction-bound low-density population (862 cells / cm^2^), before which we either incubated the supernatant without cells at normal culture conditions (37°C, 5% CO_2_) for various periods of time (left graph) or diluted the supernatant with fresh N2B27 for various volume fraction (right graph). Data shows the resulting expansion of the low-density population. See caption of Fig. S26 for full details and explanations. Error bars are s.e.m.; *n* = 3. (**E**) Diffusion profile (volume filled by diffusing molecules) for molecule that diffuses far (blue region) and molecule that diffuses short range (purple region). Cell, in yellow, that secretes both molecules is adhered to the plate bottom. Δ*H* is change in liquid height. *H* is the total height of liquid medium that we tune in (H). (**F**) Results of experiment described in (E). Fold-change in population density after 6 days of differentiation, as a function of the height of the liquid medium (“*H*” in (G)). Red points are for low-density population (862 cells / cm^2^ initially) and blue points are for high-density population (3448 cells / cm^2^ initially). Data for 46C cells differentiating in N2B27+RA that were previously self-renewing in serum with LIF. Error bars s.e.m.; *n* = 3. (**G**) Data from new experiment performed after building the model (large circles; see Fig. 4) and model’s prediction (small circles). Plotted here is the fold change in population density after 6 days as a function of the initial population density which is now measured as # cells per mL of liquid volume (instead of # cells / cm^2^ that we used up to this point). Data from experiments and model predictions are from combining multiple volumes of liquid medium (in mL: 2, 5, 10, 18, 20, 30, 40 and 60) with initial # cells / cm^2^ (172, 431,862, 1293, 1724, 1931,2155, 2586, 3017, 3448, 4310, 5172, 6034, 8621,9483, 12069 and 15517 - indicated by the color bar). To obtain # cells / mL, we multiplied the # cells / area by the total area of the cell-culture plate (same for all conditions) and then divided the resulting value by the liquid volume. Data for 46C cells differentiating in N2B27+RA that were previously self-renewing in serum with LIF. Blue shade indicates population expansion and red shade indicates population extinction. See Figs. S16–17 for full data set. Error bars are s.e.m.; *n* is at least 3. (**H**) Model-produced phase diagram (grey curve and blue-red shadings) and experimental confirmation of the phase diagram (circles are from new experiments that were not used to build the model). Same experimental data as in Fig. 5G and Figs. S16–17 (*n* is at least 3; error bars are s.e.m.). Grey curve was constructed from the model by calculating, for each liquid medium height (volume), the threshold population density at which a population can either expand or become extinct (e.g., green curves in Fig. 3E). This, in turn, determines where the blue and red shades are in this plot. Circles are from experiments. Blue circles (“always survives”) are for initial conditions - defined by liquid medium’s height and initial number of cells / cm^2^ - for which all replicate populations expanded. Red circles (“always extinct”) are for initial conditions for which all replicate populations decreased in their density after 6 days (average of their fold change was below 0.6). Green circles (“both are possible”) are for initial conditions for which some replicate populations expanded after 6 days while some did not (average of their fold change was between 0.6 and 1).

Using the Stokes-Einstein relation, we could calculate the diffusion constant *D* directly from a molecular weight (see STAR Methods). For the argument outlined below, we assumed that the survival factor was ~100 kDa, which is nearly the maximum possible value from the filter experiments. Since the diffusion length *λ* is 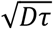, we assumed various values of *τ* (from minutes to days), calculated the corresponding *λ*, and then plugged this value into the solution to the reaction-diffusion equation to obtain the steady-state concentration profile for various *τ* (Fig. 5C). These calculations revealed that when the half-life is several days (e.g., 2 days), a cell can create a substantial concentration of the survival factor (at least ~25% of maximal possible concentration) several millimeters away (Fig. 5C). We experimentally confirmed that survival factors in the supernatant of a high-density population are stable enough to live for several days without diminishing survival-promoting effects. Specifically, we took the supernatant of a high-density population (5172 cells / cm^2^), incubated the unfiltered supernatant without any cells at 37 °C for various durations (up to four days), and then transplanted a low-density population into this aged supernatant (862 cells / cm^2^). Six days afterwards, the low-density population nearly doubled in density, which was about half of the ~4-fold growth in a ~4-hours-aged supernatant (Fig. 5D – left graph shows minor decrease over days). In another experiment, we diluted the fresh supernatant from the high-density population by various amounts into fresh N2B27 medium without ageing it, and then incubated the low-density population in the diluted supernatant. In the undiluted supernatant, the low-density population grew by ~4-fold whereas it grew by ~2-fold in a fresh medium that contained only 25% of the supernatant (Fig. 5D – plateau in the right graph). This result and the 4-day-old-supernatant experiment establish that the half-life of the survival factor (the indispensable survival factor if there is more than one specie) must be at least two days. Indeed, a shorter than two days of half-life would lead to the 4-days-old supernatant having less than 25% of the survival factor remaining. But we found that fresh medium with less than 25% of the supernatant causes the low-density population to grow by less than 2-fold. In fact, in media containing 1I6 or less of the supernatant, the low-density population went extinct (Fig. 5D – left of vertical red line in the right graph).

To verify the long-range diffusion in a different way, we considered the following: changing the height of the liquid-culture medium by millimeters would disturb the molecule’s concentration profile if the molecule can travel by millimeters in the first place (Fig. 5E - blue dome) but not if the molecule’s diffusion length is less than a millimeter (Fig. 5E - purple dome). For example, decreasing the medium height, which is ~2-mm above the cells (dish bottom) in our experiments so far, would lower the “ceiling” (top of liquid). If the secreted molecule can travel the ~2-mm distance, then it would now concentrate more at the bottom of the dish (on the cells) due to the decreased ceiling (Fig. 5E). Conversely, increasing the liquid height would let the molecule escape further away from the secreting cells and thereby concentrate less around the cells, given that the molecule can diffuse beyond the ~2-mm distance. Changing the liquid height would not change the initial population-density (number of cells per area). In accordance with this line of reasoning, a low-density population (862 cells / cm^2^) was rescued and grew towards the carrying capacity when we decreased the liquid height to ~0.3 mm whereas it became extinct if the liquid height was ~2-mm (Fig. 5F - pink points). Conversely, a high-density population (3448 cells / cm^2^) became extinct if the liquid height was ~7-mm but survived if the height was ~5 mm or 2 mm, with a faster growth in a 2-mm height than in a 5-mm height (Fig. 5F - blue points; also see Fig. S16 for percentages of cells that differentiated as a function of liquid height). Hence differentiating cells communicate through survival factor(s) that travel over a distance of at least ~5 to 7 mm (near centimeter).

With the diffusion length being at least several millimeters, we can effectively assume that the survival factor is well mixed. This justifies the stochastic model that assumed a uniform concentration for the survival factor (Fig. 3C). As a further proof of the survival factor being essentially well mixed, in another experiment, we widely varied both the initial population-density and the liquid-medium height. As a function of both, the fold-change in the population density over time closely agreed between the experiment and the stochastic model (Fig. 5G). Also, as a function of both the liquid height and the initial population-density, the stochastic model produced a phase diagram in which the boundary between population-level survival and extinction agreed well with the experimentally observed boundary. (Fig. 5H and Fig. S17). Given that a macroscopic range (near-centimeter scale) of communication dictates survival or death of the entire population in a switch-like manner (thus the sharp phase boundary in Fig. 5H), we will now refer the communication as yielding a “macroscopic quorum-sensing”.

### Differentiating cells macroscopically quorum sense by secreting and sensing FGF4

There may be more than one survival factor. To identify at least one of them, we first performed RNA-Seq on high-density (5172 cells / cm^2^), medium-density (1931 cells / cm^2^), and low-density populations (862 cells / cm^2^) (Figs. S18–S19). The medium-density population was near the threshold density and it neither expanded nor shrank to extinction over six days. For each density, we collected and lysed 46C cells that were differentiating for two days in N2B27 because this is when the growth-dynamics of a surviving population begins to be distinguishable from that of an extinction-bound population (Figs. 2C & 3B). In this RNA-Seq dataset, we identified 11 genes encoding secreted molecules whose weights were within the weight-range identified by the filters (Fig. 6A - listed along horizontal axis; also see Fig. S20–21). Of these, FGF4 and FGF5 were among the lightest molecules, with molecular weights of ~22 kDa and ~29 kDa respectively. We obtained recombinant versions of each of these 11 molecules (see STAR Methods). We then incubated a low-density population in 11 different N2B27 media, each containing just one of the 11 molecules. Of these, only FGF4 rescued the low-density population from extinction, leading to ~2-fold growth in six days (Fig. 6A - second bar). Notably, FGF5 alone did not rescue the population but the population approached extinction more slowly than any of the nine other molecules (Fig. 6A - third bar). Intriguingly, adding all 11 molecules into the N2B27 medium caused the most growth: the low-density population was rescued from extinction and grew by nearly 4-fold in six days, which is almost equal to the maximum possible growth seen in our medium-transfer experiments (see Fig. S22). This suggests that FGF4’s rescuing ability is enhanced by some of the ten other molecules. Incubating the low-density population in N2B27 with different amounts of FGF4 showed that the rescue from extinction requires FGF4 of at least ~2 ng/mL, which equals to ~0.13 nM (Fig. 6A - middle inset). The low-density population, incubated in N2B27 supplemented with 200 ngImL of FGF4, grew by ~10-fold after ten days and about ~80% of the cells had differentiated into the NE-lineage (Fig. 6A - right inset). These results establish that FGF4 alone is sufficient for rescuing the low-density population from extinction.

**Figure 6.**
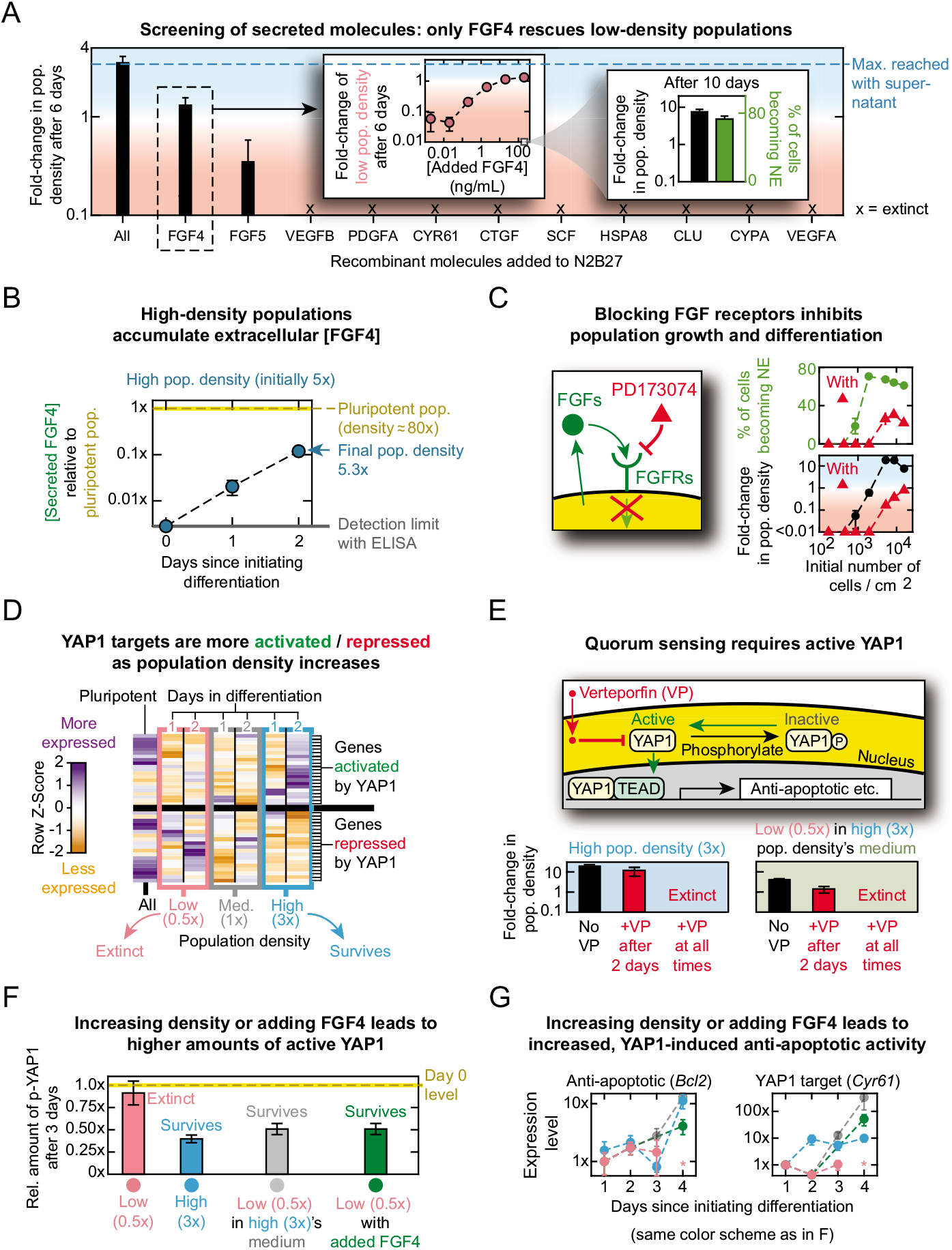
Cells collectively activate YAP1 and its anti-apoptotic targets by quorum sensing with FGF4. (**A**) Main graph: fold-change in initial population density, after 6 days in N2B27 supplemented with any of the 11 recombinant molecules (indicated on horizontal axis) or all of 11 combined (“All”) at day 0. All data are for initially low-density population (862 cells / cm^2^). Data for 46C cells differentiating in N2B27+RA that were previously self-renewing in serum with LIF. Also see STAR Methods and Fig. S22 for full details of concentrations added. Horizontal dashed line shows the maximum fold-change in density obtained in our study (nearly 4 fold), which occurs when the same low-density population is rescued by a high-density population’s (5172 cells / cm^2^ initially) medium. Middle inset: fold-change in initial population density after adding various concentrations of recombinant FGF4 (horizontal axis). Same procedure as described for main graph. Right inset: foldchange in initial population density (black bar) and percentage of cells entering NE lineage (Sox1-GFP expressing cells) (green bar), both measured 10 days after differentiation begins in presence of 200 ngImL FGF4 that we added at the start of differentiation. Error bars are s.e.m.; *n* = 3. (**B**) ELISA measurements of concentrations of extracellular FGF4 in the medium of a high-density population (8620 cells / cm^2^ initially) during unguided differentiation in N2B27 during 2 days (blue points) (previously self-renewing in serum with LIF) (see STAR Methods). Vertical axis shows FGF4 concentration relative to that of a ~80% confluent pluripotent population (denoted “1x” and marked with yellow horizontal line). 80% confluency equals i~8 x 10^6^ cells in 10-cm diameter dish. Lower detection limit of the ELISA assay is indicated (in grey). See Fig. S24 for ELISA standard curves. Error bars are s.e.m.; *n* = 3. (**C**) Cartoon shows PD173074, a well-characterized smallmolecule inhibitor of FGF receptors (*25,59*) (see STAR Methods). Fold-change in initial population density (bottom graph) and percentage of populations that enter NE lineage (top graph), both measured 6 days after differentiation began and as a function of initial population density. Data for 46C cells differentiating in N2B27+RA that were previously self-renewing in serum with LIF. Red data points in both graphs are for populations that were incubated with 2 μM (1056 ngImL) of PD173074 from the start of differentiation. PD173074 was dissolved in DMSO. Thus, as a control, black and green points are for populations without PD173074 but with the same amount of DMSO (volume per volume) as the populations represented by red data points. Blue shade indicates population expansion and red shade indicates population extinction. Error bars are s.e.m.; *n* = 3. (**D**) Heat map showing transcriptome-wide changes in unguided differentiation (N2B27) of 46C cells (previously self-renewing in serum with LIF) of low-density population (862 cells / cm^2^; enclosed in pink box), near-threshold (medium-density) population (1931 cells / cm^2^; enclosed in grey box), and high-density population (5172 cells / cm^2^; enclosed in blue box) (see STAR Methods). Leftmost column shows data for self-renewal (pluripotent) population before differentiation begins (labeled “All” since every population starts as this population before differentiation). Each column of differentiating population shows data for 1 day after (labeled “1”) or 2 days after (labeled “2”) starting differentiation. Each row shows a different gene, each of which are either activated (21 genes) or repressed (19 genes) by YAP1, either directly or indirectly. Fig. S19 lists all genes. Color represents row Z-score: a measure of by how much a gene’s expression level for a given condition deviates from that gene’s expression level averaged across all different conditions (i.e., different populations and days). Purple represents a positive row Z-score (more expressed than average). Orange represents a negative row Z-score (less expressed than average). Data based on 3 biological replicates. (**E**) Cartoon shows YAP1 which exists as either phosphorylated (labeled “P”) or dephosphorylated. Verteporfin (VP) inhibits active (dephosphorylated) YAP1 from entering the nucleus and regulating target gene expression. Fold-change in population density for high-density population (5172 cells / cm^2^ initially, in blue box) and low-density population that was rescued with medium of 2-days-old high-density population (862 cells / cm^2^ initially, in green box) after 6 days of differentiation towards NE-lineage. Data for 46C cells differentiating in N2B27+RA that were previously self-renewing in serum with LIF. Black bar: Verteporfin (VP) was always absent. Red bar in middle of each box: VP was added to medium after the first two days. Third column of each box shows absence of cells (extinction) when VP was present from the start of differentiation. Also see Fig. S28 for full data. Error bars are s.e.m.; *n* = 3. (**F**) ELISA measurements showing amounts of YAP1 protein phosphorylated at Ser397 (inactive YAP1) (see STAR Methods and also Fig. S29). Vertical axis shows the relative amount of inactive YAP1: the level of inactive YAP1 for a differentiating population divided by the amount of inactive YAP1 for a pluripotent population (in serum with LIF medium) that has the same cell numbers as the differentiating population at the time of lysing the cells for ELISA. Values are for 46C cells previously propagated in serum with LIF medium, 3 days after starting differentiation towards NE lineage with N2B27+RA medium. Pink: low-density population (862 cells / cm^2^ initially). Blue: high-density population (5172 cells / cm^2^ initially). Grey: low-density population rescued after two days by medium from a 2-day-old high-density population. Green: low-density population rescued by adding 200 ngImL FGF4 to its medium on day 0 (see (A)). Error bars are s.e.m.; *n* = 3. (**G**) *Bcl2* (anti-apoptotic gene) and *Cyr61* (YAP1-specific target) expression levels over time after initiating differentiation. Data obtained with RT-qPCR for 46C cells differentiating in N2B27+RA that were previously self-renewing in serum with LIF (see STAR Methods). Same color scheme as shown in (F). Also see Fig. S30 for other genes. On each day, we first normalized a population’s gene (*Bcl2* or *Cyr61*) expression level by that population’s *Gapdh* level (housekeeping gene). Afterwards, plotted on the vertical axis, we divided each population’s *Gapdh*-normalized gene (*Bcl2* and *Cyr61*) expression level on a given day by the *Gapdh*-normalized value for one-day-old low-density population (whose value is thus “1x” here). Error bars are s.e.m.; *n* = 3.

We found that differentiating cells indeed secrete FGF4. To determine this, we directly measured the extracellular concentration of FGF4 over time in the medium of a high-density population (8620 cells / cm^2^) that was differentiating in N2B27 without any inducers. Specifically, with ELISA, we detected the concentration of FGF4 increasing from zero to an appreciable amount during the first two days of differentiation in N2B27 (Fig. 6B - blue points). The final concentration of FGF4 that we detected after two days was ~10 times less than the FGF4 detected after two days in the self-renewal medium (FBS) of a highly confluent population (~16 times more cells than the differentiating population) (Fig. 6B - yellow line) (see STAR Methods for details of ELISA and rationale behind normalizing the concentration). Consistent with FGF4 being secreted and detected, we found with RT-qPCR that differentiating cells express *FGF4* and FGF receptors (*FGFR1-4*) during the first two days (Fig. S23).

We found that blocking the FGF receptors (FGFR1-4) causes differentiating cells to die. To determine this, we added a small molecule (PD173074) to the differentiation medium (N2B27 with RA) and then cultured populations of various starting densities (*25,59*). Inhibiting FGFRs continuously for six days drove every population towards extinction, including populations with densities higher than the threshold value (~1700 cells / cm^2^) (Fig. 6C - red points in bottom graph). A higher initial population-density meant a slower approach to extinction, suggesting that the more abundant FGF4 was competing for the FGFRs with the PD173074. The FGFR-blockage also drastically decreased the percentage of cells that entered the NE-lineage (expressed GFP) for all populations (Fig. 6C - top graph).

### FGF4 diffuses by at least several millimeters

We confirmed that FGF4’s half-life and molecular weight (diffusion constant) together yield diffusion length that is at least several millimeters. Specifically, we first determined that extracellular FGF4 in N2B27 without any cells does not measurably degrade for three days at 37°C (Fig. S24–25). Moreover, in another experiment, we took the medium of a high-density population and then incubated it at 37°C without any cells for four days. After four days, this medium still rescued a low-density population, with the low-density population growing by nearly the same amount as they do in fresh (unaged) supernatant from the high-density population (Fig. S26). Thus, the extracellular FGF4, contained in the medium of the high-density population, is stable for at least 3 days. With FGF4’s relatively small weight (~25 kDa), we used the Stokes-Einstein to compute the FGF4’s diffusion constant *D*. The diffusion length is 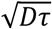, where τ is the half-life deduced above. Using the values obtained, we found that FGF4 diffuses over several millimeters (see STAR Methods). As a confirmation of the fact that diffusion is the primary means of spreading secreted factors in our culture conditions, we used a microscope to make a time-lapse movie in which a droplet of dye spreads in N2B27 without any cells at 37°C (Fig. S27). By analyzing how fast the boundary of the circular droplet spreads, we found that diffusion alone, without any other mechanism such as liquid convection, quantitatively accounts for the spreading. Thus, secreted molecules, including FGF4, would spread by diffusion rather than convection between cells.

### Cells collectively activate YAP1 by quorum sensing with FGF4

In the RNA-Seq on high-density (5172 cells / cm^2^), medium-density (1931 cells / cm^2^), and low-density populations (862 cells / cm^2^) (Figs. S18–S19), we found that the high-density population, compared to the low-density population, showed significant enrichments in genes that are involved in multicellular processes, cell-cell signaling, neurological processes, and cell adhesion (Fig. 6D - compare last column with second column; also see Fig. S18). More generally, compared to the low-density population, the high-density population had higher expression of genes that YAP1 directly or indirectly activates (e.g., *Cyr61* and *Amotl2*) and lower expression of genes that YAP1 directly or indirectly downregulates (e.g., *Angptl4* and *Tmem79*) (Fig. 6D and Fig. S19) (*60–67*). YAP1 is a master regulator of transcription for genes that control cell proliferation, apoptosis, and differentiation (*60,61,68,69*). It is part of the widely conserved Hippo-YAP signaling pathway. YAP1 is primarily known for being regulated by cell-cell contact-mediated signaling (*70*). But recent studies are beginning to show that YAP1 controls survival in a non-cell autonomous manner by cells that secrete diffusible signals such as Cyr61 (*61,71*).

The RNA-Seq results suggest that YAP1, as a master regulator, becomes more active in populations that begin with higher densities (Fig. 6D). YAP1 is inactive when it is phosphorylated. YAP1 is active when it is unphosphorylated. Verteporfin (VP) is a well-characterized inhibitor of YAP1 (*60,72*) (Fig. 6E - schematics). When we added 1 μM of VP in the serum (FBS) medium, self-renewing populations grew normally to the carrying capacity (Fig. S28). In N2B27 with RA, keeping 1 μM of VP from the start of differentiation (day 0) drove a high-density population (5172 cells/cm^2^) to extinction (Fig. 6E - “+VP at all times” in blue box). But, in N2B27 with RA, adding 1 μM of VP after two days did not hinder the high-density population’s growth: the population grew to nearly reach the carrying capacity in six days as usual (Fig. 6E - “+VP after 2 days” in blue box). This population behaved as if VP were never present. We obtained the same results when we added VP to N2B27 without RA (Fig. S28). A low-density population (862 cells/cm^2^), as shown before, survives and grows in the differentiation medium (N2B27) taken from a 2-days-old, high-density population (Fig. 6E - “No VP” in green box). Adding 1 μM of VP at the time of the medium transfer (start of differentiation) caused the low-density population to become extinct (Fig. 6E - “+VP at all times” in green box). In contrast, adding the VP two days after the medium transfer caused the low-density population to survive and grow, albeit more slowly compared to the situation in which VP was never present (Fig. 6E - “+VP after 2 days” in green box). These results suggest that the macroscopic quorum sensing requires activation of YAP1.

### Cells can only survive by collectively turning on YAP1 activity in first two days of differentiation

To confirm that YAP1 is crucial for the macroscopic quorum sensing, we used phospho-ELISA to measure the amount of phosphorylated (inactive) YAP1 in cells for various population densities. The phospho-ELISA specifically detected phosphorylation at Ser397, the primary phosphorylation site of YAP1 (*68,73*). For each initial population-density, we measured the amount of inactive YAP1 per cell, three days after starting differentiation. We also measured the amounts of inactive YAP1 per cell in populations that were self-renewing at various densities (Fig. S29). Note that we did not measure the total amount of YAP1. But we can still compare the levels of inactive YAP1 between the different populations because our RNA-Seq established that differentiating populations of different densities all expressed nearly the same level of *YAP1* (Fig. S29). In a low-density population (862 cells/cm^2^) that was headed for extinction, cells had nearly the same amount of inactive YAP1 as the self-renewing cells of the same density (Fig. 6F – first bar; and Fig. S29). In a high-density population (5172 cells/cm^2^), cells had about half as much inactive YAP1 (i.e., double the amount of active YAP1) compared to the cells in the extinction-bound, low-density population (Fig. 6F – second bar; and Fig. S29). In the low-density population that was rescued by a medium taken from a 2-days-old, high-density population, cells also had about half as much inactive YAP1 (i.e., double the amount of active YAP1) compared to the cells in the extinction-bound, low-density population (Fig. 6F – third bar; and Fig. S29). Crucially, when we rescued a low-density population by adding FGF4 to N2B27 with RA, cells had as much inactive YAP1 as the low-density population that we rescued with the medium from the high-density population (Fig. 6F - last bar; and Fig. S29). These results affirm that cells collectively secrete and sense FGF4 to activate YAP1, and that enough of the active YAP1 is necessary to avoid population extinction.

### Quorum-based activation of YAP1 increases anti-apoptotic processes

By measuring the expression levels of genes that YAP1 transcriptionally regulates, we found that *Bcl2*, an anti-apoptotic gene known to be activated by YAP1 (*60*), was upregulated in all populations that survived whereas it remained low in populations that went extinct. Specifically, with RT-qPCR, we measured *Bcl2* and other targets of YAP1 in four populations: a low-density population (862 cells/cm^2^: Fig. 6G - pink), a high-density population (5172 cells/cm^2^: Fig. 6G - blue), a low-density population cultured in the supernatant of a high-density population (Fig. 6G - grey), and a low-density population cultured with FGF4 (Fig. 6G - green). Of these, only the low-density population became extinct and maintained relatively low, basal level of *Bcl2* expression level that remained unchanged over days (Fig. 6G - pink in the left graph). The other three populations, between after 2-3 days, began to increase their *Bcl2* expression level by up to 10-fold (Fig. 6G - left graph). We also found that only the three, surviving populations had upregulated *Cyr61*, a well-known target of YAP1 (*61,71*), after 2-3 days of beginning differentiation (Fig. 6G - right graph). This supports the claim that the *Bcl2* upregulation is due to YAP1’s direct regulation (see Fig. S30 and STAR Methods for other YAP1-mediated genes that we studied). Finally, we also observed that the protein level of active caspase-3, a well-known apoptosis executioner (*60*), was ~2-fold higher in the low-density population than in the high-density and rescued populations (Fig. S31). Combined, these results establish that low-density (extinction-bound) populations have elevated apoptotic activities that are not countered by anti-apoptotic activities whereas growing (high-density or rescued) populations have elevated anti-apoptotic activities that are not countered by apoptotic activities.

### Macroscopic, millimeter-sized colony survives by itself due to intra-colony communication

Our cell-seeding method involved counting cells that were suspended in liquid (N2B27), and then spreading them out onto a 10-cm-diameter dish. Spreading a low concentration of cells (5000 cells/mL) led to a low-density population consisting of many individual colonies, each with an average area of ~400 μm^2^, that eventually went extinct (Fig. 7A). As a different seeding method, we seeded the same, low concentration of cells (5000 cells/mL) but confined within a small area of the 10-cm diameter dish at the center (Fig. 7B - schematic; see STAR Methods for the procedure). This seeding method produced a single region of ~28 mm^2^ that contained and started with many individual colonies that did not touch each other yet, which we confirmed with a microscope image taken 24-hours after the seeding (Fig. 7B - middle). This population survived whereas it would become extinct if it were spread out over the dish. Over the next five days, these colonies grew and merged to form a macroscopic colony, whose characteristic length (diameter) was 3-4 mm (Fig. 7B - right). This population was visible by eye as a single, three-dimensional colony in other biological replicates (Fig. 7C - for other biological replicates). This result matches our model’s prediction, mentioned earlier (Fig. 4B - top), which was that a colony that is sufficiently large survives purely due to intra-colony communication. Specifically, this discovery is another confirmation that the cells communicate over many millimeters: if they did not, then having a larger colony should not make a difference in the cells’ survival because many cells in the colony would be unable to communicate with each other because they are further apart than the diffusion length. If the macroscopic colony did not survive, then our models of cell-cell communication would be incorrect.

**Figure 7.**
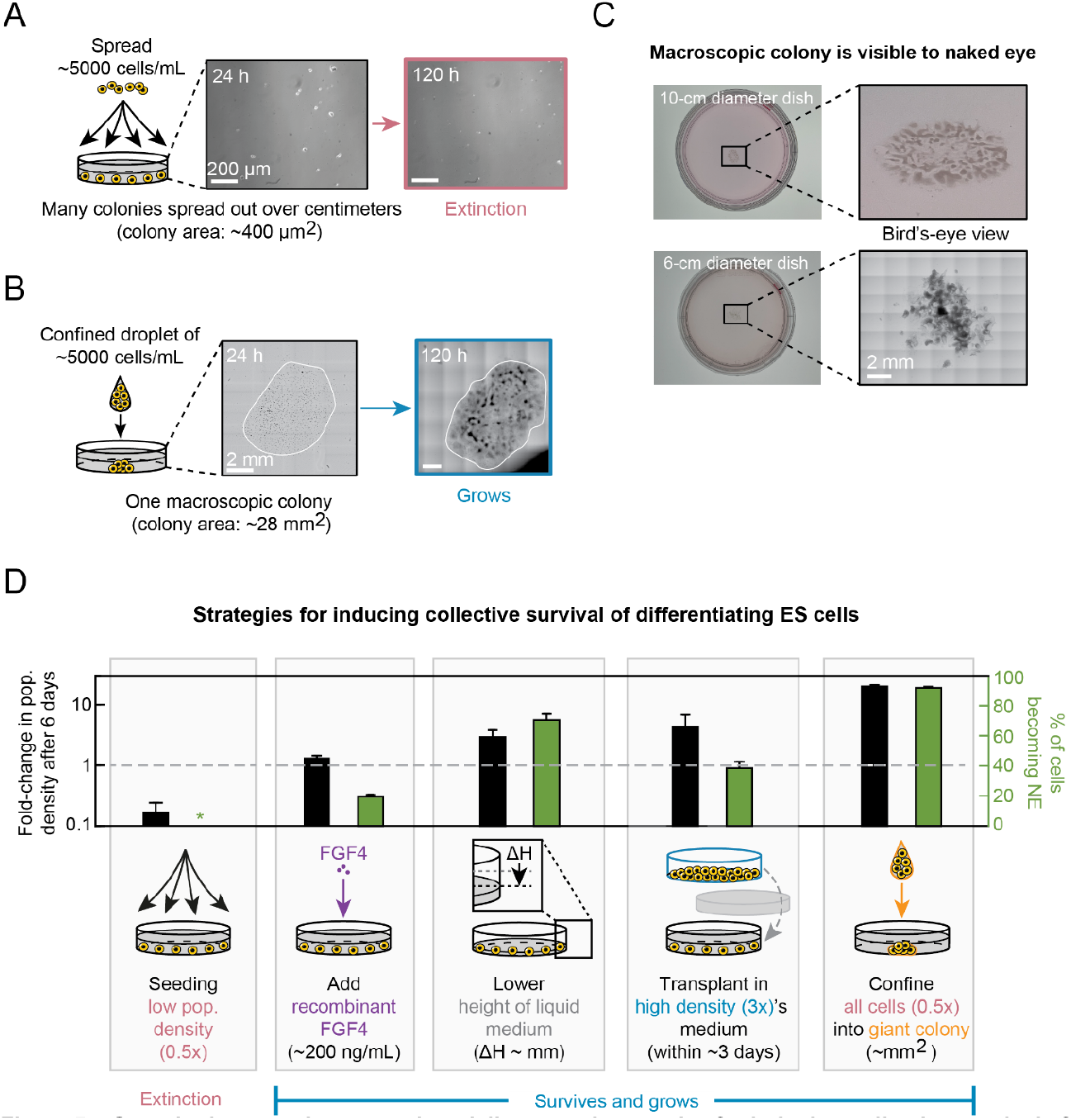
Quantitative experiments and modeling reveal strategies for inducing collective survival of differentiating ES cells. (**A**) Spreading a relatively small number of cells (~5000 cells / mL of N2B27) on a 10-cm diameter dish yields small colonies (size ~ 400 μm^2^) at the start of differentiation which eventually become extinct over time. Images are for 46C cells differentiating in N2B27+RA that were previously self-renewing in serum with LIF. Scale bars = 200 μm. Microscopy images are taken after 24 hours and 120 hours. (**B**) A same number of cells as in shown in (A) confined within a small area with a droplet on a 10-cm diameter dish yields macroscopic colony (size ~ 28 mm^2^) at the start of differentiation which eventually survives, expands and differentiates over time. Images are for 46C cells differentiating in N2B27+RA that were previously self-renewing in serum with LIF. Scale bar = 2 mm. Microscopy images are taken after 24 hours and 120 hours. In each of the two images, we stitched together multiple fields of view, with each field of view being 1.40 mm x 0.99 mm (same field size as in Fig. 2A). The stitching creates the checkered outline in both images. (**C**) Macroscopic colony, according to procedure described in (B), is visible to the naked eye after several days of differentiation (shown here is after 10 days of differentiation in N2B27+RA). Image taken after 10 days. In the greyscale image at the bottom, we stitched together multiple fields of view, with each field of view being 1.40 mm x 0.99 mm (same field size as in Fig. 2A). The stitching creates the checkered outline in this image. (**D**) Summary of strategies for inducing collective survival of differentiating ES cells. Spreading a relatively low number of cells (5000 cells / mL of N2B27) on a 10-cm diameter dish (shown here: 58 cm^2^ surface area (862 cells / cm^2^) and 10-mL N2B27) results in all cells becoming extinct during differentiation. Adding recombinant FGF4 (200 ngImL) to supernatant rescues spread cells from extinction (~1.3 fold-growth and ~19% Sox1-GFP positive). Lowering the height of liquid medium over millimetres rescues spread cells from extinction (~2.9 fold-growth and ~71% Sox1-GFP positive). Transplanting spread cells into a high-density population’s medium on day 2 rescues cells from extinction (~4.3 fold-growth and ~39% Sox1-GFP positive). Clustering a same number of cells, also used for spreading, into a macroscopic colony at the start of differentiation rescues cells from extinction (~20 fold-growth and ~92% Sox1-GFP positive). Data for 46C cells differentiating in N2B27+RA that were previously self-renewing in serum with LIF.

## DISCUSSION

In this article, we presented a systematic method to quantify the spatial range of communication between cells in cell cultures. It has been challenging to unambiguously determine the distance over which cells communicate through diffusible molecules even when the identity of the molecules involved have been known. Conventional methods, such as transferring of conditioned media and microfluidics, destroy the spatial information such as how far each autocrine factor travels when undisturbed. Aside from the travel distance of autocrine molecules, one must also determine whether a cell must sense at least a certain concentration of the molecule (response threshold) to respond to the autocrine signal. Our general model shows that the diffusion length of the autocrine molecule and the value of response threshold together determine the distance of communication (Fig. 4B).

By applying our method of determining communication distance to ES-cell cultures, we discovered that murine ES cells quorum sense at a centimeter scale during differentiation and that this quorum sensing determines whether the entire dish (population) of cells grows or becomes extinct. This determination is made in the first ~3 days of differentiation and is irreversible after this time. Crucially, FGF signaling is necessary and the FGF4 secreted by the differentiating cells, by itself, is sufficient for determining whether the population survives or becomes extinct during the first three days of differentiation.

Some cell-density dependent effects have been known for ES cells. Researchers have also known that ES cells secrete FGF4 and thus one may expect FGF4 to affect ES cells in a cell-density dependent manner. However, when one considers a molecule’s diffusion length and a response threshold for the molecule that potentially exists, then it is unclear whether any density-dependent effects, including those of FGF4, are due to a local communication between cells that are packed close to each other or due to cells that are millimeters-to-centimeters apart potentially stimulating one another. In other words, there remains an ambiguity. Namely, a cell might survive because it is helped by many, nearby cells. This can be the case in a high-density cell-culture in which the cells tend to be near each other and only local communication exists. Alternatively, a cell might survive because it is helped by enough distant cells that are millimeters-to-centimeters away. This can be the case in a high-density cell-culture in which distant cells can communicate but local communication is ineffective (e.g., long diffusion and high response-threshold, see Fig. 4B). Another possibility is a mixture of short- and long-range communication. In this case, one would need to determine how much of the density-dependent effect is due to a short-range communication as opposed to long-range communication. These ambiguities arise because all the above scenarios can lead to the same concentration of growth factors in the liquid medium that one collects from a cell-culture dish, as in the case of medium-transfer experiments. Our approach resolves such ambiguities by not disturbing cell-cell communication (paths of diffusing molecules) and analyzing the effect of communication on cell survival at every length-scale (from microns to centimeters).

By spreading out the cells in our cell-seeding method, the population densities that we achieved are lower than those in some standard cell-seeding protocols (e.g., ~15,000 cells / cm^2^ used in Mulas et al. (*53*)). All population densities, including the “high density” populations in our study, correspond to clonal densities (i.e., densities at which each colony arises from (near) single cell (Fig. 2B). Clonal densities are often used in standard differentiation protocols and for isolating individual cells from a heterogeneous population of cells. Our study shows that differentiating ES cells, at clonal densities, form colonies that communicate over vast distances. Our work shows that some colonies can still survive in the low-density populations that approach extinction (Fig. 2C - bottom: black curve gradually decreases). But the probability of a colony surviving is low (Fig. 2C - bottom: ~5% of cells remain alive after six days of differentiation). It may be possible that some cells are further along the differentiation process (e.g., towards NE lineage) than other cells in the population. Perhaps these faster differentiators are the ones that survive in these extinction-bound populations. Indeed, we found that the quorum sensing is only necessary in the early days (first 2-3 days) of differentiation (Fig. 3B). Hence it is plausible that the faster differentiating cells or cells that enter a NE- or ME-lineage faster than others survive the extinction fate (i.e., reason that the population size does not become exactly zero after six days). Future work may verify this hypothesis.

In making artificial tissues, blastoids, embryoid bodies, and other synthetic *in vitro* structures, one is interested in finding culturing methods that lead to high differentiation efficiencies (*33,37*). These culturing methods may involve aggregating cells rather than sparsely seeding them at clonal densities (e.g., in the case of making embryoid bodies or blastoids). As we demonstrated, a direct consequence of the macroscopic quorum sensing is that a murine ES-cell aggregate must be sufficiently large (i.e., comparable to the millimeters-long diffusion length) for it to survive during differentiation (Fig. 7C). The fact that a single ES-cell aggregate, whose characteristic length is several millimeters, survives on its own is consistent with our modeling which showed that intra-colony communication is sufficient for the aggregate (colony) to survive if the diffusion length is sufficiently large that the colony can reach such a size. While we do not exclude contact-mediated signaling between cells within an aggregate playing a role in the cells’ survival, it is unclear how, in an aggregate of cells, two cells that are touching each other would “feel” or determine the total size of the aggregate that they are in without diffusible signals. This result also strongly suggests that diffusion of FGF4 can indeed occur over millimeters in such a large aggregate, not just through a liquid medium. Moreover, if the aggregate had not survived, our model would be incorrect: cells can macroscopically quorum-sense if and only if single colonies (aggregate) of macroscopic sizes, but not of microscopic sizes, survive (Fig. 4B; and see STAR Methods).

Aside from concentrating cells into one macroscopic aggregate, our article reported several ways to ensure a collective survival of murine ES cells during differentiation into the NE- or ME-lineage. We also found the differentiation efficiency for each method (Fig. 7D). These methods include adding FGF4, decreasing the medium height by several millimeters, conditioning culture medium with medium from a high-density population, and concentrating cells into a millimeter-scale aggregate. Of these, we found that reducing the spatial dimension, by reducing the medium height or concentrating cells into a macroscopic aggregate, yielded the highest differentiation efficiencies. In fact, the macroscopic aggregates had >90% differentiation efficiency and exhibited the largest increase in population size among all four methods (Fig. 7D - last column). Our work lays the foundation for further studies of intracolony communication with diffusible factors in macroscopic aggregates such as blastoids and embryoid bodies. We expect that such studies, together with our systematic approach for determining spatial range of cell-cell communication, will lead to new tissue-engineering methods in synthetic biology. We also expect follow-up studies to yield further, quantitative understanding of how microscopic cells can bridge vast length-scales to perform a coherent, biological function as well as reveal elucidating physical principles that underlie collective growth of cells (*13,34,41,46,79–82*).

## AUTHOR CONTRIBUTIONS

H.D. and H.Y. conceived the project and designed the experiments. H.D. performed all the experiments and data analyses. P.vd.B. assisted with the experiments and performed the mathematical modeling with help from H.D. H.Y. provided overall guidance. H.D. and H.Y. wrote the manuscript with inputs from P.vd.B.

## ACKNOWLEDGEMENTS

We thank Thomas Fazzio, Sally Lowell, Amir Mitchell, and Erwin Frey for insightful discussions andIor comments on our manuscript; Marloes Arts for help with experiments; Brian Analikwu and William Verstraeten for help with analyzing the RNA-Seq data; and members of the Youk laboratory for discussions. We thank Austin Smith for sharing his 46C cell line, Valerie Kouskoff for sharing her Brachyury-eGFP cell line, and Angie Rizzino. H.Y. was supported by the European Research Council (ERC) Starting Grant (MultiCellSysBio, #677972), Netherlands Organisation for Scientific Research (NWO) Vidi Award (#680-47-544), CIFAR Azrieli Global Scholars Program, CIFAR Catalyst Grant, and EMBO Young Investigator Award.

## STAR METHODS

### RESOURCE AVAILABILITY

#### Lead contact

Further information and requests for resources and reagents should be directed to and will be fulfilled by the lead contact, Hyun Youk.

#### Materials availability

No newly generated materials are associated with this paper.

#### Data and code availability

RNA-Seq data in this work is available on NCBI’s Gene Expression Omnibus and are accessible through GEO Series accession number GSE157642. All data and related MATLAB codes supporting the conclusions of this study are deposited at Dryad and are publicly available under doi:10.5061Idryad.05qfttf3g. Any additional information is available from the lead contact upon reasonable request.

### EXPERIMENTAL MODEL AND SUBJECT DETAILS

We used three murine ES cell lines: E14Tg2a.IV (129IOla), 46C and Brachyury-eGFP. The 46C cell line (Sox1 promoter driving GFP) was previously described by Ying et al. (*51*) and was a kind gift from Austin Smith whose lab constructed it by targeting GFP to the endogenous Sox1 locus. Thus, the 46C cells had the Sox1 promoter controlling a GFP expression. The Brachyury-eGFP cell line was previously described by Fehling et al. (*57*) and Pearson et al. (*58*) and was a kind gift from Valery Kouskoff whose lab constructed it by targeting eGFP to the endogenous Brachyury (T) locus. Thus, this cell line had the Brachyury (T) promoter controlling an eGFP expression. To keep ES cells pluripotent, we routinely (every 2 days, with 1I10 dilution) passaged them in either a serum-based (FBS) or a serum-free (2i) pluripotency medium. The serum-based medium (denoted serum with LIF) consisted of high-glucose DMEM (Gibco) supplemented with 15% fetal bovine serum (Gibco, ES qualified), 1X MEM non-essential amino acids (Gibco), 1 mM sodium pyruvate (Gibco), 1X glutaMAX (Gibco), 0.1 mM 2-mercaptoethanol (Gibco), 1000 UImL penicillin-streptomycin (Gibco) and 1000 UImL Leukemia Inhibitory Factor (LIF, Polygene). The serum-free medium (denoted 2i+LIF) consisted of approximately half-and-half mixture of Neurobasal (Gibco) and DMEMIF12 (Gibco) and was supplemented with 1X MEM non-essential amino acids (Gibco), 1 mM sodium pyruvate (Gibco), 1X glutaMAX (Gibco), 1X N-2 (Gibco), 1X B-27 minus vitamin A (Gibco), 0.1 mM 2-mercaptoethanol (Gibco), 50 μgImL BSA (Sigma, fraction V), 1000 UImL penicillin-streptomycin (Gibco), 1000 UImL Leukemia Inhibitory Factor (LIF, Polygene), 3 μM CHIR99021 and 1 μM PD0325901. We performed sterile filtration all cell culture media with a 0.2-μm bottle top filter. Cells were maintained in the pluripotency medium on 10-cm diameter tissue-culture plates (Sarstedt, TC Dish 100 standard) that were coated with 0.1% gelatin in water (Sigma, from bovine skin Type B) at 37°C for at least 20 minutes prior to seeding cells.

### METHOD DETAILS

#### Description of mathematical model for autonomous and collective growth (related to Fig. 1)

For the model, we assumed that cell populations typically grow according to the logistic equation, which in simple terms implies exponential growth limited by the carrying capacity:

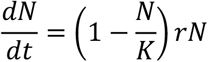

where *N* represents the number of cells in time, *K* the carrying capacity of the cells’ environment (determined by the availability of nutrients, space, etc), and *r* the net growth rate (= growth rate minus the death rate) of the cells. The net growth rate *r* can be either a constant (a positive number) or depend on a communication signal (e.g., a molecule secreted by the cells and hence depending on the number of secreting cells) that controls the net growth rate according to the Michaelis-Menten equation:

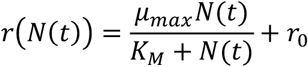

where *μ_max_* is the maximum net growth rate, *K_M_* (Michaelis constant) the number of cells at which the net growth rate equals half of the maximum net growth rate, and *r_0_* a constant (a positive or negative number). Here, we assumed that the net growth rate depends on the concentration of the communication signal which in turn is instantaneously set by the number of cells in time (see Figs. 1A, 1D and 1G). We used a custom MATLAB script to solve the differential equation for the following initial number of cells (initial population size) *N(0)*: 1; 17; 172; 1724; 5172; 20690. We computed the number of cells in time between 0 and 500 hours (see Figs. 1B, 1E and 1H) and the fold-change in the number of cells at an intermediate time-point (called “day 6”) (see Figs. 1C, 1F and 1I). For all three scenarios, we used the following (fixed) values: carrying capacity *K* = 172414, maximum net growth rate *μ_max_* = 0.0519 and Michaelis constant *K_M_* = 3500. For autonomous growth (scenario 1) (Figs. 1A-C), the net growth rate *r* is a constant equal to 0.0264. For collective growth without a size threshold (Figs. 1D-F) constant *r_0_* = 0.02 and for collective growth with a size threshold (Figs. 1G-I) constant *r_0_* = −0.0256. The carrying capacity *K* is indicated with a horizontal dashed line, and a “population expansion” (blue shade) and “population extinction” (red shade) refer to a fold-change that is above and below a foldchange = 1, respectively.

#### Description of stochastic model (related to Figs. 3 and 5)

Our model is a modification of another model that we previously built to explain collective growth of yeasts at high temperatures (*51*). In fact, the dynamics of differentiating ES cell populations (Fig. 2C) are strikingly similar to the population dynamics of yeast cells at high temperatures (see Fig. 2 in (*51*)) which also exhibits a survival-versus-extinction fate that depends on the initial population density (our yeast work examined how yeast populations grew as a function of temperature and the initial population density). In our model, differentiating ES cells constantly secrete a factor that degrades. This factor can be any molecule that promotes cell proliferation. We assume for simplicity that cells secrete such factor at a constant rate that in turn extracellularly accumulates. The probability of a cell replicating nonlinearly increases as the extracellular concentration of the secreted factor increases (Fig. 3D - blue curve). For simplicity, we assume that the probability of a cell dying is constant, as we experimentally verified (Fig. 3D).

Let *N_t_* be a stochastic variable that represents the number of alive cells at time *t*. We take time in discrete integer steps. At each time step, cells die with a constant probability *P_γ_* and replicate with probability *P*_μ_(*t*) which depends on the extracellular concentration of the secreted factor (denoted *M_t_*) at time *t*. The probability of replication *P*_μ_(*t*), as a function of *M_t_*, follows the Hill equation, with the Hill coefficient set to one for simplicity, and the maximum value μ. It is:

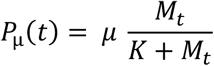

where *K* is a constant.

The secreted factor degrades with rate *d*. Later on, our experiments establish a long half-life of one particular secreted factor (namely FGF4) - it is at least 80 days (Fig. S25) - and a long, half-life for the mixture of all secreted molecules that determines the survival-versus-extinction fate (at least two days (Fig. S26)). At each time step *t*, the number of cells that replicate is sampled from a binomial distribution with *P*_μ_(*t*) being the probability of one cell replicating. The number of cells that die at time *t* is sampled from a binomial distribution with *P_γ_* being the probability of one cell dying. We assume that the molecule is well-mixed since our experiments established that the population effectively acts as a single entity that either dies or lives, due to the molecules that diffuse over several millimeters. Combining all the elements above, our model is completely described by the following set of stochastic equations:

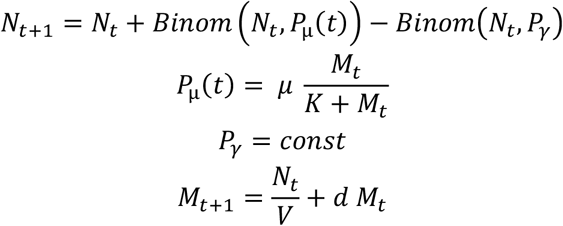

where *V* is the volume of liquid medium, and *N*_0_ is the initial number of cells. Since cells start without any extracellular secreted factor (since the differentiation medium initially has no molecules that determine the survival-versus-extinction fate), we must have *P*_μ_(0) = 0, which indeed follows from above equation. Note that we measure the secreted factor concentration (*M_t_* and *M_t+1_*) in units of secretion rate - the additional concentration of the secreted factor generated in one time step - as the secretion rate per cell is 1IV in the above equation.

The above set of equations has five parameters, four of which are directly fixed by our experimental data. Hence, we cannot freely tune the four parameters - *μ, P_γ_, V*, and *d* - and hence our model is highly constrained. We measured the maximum possible growth rate *μ* in two ways: (1) time-lapse microscopy in which we tracked the area of each colony over time for four days (Fig. S9 and Fig. S10); and (2) by counting the number of cells over time in a population (Fig. 2C, and Fig. S1 and Fig. S3). Both methods yielded a similar value: *μ* = 0.0519. To directly read off the value of the death rate from the experiments, we used the fact that a population of a very low initial density never grows and thus its rate of decline in cell number is *P_γ_* (green circle in Fig. S7 and Fig. S8) which we found to be: *P_γ_* = 0.0256. We determined *V* by directly measuring the volume of each liquid medium. Finally, we determined *d* by measuring the half-life of the mixture of all secreted molecules (Fig. S26) and the half-life of extracellular FGF4 alone (Fig. S25). We used the half-life of FGF4, which leads to d = 0.99 if we let each time step to represent 1 hour. With these constraints by the experiments, only one parameter, *K*, remains free for us to tune in the above set of equations. We chose the value for *K* such that the threshold density - at which the population can either survive or shrink towards extinction (Fig. 2C - middle row) - matches the experimentally determined threshold density. We thus let *K* = 485000.

We ran stochastic simulations that are based on the equations of our model, with the above-mentioned parameter values, for a wide range of initial density *N*_0_. For sufficiently low initial densities, we have *P*_μ_(*t*) < *P_γ_* for all times since cells are not able to accumulate enough of the secreted factor (cells continue to die and as this occurs, the cells’ efforts to accumulate the secreted factor is even further hindered). Thus, such a population goes extinct. For sufficiently high initial densities, we eventually have *P*_μ_(*t*) > *P_γ_* after some time. Here, *M_t_* becomes sufficiently high - it goes above the necessary threshold concentration at which the probability of replicating equals the death rate - which results in the population expanding towards the carrying capacity. For a particular set of initial population densities, we eventually have *P*_μ_(*t*) ≈ *P_γ_* for some stretch of time. This results in the population remaining nearly constant in cell numbers for some time. However, the population eventually gets pushed stochastically to either extinction or the carrying capacity (Fig. 3E). This population-level stochasticity occurs due to the stochastic actions by just a few cells in the population - whether or not a few cells stochastically divide while *P*_μ_(*t*) ≈ *P_γ_* and thus just slightly increases *P*_μ_(*t*) above *P_γ_*.

By running simulations for various initial densities at liquid volumes, we determined the phase-boundary that separates the survival phase and extinction phase (boundary curve in Fig. 5H). To get the boundary curve, we ran the simulation eight times for each pair of initial density and liquid volume. We then determined how many of these simulations, with the same initial condition, led to survival (and thus eventually reaching the carrying capacity) and how many of them led to a population extinction. In the extinction phase (red region in Fig. 5H), 100% of the simulations caused the population to become extinct. In the survival phase (blue region in Fig. 5H), 100% of the simulations caused the population to survive and eventually reach the carrying capacity. These determinations then allowed us to identify the boundary between the two phases - the phase boundary - which is shown as a curve in Fig. 5H. At this boundary, the probability of dying is equal to the probability of replicating, leading to equal numbers of cells replicating and dying, until stochastic fluctuations determine the population’s fate (survival or extinction). We found that this boundary nearly linearly increases with the liquid volume in the regime of media volume that we could experimentally access.

#### Description of mathematical model for colony size and response threshold (related to Fig. 4)

In this section we derive the criterion that determines when an intra-colony communication is sufficient for surviving. The steady-state concentration of a secreted molecule is

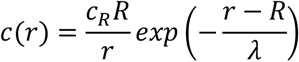

where 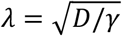 is the diffusion length and *c_R_* is the concentration on the cell surface (i.e., *r* = *R*). Let us consider *c* in units of *c_R_*. This normalized concentration is simply

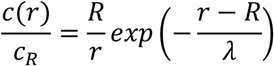

Consider a line of identical, spherical cells without any gap between them. Let us call the cell at the leftmost end of the line as a “receiver cell”. The cell to its immediate right is called the “1st cell”. The cell to the immediate right of this cell is “2nd cell” and so on. We want to calculate the normalized signal (secreted molecule) concentration created by all the other cells on the receiver cell. The distance *r_m_* between the receiver cell’s surface and the center of the *m*-th cell is

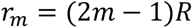

where *m* ≥ 1. Then the normalized concentration created by the *m*-th cell on the receiver cell’s surface is

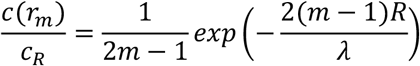

Now, imagine a 2-dimensional, circular colony formed by adherent cells. For simplicity, we can imagine that this colony is formed by a series of concentric circles, with spherical cells positioned circularly around each concentric circle. At the center of this circle is our receiver cell. It is surrounded by rings (concentric circles) of radii *R+r_1_, R+r_2_, R+r_3_*, and so on. Let *N_m_* be the total number of spherical cells that are circularly arranged on the ring of radius *r_m_*. We can estimate *N_m_* as the total number of cell diameters (2*R*) that can fit on the circumference of the ring:

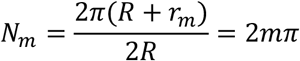

This is an overestimate of *N_m_* because this ignores parts of the ring that cannot be accessed by another cell due to the curvature of an occupying cell (i.e., it assumes that one can bend a diameter). One can quickly see that this is a good estimate by noting that *N_1_* ~ 6, which is also the number that one gets from hexagonally packing spheres (cells) in a ring whose center has the receiver cell. Note that hexagonal packing yields the maximum possible number of spheres packed in a given space, both in 2D and 3D. Then the total concentration *c_tot_* of signaling molecule created by all the other cells in a colony on the receiving cell at the colony center is

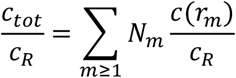

which simplifies to

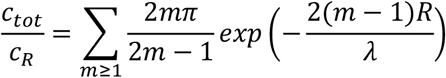

We can see that when *m* is large, the cells on the ring of radius *R* + *r_m_* contributes negligible concentration to the center cell because the summand in the above equation approaches zero as *m* increases. Specifically, we have

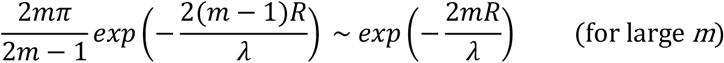

Hence, in our circular colony, cells that are much further away from the diffusion length *λ* contribute negligible concentration of signal to the colony center. This makes intuitive sense.

Note that above analysis also shows that dispersed cells (or other colonies) can contribute a non-negligible concentration on a receiving cell only if the diffusion length *λ* is sufficiently large. Only then, we have 2*mR/λ* < 1, meaning that the contributed concentration (i.e., 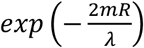) is non-negligible. This is consistent with our finding that the diffusion length is near-centimeter scale.

But total concentration *c_tot_* alone is insufficient to determine when a survival of a cell can be determined by an intra-colony communication with the same signaling molecule. To determine that, we need to compare the threshold concentration *c_thres_* to *c_tot_*. Basically, if *c_thresh_* is sufficiently low, then a small colony can generate enough concentration at the receiver cells so that *c_tot_* is above the threshold. In this case, according to our stochastic model (Fig. 3), we should observe a positive correlation between a colony’s initial size and its probability of surviving (equivalently, its growth rate since the net growth rate is proportional to the difference between a cell’s growth rate and its death rate). Having no correlation between the two quantities means that *c_thres_* is sufficiently high, meaning that it is higher than the *c_tot_* realized by any of the colonies. Since *c_tot_* is larger for larger colonies, a sufficiently high *c_thres_* means that no colony is sufficiently large to begin with. Our data is consistent with this latter scenario: we do not observe any correlation between colony size and its survivability (Fig. 4E and Fig. S12).

Note that if had a collective behavior *without* a threshold (Fig. 1D-F), above argument would not hold because in this case, a larger colony should grow faster. Having a threshold concentration binarizes the outcome so that a larger colony does not survive better than a smaller colony because both colonies can be too small to realize *c_tot_* that is larger than *c_thresh_*.

Finally, note that according to the above analysis, a sufficiently large enough colony should survive based solely on the intra-colony communication (i.e., even when the colony is by itself on a plate). Evidently, the colony sizes in our experiments are not sufficiently large (Fig. 4). The condition that very large colonies survive on their own accompanies the condition that the diffusion length must be at least as large as such colonies. Otherwise, due to the exponential decay term, 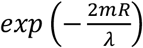, having a larger colony would make no difference to the concentration of the receiving cell. This reasoning is consistent with our finding that the diffusion length *λ* is near centimeter (every colony we observed is much smaller than a millimeter in diameter). We found that seeding a low number of cells as a single colony, by putting a single droplet of the cells in the middle of the dish, leads to a large colony of >1 mm^2^ (Fig. 7) that survives by itself. However, the same number of cells, if dispersed over the dish through our usual seeding method, causes that dish of cells to become extinct. This finding is consistent with our analysis above. There may be contact-mediated (mechanically mediated) signaling inside a colony that is also important for cell survival. Our analysis does not exclude this possibility.

#### Derivation of conditions required for millimeter-scale diffusion (related to Fig. 5)

Our experiments show that the cells secrete molecules that diffuse on the order of 1 mm. In this section, we use calculations to show that physics of diffusion also allows for biomolecules to diffuse on the order of 1 mm.

At first, a millimeter-scale diffusion may seem counterintuitive. This is because one often does not think about two cells, 1 mm apart, signaling to each other with diffusible molecules. Aside from the traveling axion potentials among neurons, one often does not think of millimeter scale communication among mammalian cells that are relatively immobile like ES cells. But physics allows for this for the proteins with the sizes that we experimentally found. Consider the general form of reaction-diffusion equation for the concentration *c* of a secreted molecule at position 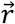 and time *t*:

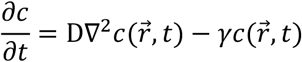

where *D* and γ are the molecule’s diffusion constant and degradation rate respectively. This equation lacks the molecule’s source (i.e., secreting cell) for simplicity and because we are interested in how far the molecule diffuses, which is independent of the secretion rate. A molecule’s diffusion length *L* is the characteristic (typical) distance that it travels before degrading. We can deduce *L* from dimensional analysis. Later on, we give the full, time-dependent solution to above equation which yields the same conclusion as the one we give in this section. The only parameters in the above equation are *D* and γ. *D* has the dimension, length^2^/time, and γ has the dimension, 1/time. Hence it follows that 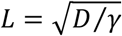, which is a well-known result. We can arrive at the same result through a more complicated way - by solving the reaction-diffusion equation above with a constantly secreting cell. But that is an overkill. We also have

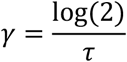

where τ is the molecule’s half-life. Next, as is common for biomolecules, we can estimate *D* by using the Stokes-Einstein relation for diffusing, spherical particles:

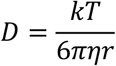

where *k* is the Boltzmann constant, *T* is temperature, η is the dynamic viscosity of the medium in which the molecule is diffusing, and *r* is the radius of the spherical particle that is diffusing. The ES cells grew at 37 °C and in an aqueous medium. Thus, for an order of magnitude estimate, we can use the dynamic viscosity of water at 40 °C: *η* = 0.653 × 10^−3^*N* · *s/m*^2^ (http://www.engineersedge.com). This value nearly stays the same at 30 °C and is thus insensitive to temperature for our purpose. Then, at 37 °C, we have

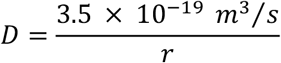

Before estimating the *r*, we can estimate the maximum diffusion constant *D_max_* for our molecules of interest. Our experiments with molecular filters showed that the secreted protein(s) must be in the neighborhood of ~100 kDa (specifically, between 50 to 300 kDa (Fig. 5A-B); or as low as ~25 kDa if we take the conservative estimate of ±50% error in the filter-pore sizes (Fig. S15)). Each amino acid has a mass of ~110 Da and end-to-end length of ~0.35 nm. Thus, our proteins of interest (of 100 kDa) would consist of ~1000 amino acids which, when joined stretched end-to-end, have a length of 350 nm (clearly a gross overestimate of the *r* for the secreted proteins). With this overestimate, we have *D_max_* = 1 *μm*^2^/*s*, which is the conservative, lower bound given for the diffusion constant of large biomolecular machines such as ribosomes. In fact, the conventional values assigned to the diffusion constant of typical proteins inside a cytoplasm fall in the range of 5 - 50 *μm*^2^/*s* (from BioNumbers (*74*)). Since the cytoplasm is a highly crowded environment that limits diffusion, we can expect a higher value for the protein of our interest that diffuses in the liquid medium.

We now estimate the *D*. The typical value of *r* for proteins is ~5 nm (from BioNumbers (*74*)). In contrast, *r* ~ 30 nm for eukaryotic ribosomes (from BioNumbers (*74*)), which are macromolecular complexes that are certainly heavier and larger than our proteins of interest as our experiments revealed. As a conservative estimate (i.e., to estimate a reasonable lower bound on the diffusion length), suppose that *r* ~ 20 nm for our proteins of interest, four times larger than the value of *r* for typical proteins (~5 nm). Then according to above equation, we have

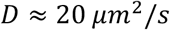

which is still reasonable in the ES cell’s relatively non-viscous liquid medium (recall that 5 - 50 *μm*^2^/*s* for a crowded cytoplasm). Note that

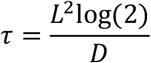

Hence, for a diffusion length *L* of 1 mm, we need τ ~ 12 hours. Note that this is a conservative, overestimate of the secreted molecule’s half-life since we underestimated the diffusion constant by assuming that the molecule is larger than half the radius of a ribosome (i.e., assumed *r* ~ 20 nm). A more realistic estimate such as, for example, *r* ~ 10 nm, would yield τ ~ 6 hours. A more typical value that one assigns to *r* for proteins is *r* ~ 5 nm, which would yield τ ~ 3 hours. In summary, both the conservative and more realistic estimates yield protein half-lives, which are required for a near millimeter-scale diffusion, that are well within the typically cited values of protein half-lives. Crucially, we measured the half-lives of the secreted molecules, including FGF4, in the supernatants of our cell cultures (Figs. S25–S26) and found that they are longer than the calculated half-lives here, meaning that the secreted molecules can, in fact, reach further than the 1-mm diffusion length we used in our calculation here.

Next, we use the steady-state solution to the three-dimensional reaction-diffusion equation to confirm our conclusions in the previous section, which were based on dimensional analysis. Assuming the ES cell to be a spherical cell of radius *R* that secretes a molecule at a constant rate *η*, equally in all directions, the three-dimensional reaction-diffusion equation is

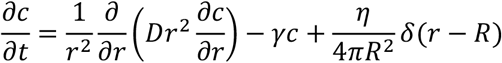

where *c* is the concentration, *r* is radial distance from the center of the ES cell, and *δ* is the Dirac delta function. As one can verify by substituting into the reaction-diffusion equation, its steady-state solution is

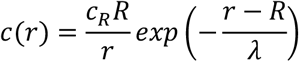

where 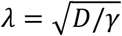 and

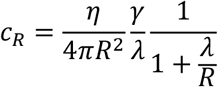

Note that *c_R_* is the steady-state concentration on the surface of the cell. The exponential term in the steady-state solution tells us that the characteristic length (i.e., diffusion length) is exactly 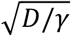, which exactly matches the diffusion length that we derived from dimensional analysis. Thus, the conclusions based on the dimensional analysis holds true. Note that this was not guaranteed since there could have been a small, numerical factor that scaled the *λ* down by several orders of magnitude.

#### Differentiation experiments

To initiate differentiation, we detached and collected ES cells from gelatinized dishes by incubating them with 1 mL of accutase (Gibco, StemPro Accutase Cell Dissociation Reagent) in 37°C for 5 minutes. After collecting the cells, we washed them twice with 1X PBS and then centrifuged them to remove any remaining accutase from the resulting cell pellet. We then resuspended the cell pellet in N2B27 which was prepared according to established protocols (*51,52*). Also see “Experimental model and subject details” for recipe of serum-free medium (N2B27), excluding LIF, CHIR99021 and PD0325901. We then counted the number of cells per mL in this resuspension, as described in the “Cell counting” section below. Afterwards, we calculated the volume of this resuspension required to achieve a desired number of cells per cm^2^ on a dish and then pipetted this volume into a tube containing 10 mL of N2B27 that was pre-warmed to 37 °C. We then transferred this onto a 10-cm diameter dish whose bottom was coated with a 0.1%-gelatin. We distributed the cells across the area of the dish by gently shaking the dish and then incubated the cells at 37°C with 5% CO_2_. Importantly, we did not disturb the plate for at least 6 hours after the plating to allow the cells to sediment and attach to the gelatinized plate bottom. We defined this moment to be the start of differentiation time-course. Cells were left for 2 days in N2B27 and then the spent medium was replaced with either fresh, pre-warmed N2B27 (for unguided differentiation), or N2B27 supplemented with 500 nM of Retinoic Acid (for Neural Ectodermal differentiation), or N2B27 supplemented with 3 μM of CHIR99021 (for Mesendodermal differentiation). We then left the dish for further incubation at 37°C with 5% CO_2_. Subsequently, we collected the cells from plates for counting (see “Cell counting” section below) and flow cytometry (see “Flow cytometry” section below). Importantly, we confirmed that large majority of ES cells (≥80%) differentiates into the Neural Ectoderm lineage without any inducer (such as RA; denoted “unguided differentiation”) and regardless of which medium (2i+LIF or serum with LIF) the ES cells were previously self-renewing in (Fig. S2).

#### Flow cytometry

We collected cells using accutase, washed them with 1X PBS, resuspended them in 1X PBS + 4% FBS and kept them on ice before doing instant measurements with a flow cytometer. We used a BD FACSCelesta system with a High-Throughput Sampler (HTS) and lasers with the following wavelengths: 405 nm (violet), 488 nm (blue) and 561 nm (yellow/green). We calibrated the FSC and SSC gates to detect only mouse ES cells (FSC-PMT = 231 V, SSC-PMT = 225 V, GFP-PMT = 476 V; as a control, flowing plain 1X PBS yielded no detected events). We measured the GFP fluorescence using the FIT-C channel. We analysed the flow-cytometry data using FlowJo and custom MATLAB script (MathWorks).

#### Time-lapse microscopy

We used a wide-field microscope (Nikon SMZ25) to track the growth of microcolonies as a function of the initial population density (E14Tg2A cell line). Cells were cultured on a 6-cm diameter tissue-culture dish (Sarstedt, TC Dish 60, Standard) coated with 0.1% gelatin (Sigma, from bovine skin Type B) in water, fed with 4-mL differentiation medium (N2B27) and incubated inside a temperature-, CO_2_- and humidity-controlled microscope chamber (Okolab) with steady conditions of 37°C with 5% CO_2_. We imaged microcolonies with the following initial population densities (in # cells / cm^2^): 455, 818, 1227, 1636, 2045, 2727, 3409 and 4091 cells per cm^2^ dish area. Previously, the cells were self-renewing in serum with LIF on 10-cm diameter dishes and routinely passaged (every 2 days). Importantly, cells were allowed for approximately 6 hours to settle and attach to the gelatinized bottom of the dish before the image acquisition. The microscope’s mono acquisition settings included a 1X microscope objective, 90.0X magnification, 28.5 (arbitrary units) of DIA intensity, 300-ms exposure time and 2.2X analog gain. Before image acquisition we picked a total of 17 fields-of-view that were evenly spread across the entire 6-cm diameter dish with each field-of-view being 1399.16 μm x 994.95 μm. For each initial population density we analyzed three biological replicates (three separate dishes), each in a different week and each consisting of 17 fields-of-view per dish. Images were acquired with 1-hour intervals in a total of 96hour duration (i.e., 4 days of imaging) during which cells were maintained in N2B27 without any inducer (unguided differentiation) without refreshing and thus disturbing cells. We analyzed the microscope images using custom MATLAB script (MathWorks). We found at most ~10 colonies per field-of-view for the lowest population densities and at most ~50 colonies per field-of-view for the highest population densities.

#### Cell counting

We detached cells from a cell-culture dish with accutase and then washed them twice with 1X PBS. We used a brightfield microscope (Motic AE31, 100X total magnification) and a hemocytometer (Marienfeld Buerker, #631-0921) to count alive cells by excluding dead cells with a Trypan Blue staining (i.e., dead cells appear blue). We counted the total number *N* of alive, non-stained cells in all 9 large squares – consistently excluding alive cells on two out of the four edges of each square. This way of counting enabled us to determine the total number of harvested, alive cells per unit of volume (mL) by back calculating, with the following formula: *N* x [dilution factor] x 10,000. In case we did not visibly detect any alive cells with this hemocytometer-based method of counting, we used a flow cytometer to estimate the cell counts (see “Flow cytometry” section and Fig. S1).

#### Medium-transfer experiments

We collected liquid medium of a high-density population (5172 cells/cm^2^), centrifuged it at 200 x g for 5 minutes to pellet and eliminate any remaining cells or other debris from the medium, and then transferred the medium to a low-density population (862 cells/cm^2^) after first removing the liquid medium of the low-density population. Specifically, we did this medium transfer in two ways. In one scenario (Fig. 3A – labelled as “1”), we collected the medium of a high-density population as described above on day X – the X here means X days after we initiated differentiation–and then incubated a low-density population in this medium to initiate its differentiation (i.e., the low-density population was in a self-renewing state before this medium transfer). In the second scenario (Fig. 3A – labelled as “2”), we collected the medium of a high-density population as described above on day X, and then incubated in it a low-density population that was differentiating for X days in its own medium. In this method, we measured the population density of the low-density population four days after the medium transfer (i.e., X+4 days) rather than on the same day for all values of X. This ensured that we could fairly compare the different low-density populations (i.e., same number of days (4) spent in the medium of the high-density population).

#### Medium-filtration experiments

We collected the liquid medium of a high-density population (5172 cells/cm^2^) 2 days after we initiated its differentiation, centrifuged the medium at 200 x g for 5 minutes to eliminate any cells and other debris from the medium, and then transferred the medium to a second centrifugal tube for ultrafiltration. The filter unit consisted of two compartments that were physically separated by a regenerated cellulose membrane which separates soluble molecules, depending on their molecular size and shape. Specifically, the membrane has pores that either pass or hold soluble molecules based on their molecular weight (in kDa) during a high-speed centrifugation. We used filter sizes of 3 kDa (Merck, Amicon Ultra-15 Centrifugal Filter Unit, UFC900324), 30 kDa (idem, UFC903024), 50 kDa (idem, UFC905024), 100 kDa (idem, UFC910024) and 300 kDa (Merck, Vivaspin 20 centrifugal concentrator, Z629472). We centrifuged the medium of the high-density population with spin times specified by the manufacturer. After the filtration, the centrifugal tube with the membrane filter contained two supernatants, each in separate compartments: one that contained all molecules that were larger than the filter size - this portion was much less than 1 mL and stayed on top of the membrane filter - and one that contained all molecules that were smaller than the filter size. We added the supernatant containing larger-than-filter-size molecules to a 10-mL N2B27 with 500 nM of retinoic acid (for Neural Ectodermal differentiation). In this mixed medium, we incubated a low-density population that had been differentiating for 2 days in its own N2B27. The results of this experiment are in the bottom graph of Fig. 5B. For a second experiment, we added 500 nM of retinoic acid to the ~9 mL of the supernatant that contained all the molecules that were smaller than the filter size. We then incubated in it a low-density population that had been differentiating for 2 days in its own N2B27. The results of this experiment are in the top graph of Fig. 5B. According to the manufacturer, in order to ensure that one captures proteins of a desired molecular weight, one needs to use a filter size that is at least two times smaller than the desired molecular weight. This sets a conservative safety/error margin that we took into account in the conclusions that we drew from the results in Fig. 5B, as explained in the main text. Finally, the few large molecules (> 3 kDa) that are ingredients of N2B27 were previously shown to have either no effect or a small growth-promoting effect on ES cells (i.e., they do not inhibit ES cell growth) (*16*). Additionally, we performed control experiments to confirm that the filters indeed did not catch any ingredients of the medium vital for cell growth (Fig. S15). Moreover, we tested the accuracy of the filters in catching a specific molecule of known size and effect by filtering a differentiation medium (N2B27) supplemented with recombinant LIF (1000 U/mL; 21.2 kDa according to manufacturer (PolyGene, ESLIF)) and then giving this filtrated medium to a low-density population (862 cells / cm^2^) otherwise bound to becoming extinct if not rescued. For this method, we measured the population density of the low-density population six days after giving either unfiltrated or filtrated N2B27+LIF (filtrated with a 50-kDa filter size). This ensured that we could fairly compare the different low-density populations.

#### RNA-Seq

We performed RNA-Seq on differentiating 46C populations that were previously self-renewing in serum with LIF of three different initial densities which were (in # cells/cm^2^): 862, 1931 and 5172. These populations were undergoing an unguided differentiation (i.e., in N2B27 without any inducers) and we examined their transcriptome 1 day after and 2 days after initiating their differentiations (Fig. 6D). We also performed RNA-Seq on a pluripotent 46C population, which would show the initial transcriptome of the three differentiating populations (Fig. 6D - first column). To perform RNA-Seq, we collected cells from each population and then centrifuged them using a pre-cooled centrifuge. We then extracted RNA (DNase-treated) from each cell pellet using the PureLink RNA Mini Kit (Ambion, Life Technologies) according to the manufacturer’s protocol. We next prepared the cDNA library with the 3’ mRNA-Seq library preparation kit (Quant-Seq, Lexogen) according to the manufacturer’s protocol. We measured the concentrations of each cDNA library before pooling using Quant-iT dsDNA Assay Kit (Invitrogen) and a Qubit Fluorometer (Invitrogen). We then loaded the cDNA library pool onto an Illumina MiSeq system using the MiSeq Reagent Kit v3 (Illumina). We analyzed the resulting RNA-seq data as previously described in Trapnell et al. (*75*). We performed the read alignment with TopHat (also using Bowtie, Samtools and BBDuk), read assembly with Cufflinks, and analyses of differential gene expression with Cuffdiff and CummeRbund. As a reference genome, we used the genome sequence of *Mus musculus* from UCSC (mm10). We performed enrichment analysis of genes based on their FPKM values (i.e., more than 2-fold expressed when two initial population densities are compared) by using GO-terms from PANTHER (*76*) and a custom MATLAB script (MathWorks). We visualized the results of pre-sorted, YAP1-related genes (*60–67*) as heat maps using CummeRbund and custom MATLAB scripts (MathWorks) which displayed the normalized expression value (row Z-score) for each gene and each condition.

#### RT-qPCR

We performed RT-qPCR on differentiating 46C populations that were previously self-renewing in serum with LIF of two different initial densities which were (in # cells/cm^2^): 862 and 5172. We observed them on each of four days of differentiation in N2B27 that was supplemented with 500 nM of Retinoic Acid (for Neural Ectodermal differentiation). We collected the cells and extracted their RNA with PureLink RNA Mini Kit (Ambion, Life Technologies) according to the manufacturer’s protocol. Then, we reverse transcribed (DNase-treated) RNA into cDNA using iScript Reverse Transcription Supermix for RT-qPCR (Bio-Rad). Next, we performed qPCR in 10-μL reactions with iTaq Universal SYBR Green Supermix (Bio-Rad) and 100 nM of forward and reverse primers. We verified the primer specificity and primer-dimer formation by using the melt curve analysis which showed one peak. See the list of primers that we used in Table S1. On each day, we normalized a population’s gene expression level by that population’s *GAPDH* (housekeeping gene) level. Afterwards, we compared each population’s *GAPDH*-normalized gene expression level for a given day to that of one-day-old low-density population (whose value is thus “1x” in Fig. 6G and Fig. S30). We performed all reactions in triplicates on a QuantStudio 5 Real-Time PCR System (Thermo Fisher).

#### Population-rescue experiments with recombinant proteins

We examined whether we could rescue a low-density population from extinction by adding one or a combination of 11 different autocrine-signaling molecules (all recombinant versions from mouse or human). We considered differentiating 46C cells that were previously self-renewing in serum with LIF in a low-density population (862 cells/cm^2^ initially). After 2 days of culturing in N2B27, we added 500 nM of Retinoic Acid, and one or combinations of the following recombinant proteins to the medium: 200 ng/mL of recombinant mouse FGF4 (R&D Systems, #7486-F4), 200 ng/mL of recombinant human FGF5 (R&D Systems, #237-F5), 100 ng/mL of recombinant mouse PDGFA (Novus, NBP1-43148), 100 ng/mL of recombinant mouse VEGFB 186 (Novus, #767-VE), 100 ng/mL of recombinant mouse VEGFA (Novus, #493-MV), 500 ng/mL of recombinant human CYR61/CCN1 (Novus, #4055-CR), 500 ng/mL of recombinant human CTGF/CCN2 (Novus, #9190-CC), 200 ng/mL of recombinant mouse CLU (Novus, #2747-HS), 500 ng/mL of recombinant human HSPA8/HSC70 (Novus, #NBP1-30278), 1000 ng/mL of recombinant human Cyclophilin A (PPIA) (Novus, #NBC1-18425), or 2000 ng/mL of mouse recombinant SCF (STEMCELL, #78064). After incubating in a medium containing one or a combination of these molecules for four days, we collected the cells for counting (see “Cell counting” section) and flow cytometry (see “Flow cytometry” section) to determine whether the population survived or not and its differentiation efficiency. Results of these experiments are in Fig. S22.

#### FGF4 ELISA

We measured concentrations of extracellular FGF4 in 10-mL liquid media (N2B27) as follows. We used Mouse FGF4 ELISA Kit (ELISAGenie / Westburg, MOES00755) and followed the manufacturer’s protocol. The assayed involved measuring the absorbance at 450 nm for various samples as a direct measure of the FGF4 concentration in the sample. We verified the absorbance signals are real and sufficiently high relative to the lower detection-limit of the ELISA kit by constructing a standard curve (Fig. S24). We measured the absorbances on a Synergy HTX Multi-Mode Reader (BioTek). We normalized the ELISA measurements of FGF4 concentration by comparing it to that of a highly confluent population of self-renewing ES cells. The latter is expected to have a high concentration of extracellular FGF4, based on previous studies’ finding that pluripotent ES cells highly express FGF4 (*25,77*). Normalizing all our ELISA measurements of FGF4 concentration by that of the pluripotent population also makes our result interpretable in the event that ELISA does not detect 100% of all FGF4s that are secreted (e.g., due to antibodies not binding to all their targets) (see Fig. S24 for validations of these justifications). With these justifications in mind, we used ELISA to measure the concentration of extracellular FGF4 in the medium of a high-density population (“5x” = 8620 cells / cm^2^) during the first two days of differentiation (Fig. 6B and Fig. S24). The 46C cells were differentiating for 2 days in N2B27 without any inducer and previously self-renewing in serum with LIF. We also used ELISA to measure the concentration of extracellular FGF4 in the medium of a highly confluent population of pluripotent cells (population density of ~80x) (Fig. 6B - yellow line). The differentiation medium did not initially have any detectable amounts of FGF4 (Fig. 6B - “day 0”). By using three different forms of FGF4 - one from the ELISA kit, FGF4 secreted by the cells in our experiments, and a recombinant form of FGF4 from a different company (R&D Systems, #7486-F4) - we found that indeed the ELISA kit does not detect all forms of mouse FGF4 but that it does detect the form secreted by the cells in our experiments, though less efficiently than it detects the recombinant FGF4 that accompanied the ELISA kit (Fig. S24).

#### Phospho-YAPI ELISA

We examined the endogenous levels of phosphorylated YAP1 protein in four different conditions (Fig. 6D and Fig. S29). We examined 46C cells that were differentiating for 3 days in N2B27 supplemented with 500 nM Retinoic Acid on day 2, and that were previously self-renewing in serum with LIF. For each measurement, we collected cells in 10-mL tubes, counted the total number of collected cells with the counting method described in “Cell counting” section, and then centrifuged them to form a pellet. We then lysed the cells with a lysis buffer (Cell Signaling Technology, #9803) and 1 mM of PMSF (Sigma-Alderich, P7626). We incubated the cell lysates with Phospho-YAP (Ser397) rabbit antibody and performed a sandwich-ELISA assay by using PathScan Phospho-YAP (Ser397) Sandwich ELISA Kit (Cell Signaling Technology, #57046). We then used a Synergy HTX Multi-Mode Reader (BioTek) to measure each sample’s absorbance at 450 nm. The absorbance is a direct measure of the abundance of phosphorylated YAP1. To compare the different absorbances, we constructed a standard curve by serially diluting a lysate of pluripotent cells (Fig. S29). We used the standard curve to report the levels of phosphorylated YAP1 in all differentiating populations.

#### Caspase-3 assay

We measured the protein-level activity of Caspase-3 (well-known apoptosis executioner inside cells) in E14 cells that were self-renewing in serum with LIF or were differentiating in N2B27 with 500 nM of Retinoic Acid (after self-renewing in serum with LIF). We examined three differentiating populations: high-density population (6896 cells/cm^2^ initially), a low-density population (517 cells/cm^2^ initially), and a low-density population that was rescued by the medium of the high-density population after two days of differentiating. We collected the cells of each of these populations and then performed a membrane-permeable DNA-dye-based assay that measures the amounts of active Caspase 3I7 in intact, alive cells (NucView 488 Caspase-3 Assay Kit for Live Cells). We followed all steps according to the manufacturer’s protocol. We used a flow cytometer to measure the amounts of active Caspase-3 in single cells. We normalized the Caspase-3 level per cell by the average Caspase-3 level of an ES cell (i.e., mean level per cell of the pluripotent population.) Results of these experiments are in Fig. S31.

#### Inhibiting FGF receptors

We examined the fold-changes in population densities after several days of inhibiting FGF receptors (FGFRs) with a small-molecule inhibitor, PD173074 (PD17, Tocris, #3044) (previous studies characterized this inhibitor: see (*25,59*)). We used 46C cells that were differentiating in N2B27+RA (RA added on day 2) and that were previously self-renewing in serum with LIF. We examined the following initial population densities (in # cells / cm^2^): 172, 431, 862, 1931, 5172, 8620 and 15517. To inhibit the FGFRs, we added 2 μL of 10-mM PD173074 (PD17) to a 10-mL N2B27 medium. We dissolved the stock of PD17 in DMSO to a final concentration of 2 μM (1056 ng/mL). After 6 days, we measured each population’s density (see “Cell counting” section) and differentiation efficiency (see “Flow cytometry” section). As a control, we examined the effect of adding 2 μL of DMSO to cell-culture medium without any PD17. This ensured that our results were not due to any side effects of having DMSO that was carried over with the PD17 that we added to each medium. Results of these experiments are in Fig. 6C.

#### Verteporfin experiments

We examined the fold-changes in population densities after several days of incubation verteporfin (R&D Systems, #5305), which prevents active YAP1 from entering the nucleus to control expression of its myriad target genes. We used 46C cells that were differentiating in N2B27 with and without RA (RA added on day 2) and that were previously self-renewing in either serum with LIF or 2i+LIF. We supplemented the differentiation medium with 1 μM of verteporfin that was dissolved in DMSO (based on LeBlanc et al., (*60*)). After 6 days, we measured each population’s density (see “Cell counting” section) and differentiation efficiency (see “Flow cytometry” section). As a control, we examined the effect of adding only DMSO to cell-culture medium without any verteporfin. This ensured that our results were not due to any side effects of having DMSO that was carried over with the verteporfin that we added to each medium. Results of these experiments are in Fig. 6E and Fig. S28.

#### Procedure for seeding a macroscopic colony

Our standard cell-seeding method involves spreading a desired number of cells across the gelatin-coated surface of centimeter-sized dish. Unlike this method, we clustered a relative low number of cells (~5000 46C cells per mL of N2B27) by injecting a few-microliter droplet of N2B27 containing the amount of cells at the center of a centimeter-sized (e.g., 10-cm or 6-cm) dish that was coated with 0.1% gelatin and contained the same volume of N2B27 as in the case of the standard cell-seeding method (e.g., 10mL for 10-cm diameter dish and 4-mL for 6-cm diameter dish). The cells that were initially confined by a droplet were then allowed to sediment and attach to the gelatinized dish bottom, which resulted in ~mm^2^ area occupied by microcolonies. We observed localized, individual microcolonies that were not touching each other within a small area (~28 mm^2^) with a brightfield microscope 24-hours after clustering of cells (Fig. 7B) as opposed to sparsely spread microcolonies (each ~400 μm^2^) when applying the standard seeding method (Fig. 7A).

### QUANTIFICATION AND STATISTICAL ANALYSIS

We generated all graphs and illustrations with Adobe Illustrator and with custom scripts in MATLAB, ImageJ, RStudio and FlowJo. Unless stated otherwise, we reported all experiments as means of biological replicates (obtained on different days) and the error bars are standard error of the means (s.e.m.). The number of biological replicates is denoted as *n* and at all times is at least 3. We included all such statistical details of experiments in the figure legends, main text or supplemental information.

## Supplemental Information

**Figure S1.**
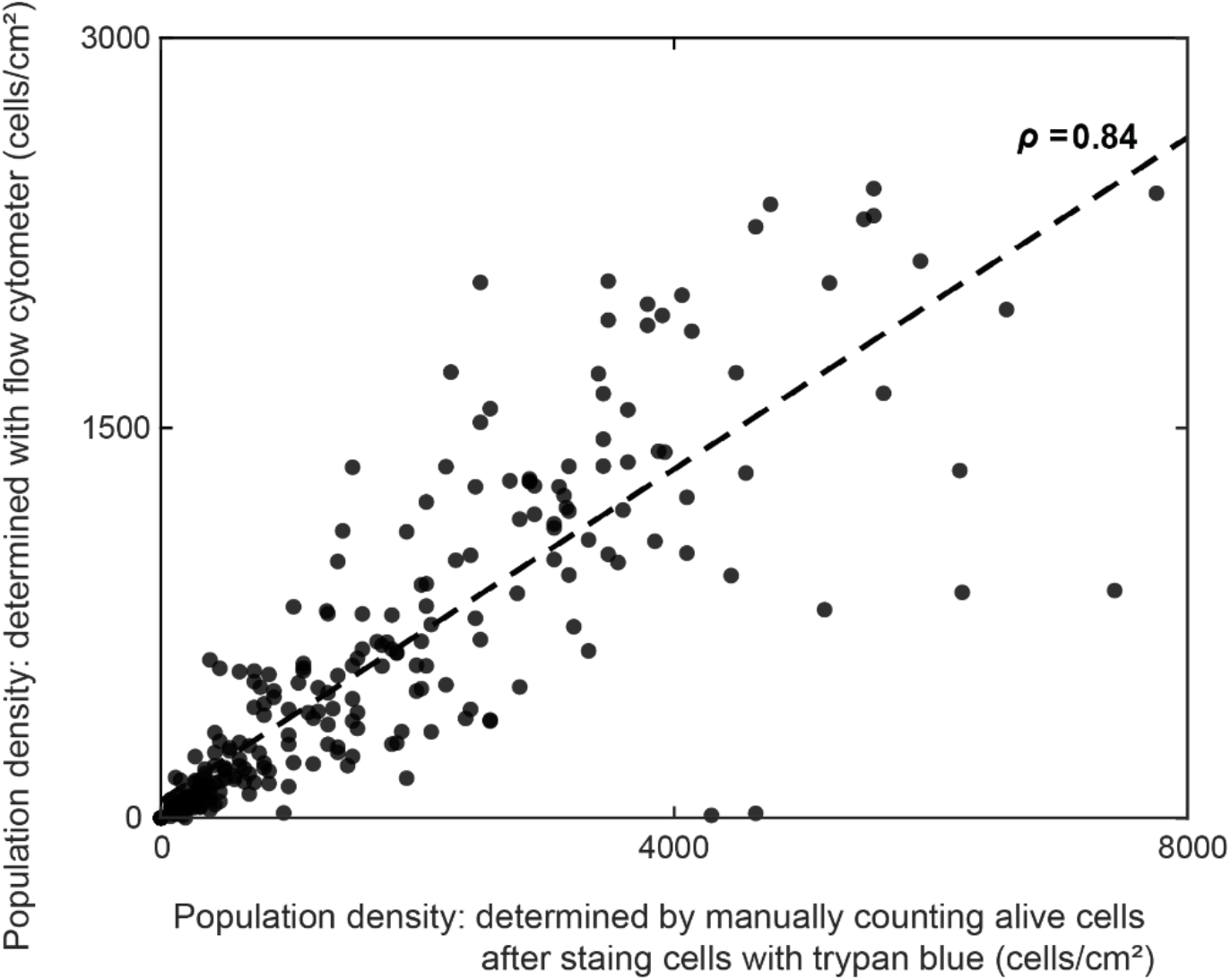
Two different methods of counting cells yield population densities that are directly proportional to each other (i.e., offset by a constant factor of order one) and in the same order of magnitude, thus affirming both methods for determining population densities (related to Fig. 2). Each data point represents a single population whose density (# of cells / cm^2^) was determined by two independent methods. Each axis represents a different method. As one method, we used a standard hemocytometer to count individual cells after subjecting the cells to the dye, trypan blue. Trypan blue penetrated only dead cells within the population. We counted the unstained (non-blue) cells with a hemocytometer to determine the resulting population density (# of cells / cm^2^) of alive cells on a cell-culture dish. As another method to determine the population density, we used a flow cytometer to count the number of cells (events) that belonged to a specified FSC-SSC gate. We set the FSC-SSC gate so that it captured alive cells while excluding dead cells. These two methods yielded cell counts - and thus the corresponding population densities - that were directly proportional to one another, as indicated by a high Pearson linear correlation coefficient (black line; *ρ* = 0.84). The proportionality factor is on the order of one, meaning that, for the same population, the two methods yield numbers that are in the same order of magnitude as the population density. Given this result, throughout our study, we primarily counted cells manually (i.e., with trypan blue) and used the flow cytometer when this was not possible. Specifically, we used the flow cytometer to count populations that had very few cells such as those near extinction (i.e., populations whose fold-change in density was near or below 0.1 after some days).

**Figure S2.**
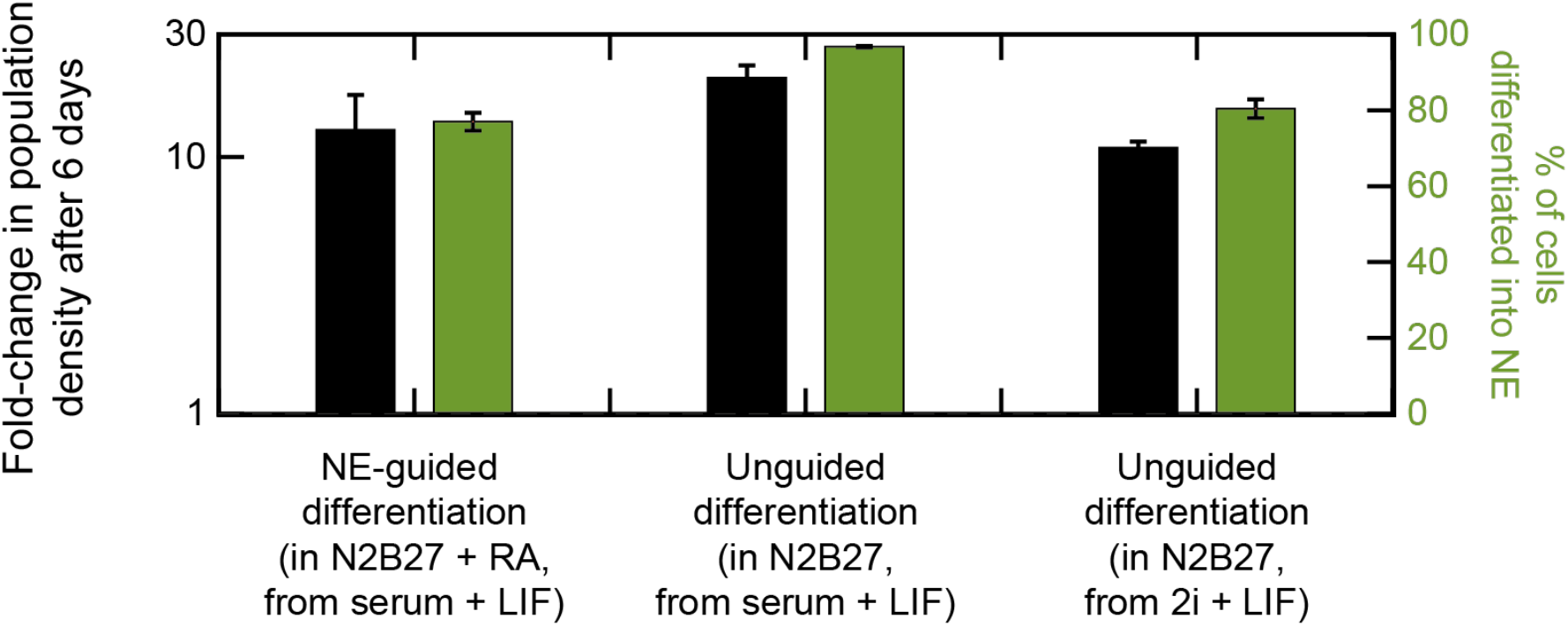
Large majority of ES cells (≥80%) differentiates into the Neural Ectoderm lineage without any inducer (such as RA) and regardless of which medium (2i or serum) the ES cells were previously self-renewing in (related to Fig. 2). **(A)** Data shown 46C cell line. The 46C cells have a Sox1 promoter controlling their GFP expression (Sox1 is a marker of the Neural Ectoderm (NE) lineage). Black bars represent fold-change in initial population density after 6 days of differentiation. Green bars represent percentage of cells becoming Sox1-GFP positive. For each differentiation, we took 46C cells that were kept pluripotent with serum+LIF or with 2i+LIF (serum-free) medium. These starting conditions are indicated in the captions below the black and green bars (e.g., “from serum + LIF”). The cells were then differentiated in N2B27 (without any serum), in the presence or absence of the inducer Retinoic Acid (RA) (added on day 2). These differentiation conditions are indicated in the captions below the black and green bars (e.g., “in N2B27 + RA”). As seen here, regardless of the use of inducer or which medium (2i or serum) the cells were previously self-renewing in, all populations expand significantly (≥10 fold) and differentiate substantially (≥80% becoming Sox1-GFP positive) into the NE lineage in accordance with literature/established protocols (*51,52*). *n* = 3; error bars are s.e.m.

**Figure S3.**
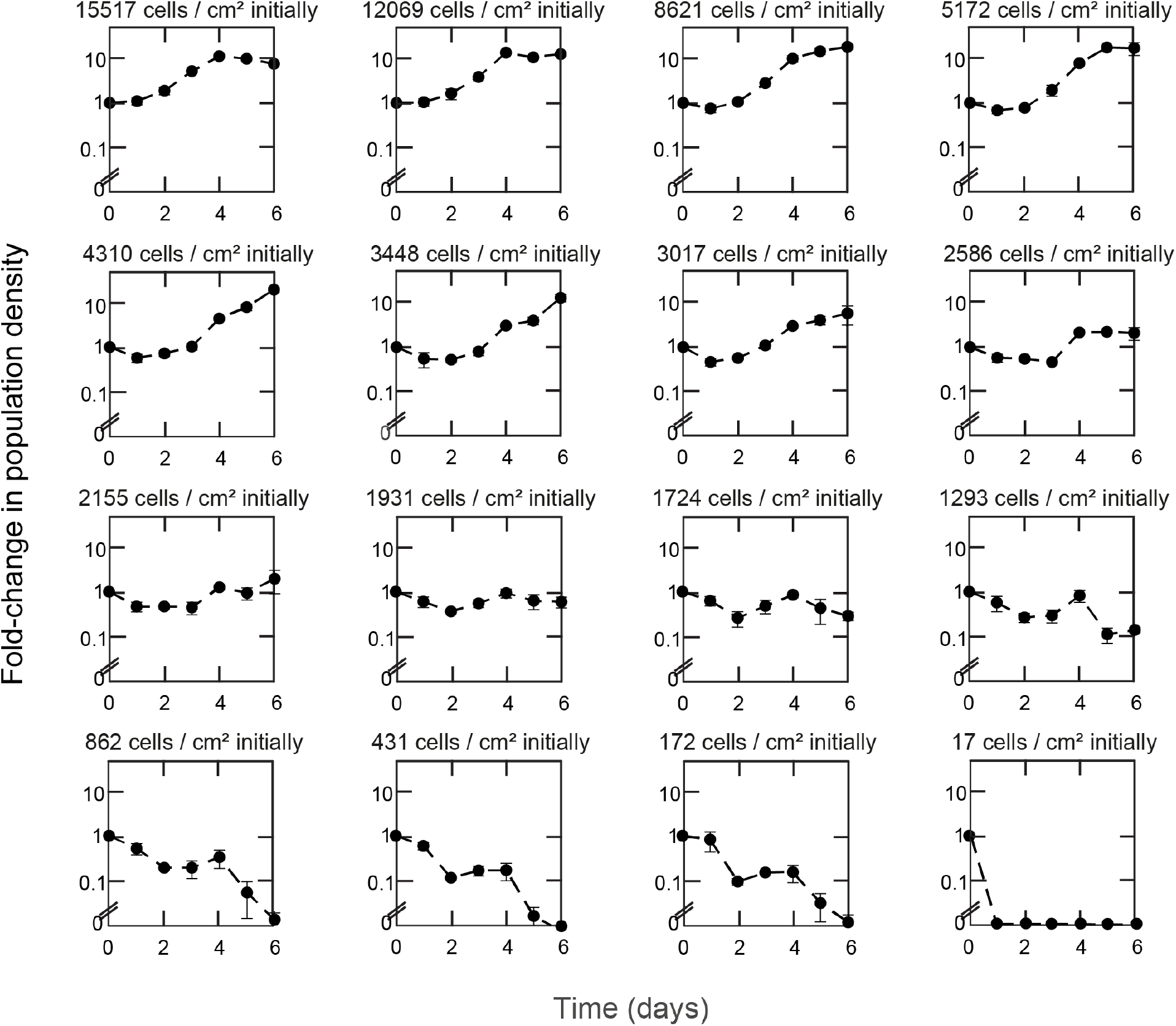
Population growth during differentiation towards the Neural Ectoderm lineage for a wide range of starting population densities (related to Fig. 2). Data for 46C cells (which have Sox1 promoter driving GFP expression) differentiating towards NE lineage in N2B27+ Retinoic Acid (RA) that were previously self-renewing in serum+LIF (see STAR Methods). We triggered pluripotency exit to begin each time course shown here (protocol in Fig. 2A). Each box shows the population dynamics for a different starting population-density (indicated above each box). Each box shows the fold-change in population density (vertical axis) as a function of the time passed since triggering differentiation (horizontal axis). *n* is at least 3 for each data point. Error bars are s.e.m.

**Figure S4.**
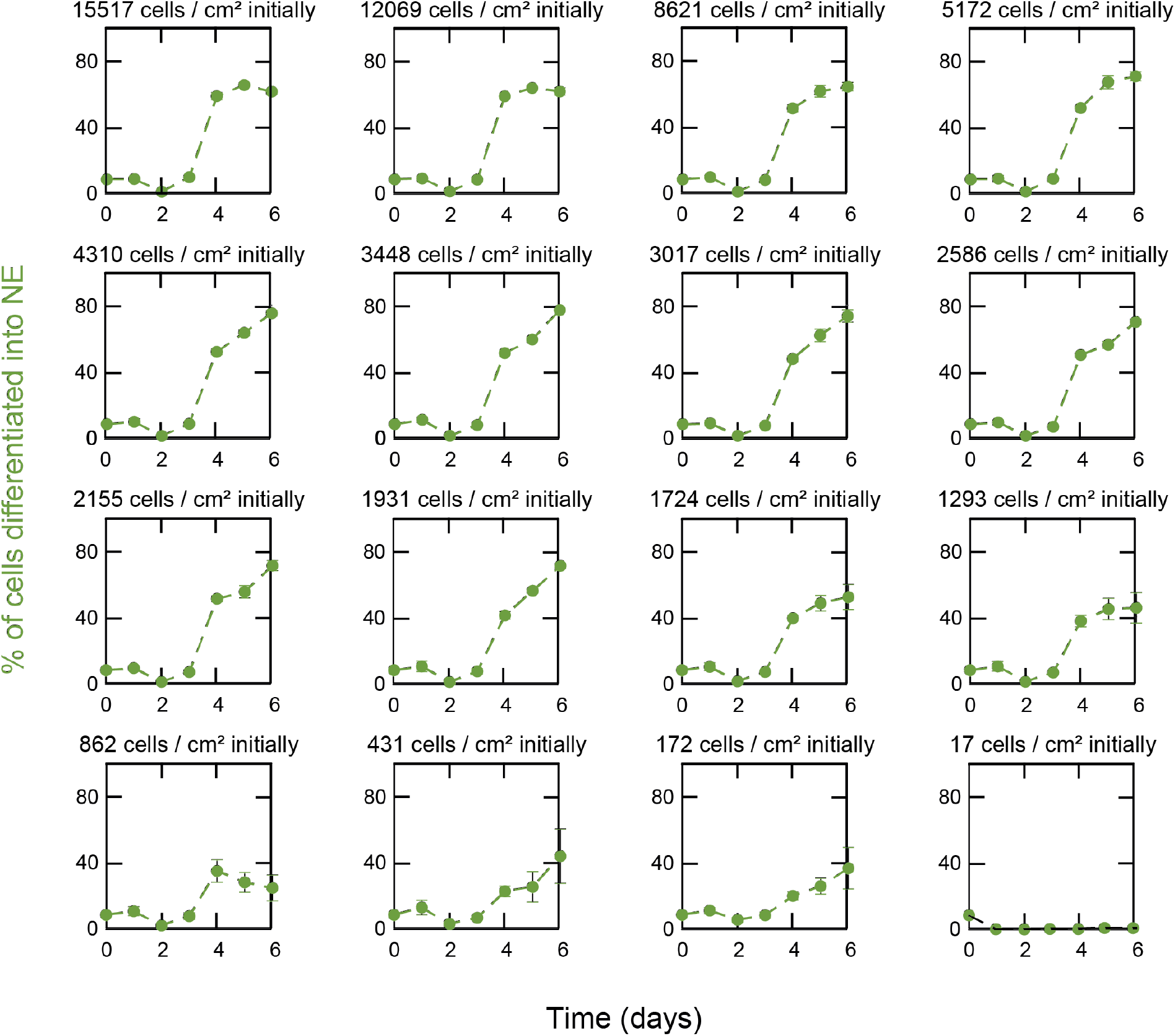
Populations with higher initial densities achieve a higher differentiation efficiency (% of cells successfully entering Neural Ectoderm lineage) (related to Fig. 2). Same data and procedure as described in Fig. S3. Each box shows the differentiation efficiency for a different starting population-density (indicated above each box). On each day, we collected all cells from a 10-cm diameter dish and then flowed them into a flow cytometer to measure the percentage of alive cells in the population that expressed GFP (i.e., percentage of cells that expressed Sox1 - a marker of NE-lineage commitment) (see STAR Methods). Differentiation efficiency reached a maximum ~80% for populations that started above the threshold density of ~1700 cells/cm^2^. Differentiation efficiency was below ~50% for populations that began with a below-threshold density. *n* is at least 3; error bars are s.e.m.

**Figure S5.**
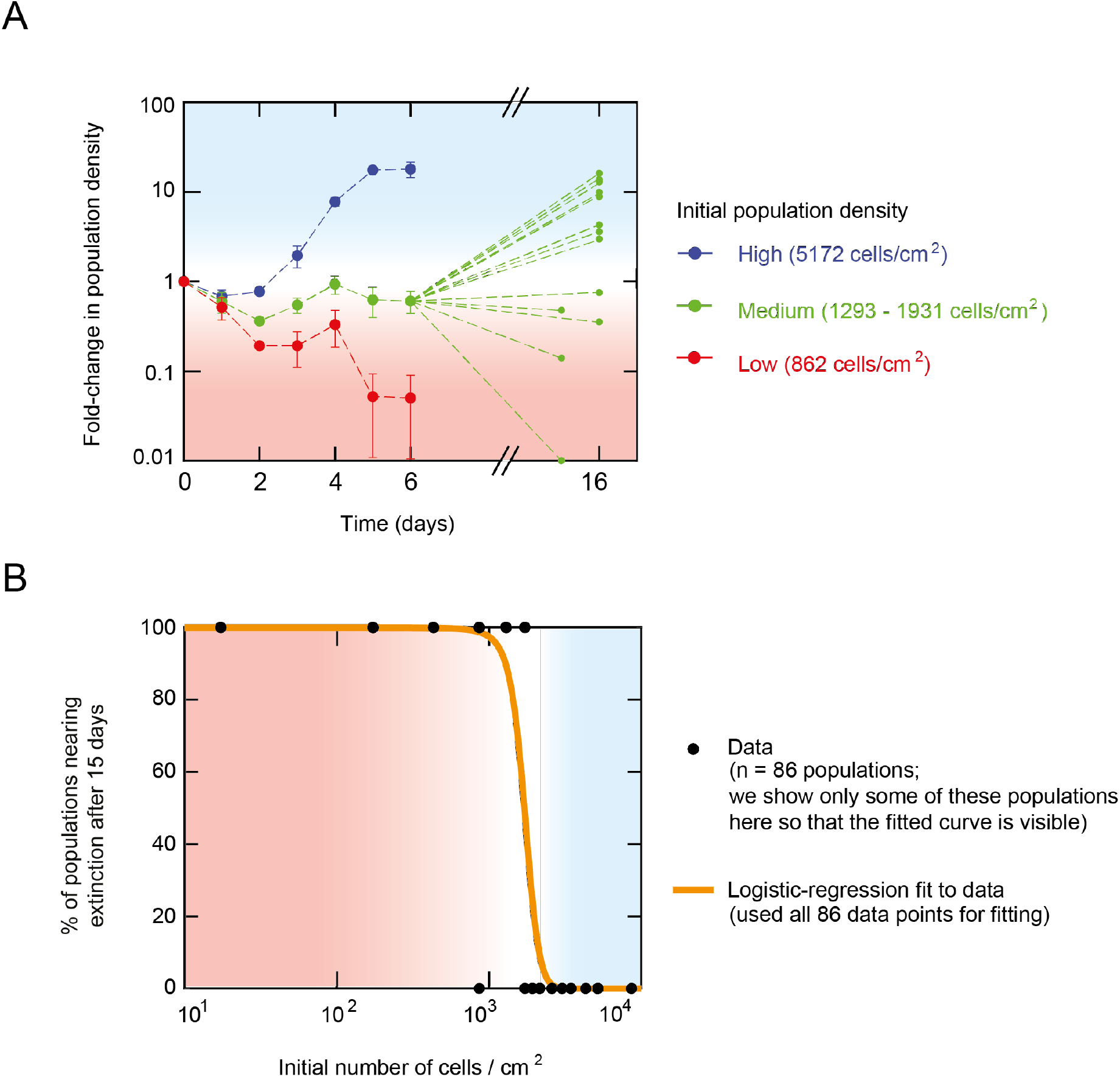
Differentiating populations that start with a (near) “threshold density” (~1700 cells/cm^2^) neither expand nor shrink during the first 6 days. But, after more days, these populations either expand towards the carrying capacity or shrink towards extinction in a stochastic manner (i.e., two populations of the same starting density can have two different fates (one survives and one becomes extinct)) (related to Figs. 2 and 3). Data for 46C cells (which have Sox1 promoter driving GFP expression) differentiating towards NE lineage in N2B27 + Retinoic Acid (RA) that were previously self-renewing in serum+LIF (see STAR Methods). Note that Sox1 is a marker of Neural Ectoderm lineage. **(A)** Blue: populations that started with a sufficiently high density (5172 cells/cm^2^) all grew towards the carrying capacity. Red: populations that started with a sufficiently low density (862 cells/cm^2^) became extinct within the first 6 days. Green: populations that started near a “threshold density” (between 1293 and 1931 cells/cm^2^) neither grew or shrank in the first 6 days. However, by 15-16 days after removing LIF (i.e., beginning differentiation), some of these populations reached the carrying capacity (i.e., population density increased by ~10-folds) while some others became extinct (i.e., population density became ~0.1-fold or less than its starting value). Still, some populations maintained nearly the same density for these 15-16 days (i.e., green curves with fold change of nearly one after 15-16 days). Thus, populations having virtually the same initial density can have distinct fates: some would become extinct and some would survive (i.e., the fate is stochastically determined). For all the data shown for the first six days, *n* = 3 and error bars are s.e.m. Each green data point, taken 15-16 days after triggering differentiation, represents a single population (to show the stochasticity) rather than being averaged over multiple populations. They thus do not show error bars. **(B)** To quantify the stochastic nature of the survival-versus-extinction fate for populations that start with a near-threshold density (green data in (A)), we measured the foldchange in population density for 86 populations that collectively spanned a wide range of initial densities (black points). To each population, we either assigned a value of zero if it eventually grew towards the carrying capacity (i.e., fold-change of larger than 1) or a value of one if it eventually approached extinction within the first 16 days of differentiation. Then, we performed a logistic regression on these black data points by fitting a logistic function, 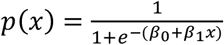 (orange curve). Consequently, *p*(*x*) represents the probability that a population approaches extinction by 15-16 days after differentiation began. By fitting, we found *β*_1_ = −8.7856 ± 0.8780 with a p-value of 3.76 x 10^-25^ according to the Wald test. This logistic regression is the simplest model (null model) that we can have for describing the probability of becoming extinct without any information about the mechanisms that determine the survival-versus-extinction fate. Later, we will introduce a mechanistic model that replaces this logistic regression fit (in Fig. 3).

**Figure S6.**
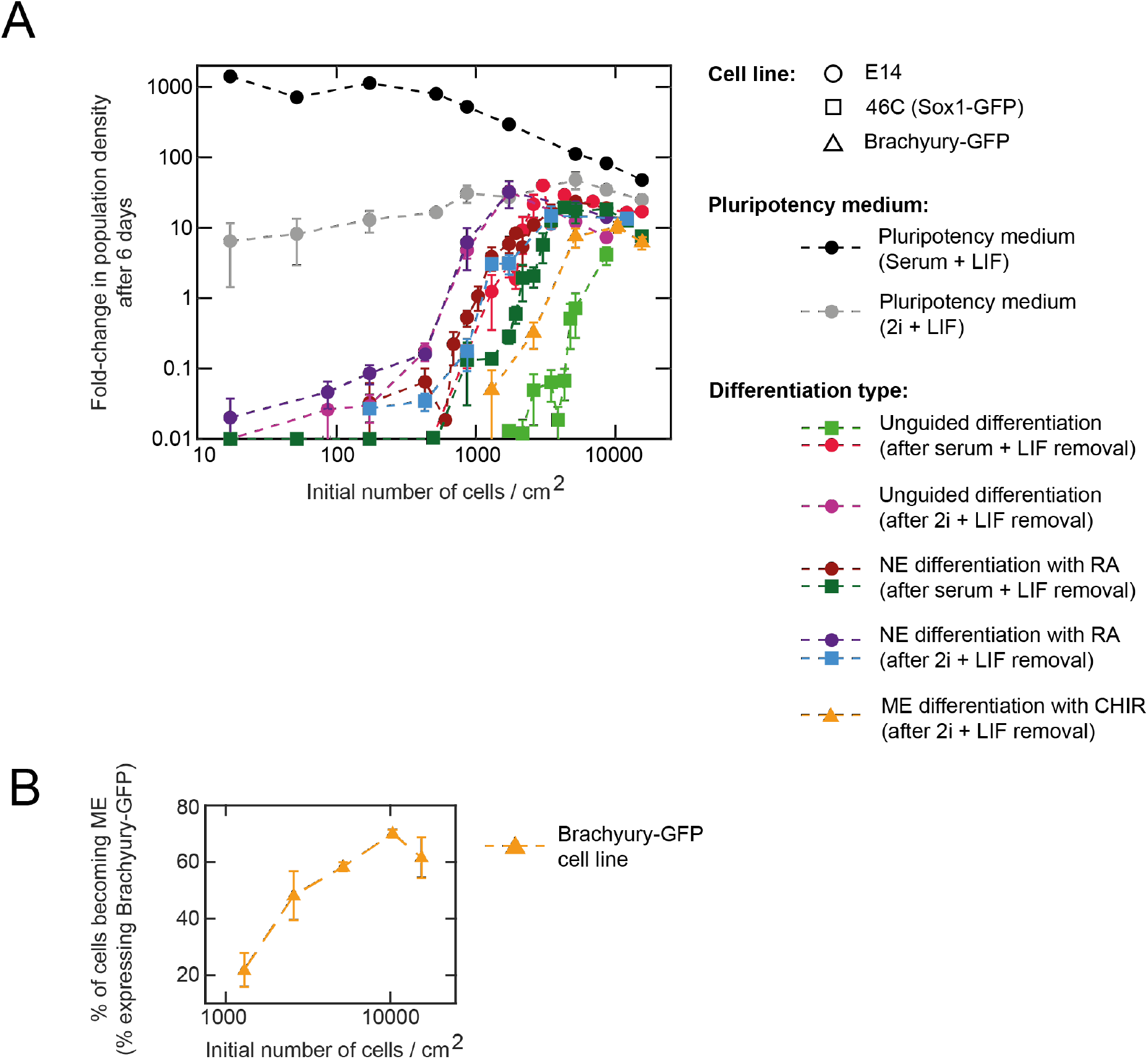
Different cell lines and multiple types of differentiation all exhibit the same phenomenon: differentiating population’s initial density determines its survival-versus-extinction fate (related to Fig. 2). **(A)** Data shown for three different cell lines: E14 (circles), 46C (squares), and Brachyury-GFP (triangles). The 46C cells have a Sox1 promoter controlling their GFP expression (Sox1 is a marker of the Neural Ectoderm (NE) lineage). The Brachyury-GFP cells have a Brachyury promoter controlling their GFP expression (gift from V. Kouskoff and described in (*57,58*). Brachyury is a marker of Mesendoderm (ME) lineage. Different colors represent different types of cell-culture media as indicated in the legend. For each differentiation, we took one of the three cell lines that were kept pluripotent with LIF in either a serum-based medium or a serum-free (2i) medium (indicated in legend). The three types of differentiations are: unguided differentiation in which no inducer was added after triggering pluripotency loss, NE differentiation in which we added Retinoic Acid (RA), and ME differentiation in which we added the small molecule, CHIR (see STAR Methods). As seen here, regardless of the cell line and differentiation type, a differentiating population’s initial density determined its survival-versus-extinction fate. **(B)** After four days of CHIR-induced differentiation, we used a flow cytometer to measure the percentage of the Brachyury-GFP cells (previously self-renewing in 2i+LIF) that expressed GFP (i.e., percentage of cells in a population that entered the ME lineage (*53*). As with the 46C cells that differentiated towards the NE lineage (Fig. 2C), populations of Brachyury-GFP cells that start with higher densities have higher differentiation efficiencies (larger percentages of cells entering the ME lineage). For both (A) and (B): *n* = 3; error bars are s.e.m.

**Figure S7.**
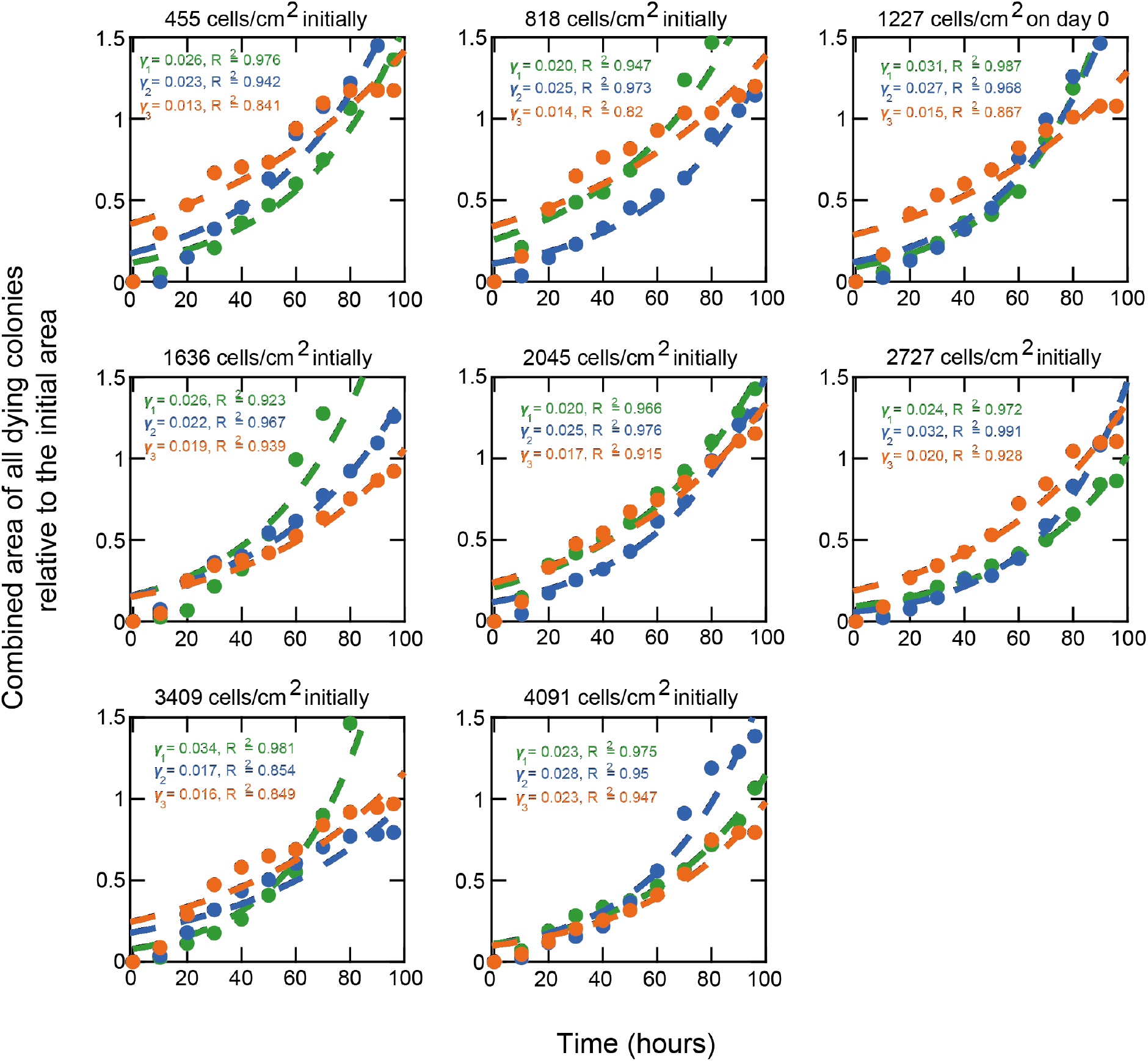
Determining the rate of cell death with time-lapse microscopy (related to Figs. 3 and 5). We sought to verify that the rate of cell death is independent of the initial population density. This is an ingredient of our mathematical model (see STAR Methods). With time-lapse microscopy, we measured the growth of individual microcolonies over four days for a wide range of initial population densities. Data for E14 cells during unguided differentiation in N2B27 (without any inducers such as RA) that were previously self-renewing in serum+LIF (see STAR Methods). Each box shows a population of a different starting density (indicated above each box). For each replicate of a population density (shown in colors green, blue and orange), we tracked microcolonies in 17 fields of view on a 6-cm diameter dish. Each field of view has a dimension of 1.40 mm x 0.99 mm. We examined all densities in triplicates (each color represents an independent replicate). Dying colonies typically lifted off the plate (and thus disappeared from the field of view) or started to display clearly visible apoptotic bodies. From these movies, we inferred the death rate of cells by taking the cumulative sum of the last recorded areas of each colony just before it died, as a way to measure how much colony area is “lost” (i.e., deaths) in time. We then divided this combined area of all dead cells by the combined initial area. We determined the death rate by fitting a single exponential function - with the death rate γ (in 1/hours) - to the data points (dashed curves represent the fits to data points of corresponding color; each color is a single replicate; *n* = 3 for each initial density) (see Fig. S8A for a demonstration of this procedure). The values of γ and corresponding R^2^ are indicated in each plot. Error bars are s.e.m. This figure complements Fig. S8.

**Figure S8.**
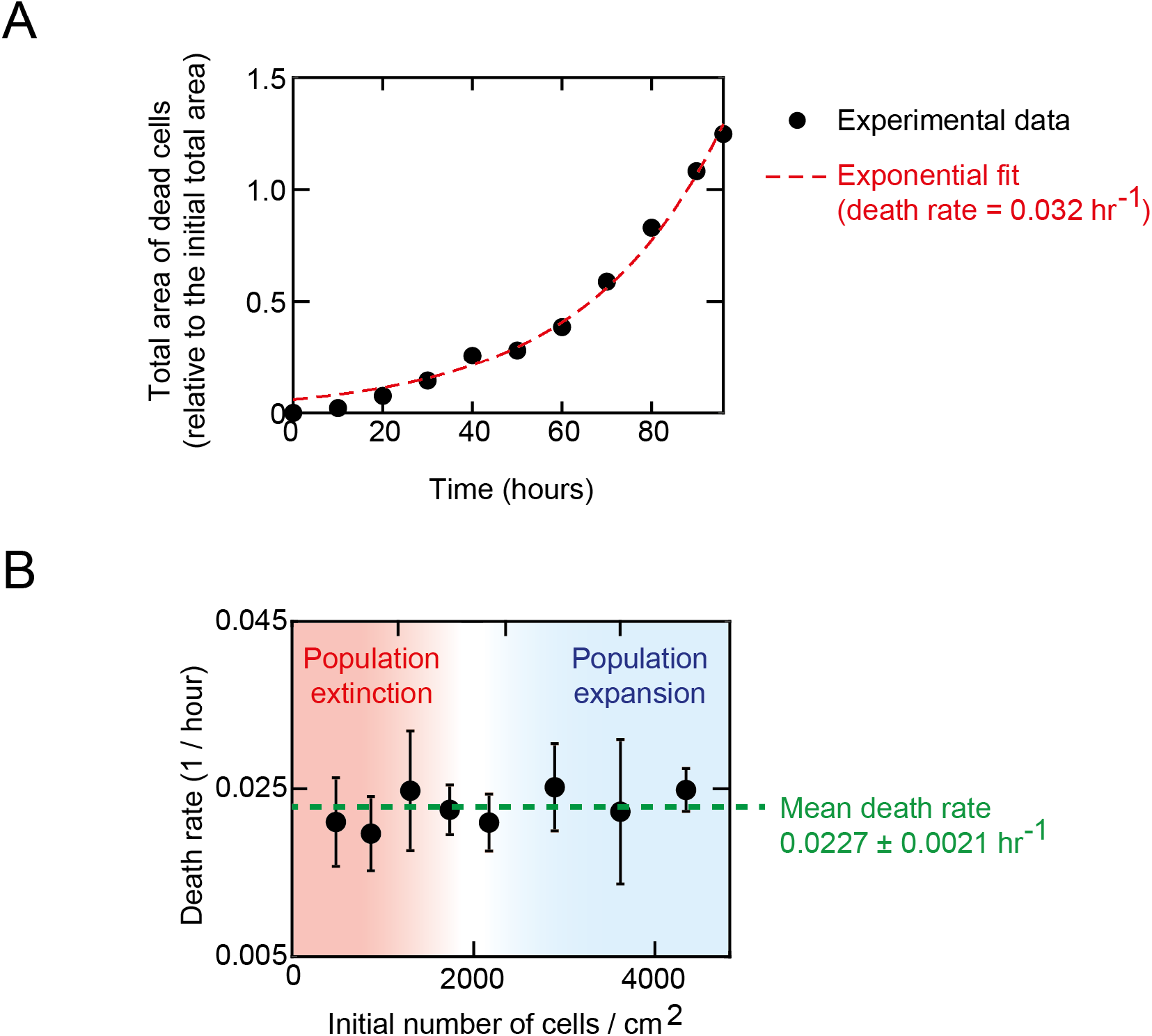
Rate of cell death is independent of initial population density (related to Figs. 3 and 5). Data for E14 cells during unguided differentiation in N2B27 (without any inducers such as RA) that were previously self-renewing in serum+LIF (see STAR Methods). **(A)** Example that shows how we inferred the death rate from measuring the area of dead cells per unit time for each initial population density. See explanation in the caption of Fig. S7. **(B)** Death rate extracted in Fig. S7 for many different initial population densities. This plot shows that the death rate of differentiating cells is constant (0.0227 ± 0.0021 hr^-1^). It is independent of initial population density. *n* = 3; Error bars are s.e.m.

**Figure S9.**
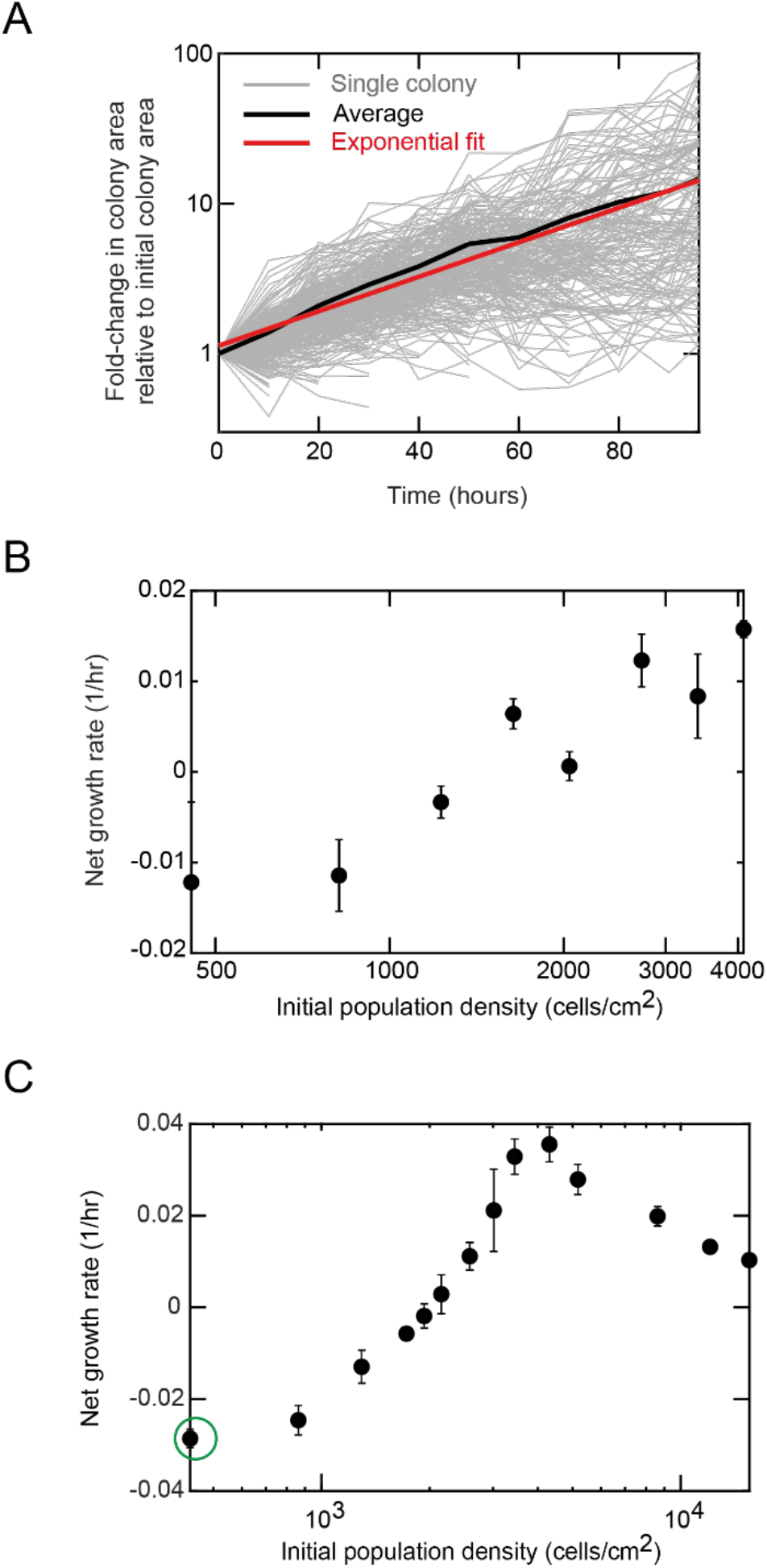
Density-dependent net growth rate (= growth rate - death rate) measured in two ways (related to Figs. 3 and 5). Data for E14 cells during unguided differentiation in N2B27 (without any inducers such as RA) that were previously self-renewing in serum+LIF (see STAR Methods). Net growth rate is negative if the death rate is higher than the growth rate (i.e., for a population that becomes extinct). It is positive if the growth rate is larger than the death rate (i.e., for a population that grows). We measured the cells’ net growth rates in two different ways. In one, we used time-lapse microscopy data to measure colony growths. In the other, we manually counted cells -and thus the population density over time as in Fig. 2C. **(A)** Example showing the microscope-based method of determining the net growth rate for a given initial population density. As shown in Fig. S10 and described in the caption there, we measured the area of each colony over time for four days, imaging 17 fields of view for each of the three replicates of a starting population density. Each grey curve shows the area of a single microcolony over time. Each colony’s area is normalized to its initial area (thus all curves here start at a value of one on the vertical axis). Black curve is the average of all the grey curves. Red line is an exponential curve (line in this semi-log plot) that we fitted to the black curve (i.e., fitted to the population average). The slope of the red line is the net growth rate for this population. **(B)** Using the microscope-based method outlined in (A) for every initial population density, we obtained the growth rate (black dots) as a function of the initial population density. *n* = 3; Error bars are s.e.m. **(C)** Net growth rates determined by manual counting of cells (i.e., data in Fig. S3) rather than from the microscopy data. *n* = 3; Error bars are s.e.m. To determine the (non-net) growth rate for each initial population density, we added the constant death rate (determined in Fig. S8) to the net growth rate (determined in (B-C)). We found that the maximum possible growth rate is 0.05219 hr^-1^, which we used as one of the parameter values in our mathematical model. From (C), we found that the lowest possible net growth rate is −0.026 hr^-1^ (indicated with green arrow in (C)). This value nearly matches the death rate (0.023 ± 0.002 hr^-1^; found in Fig. S8B), a consequence that we explore in our mathematical model (see STAR Methods).

**Figure S10.**
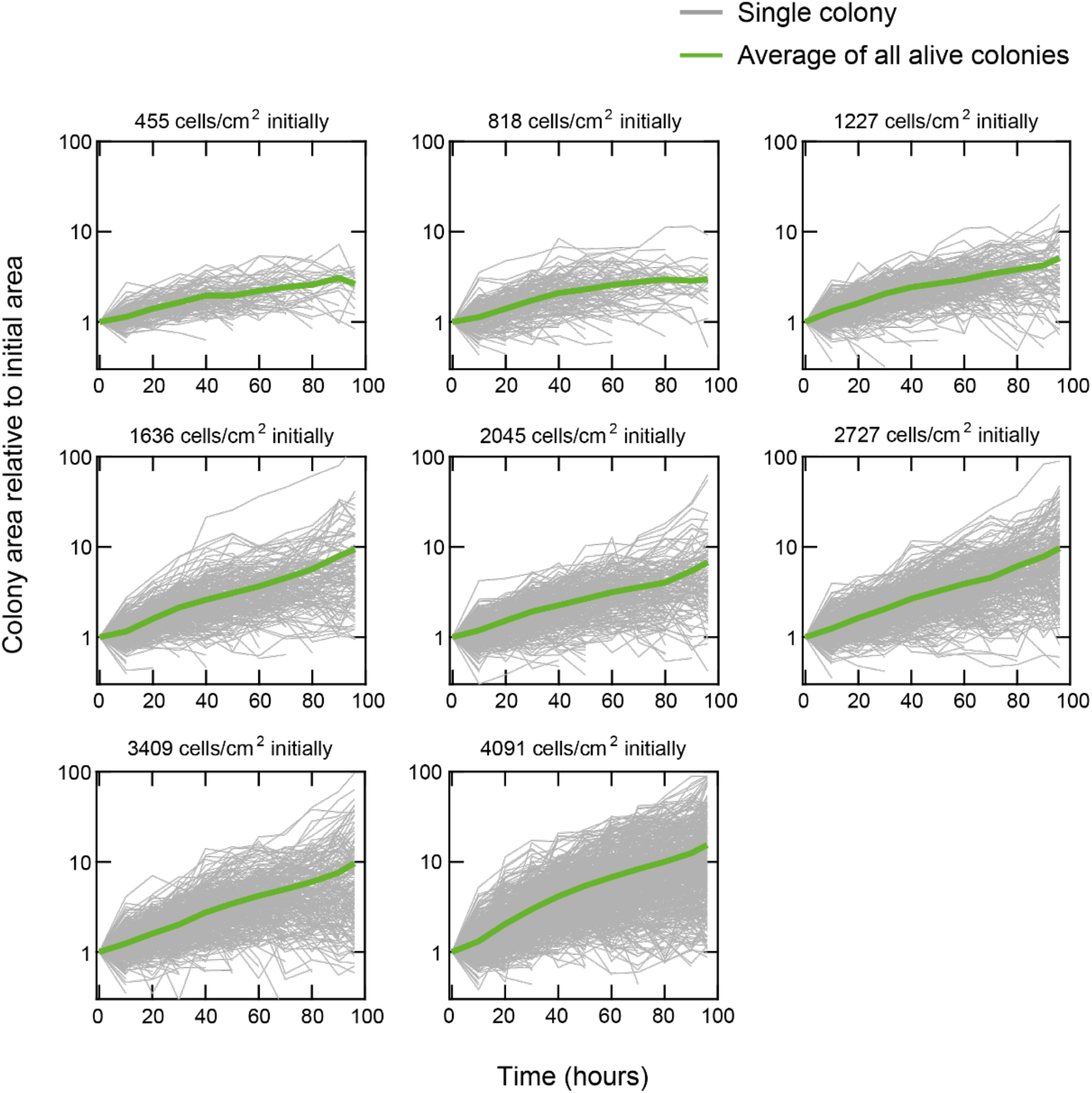
Time-lapse microscopy over four days reveals the growth rate of each microcolony for a wide range of initial population densities (related to Figs. 3 and 5). Data for E14 cells during unguided differentiation in N2B27 (without any inducers such as RA) that were previously self-renewing in serum+LIF (see STAR Methods). We used a wide-field microscope to image and measure the area of each microcolony over four days (each grey curve). During the four days, we measured the area of a given microcolony every 10 hours (see STAR Methods). Each box shows the fold-change in the colony area (vertical axis) as a function of the time passed since triggering differentiation (horizontal axis) for different starting population densities. In these movies, we observed both growing and dying colonies. For the colonies that died, the grey curves abruptly end, at the last time frame in which they were alive and before the end of the four-day period. The dying colonies visibly stood out as they typically displayed apoptotic bodies or lifted off the plate and thus disappeared from the focal plane. At each time frame, we computed the average area of all living colonies (green curve) from which we can extract the average growth rate of a colony for each population density.

**Figure S11.**
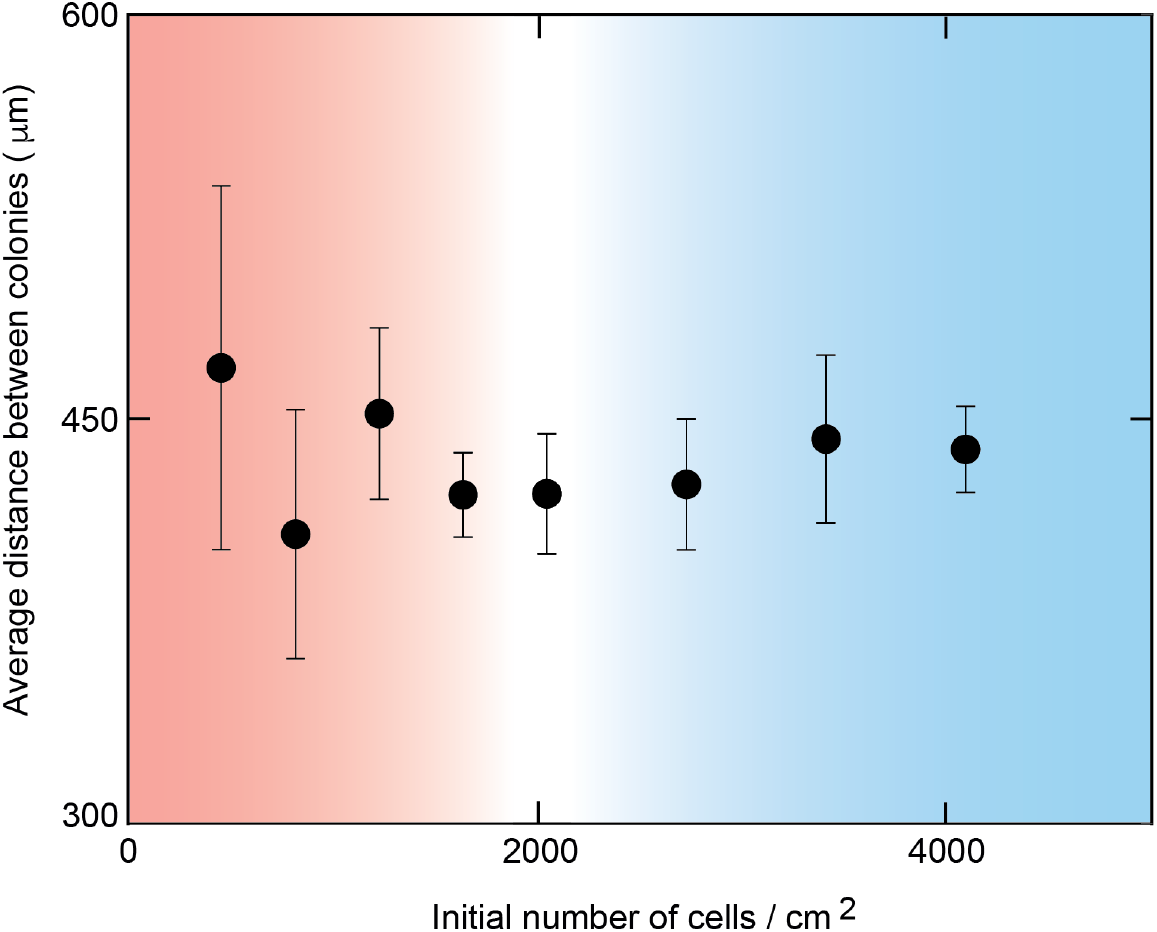
For every population density, microcolonies are hundreds of microns apart from each other when differentiation begins (related to Figs. 2 and 4). Data for E14 cells during unguided differentiation in N2B27 (without any inducers such as RA) that were previously self-renewing in serum+LIF (see STAR Methods). We scattered a relatively a desired number of cells across a 6-cm diameter dish containing N2B27 to trigger pluripotency loss. Then, we used a wide-field microscope to locate and image the microcolonies in 17 fields of view. Each field of view has a dimension of 1.40 mm x 0.99 mm. From these images, we determined the distance between every pair of colonies that resided in the same field of view. Then, we averaged these distances (averaging over all pairs of colonies from all 17 fields of view per plate). The resulting, average distance between colonies is plotted here as a function of the initial population density. As shown here, for a wide range of initial population densities, the average distance between microcolonies were virtually identical (~450 μm). *n* = 3 plates for each initial population density; error bars are s.e.m. A way to understand why the average colony-colony distance is nearly independent of the initial population density is that the dish area is much larger than can be covered by the cells, even for the high-density populations (at most ~1% of plate area is covered by the cells, as one can also check by a back-of-envelope calculation). In short, the vast difference in length-scale between individual cells and the plate area leads to above result. Given this, more informative metric is actually the distance between a colony and its nearest-neighboring colony (see Fig. S14).

**Figure S12.**
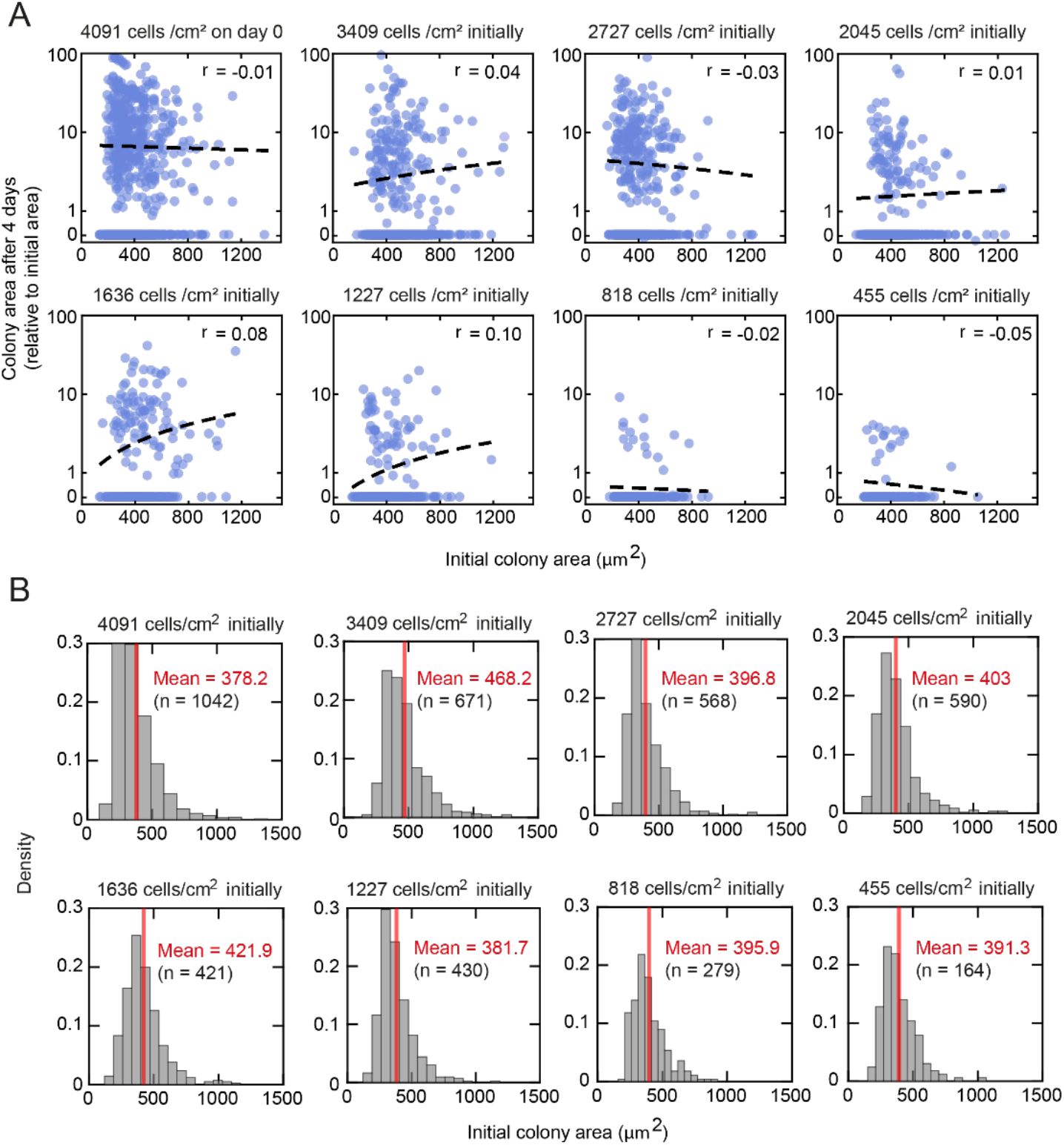
Growth and survival of differentiating cells within a colony do not depend on how many cells are initially in the colony (i.e., colony area) for any population density (related to Fig. 4). **(A, B)** Data for E14 cells during unguided differentiation in N2B27 (without any inducers such as RA) that were previously self-renewing in serum+LIF (see STAR Methods). Same protocol as in Fig. S10. We used a time-lapse microscope to measure the colony area during four days of differentiation. Each blue point represents the final area (after 96 hours) of a colony relative to its initial area (i.e., fold-change in colony area compared to the initial area). If a colony died during the time-lapse (spotted as explained in caption for Fig. S10), then we assigned it a value of zero as the fold-change in its area. Each box shows the fold-change in the colony area (vertical axis) as a function of the initial colony area (horizontal axis) for 17 fields of view and a specific population density. Dashed curves in each box denote the linear regression with the Pearson correlation coefficient *r* in each box. There is virtually no correlation between a colony’s final area (and whether it dies or not) and its initial area, for any starting population density. These results suggest that the initial area of a microcolony – which in turn is set by the number of cells in a microcolony – does not predict whether any cells in the colony survive or not and also does not predict how fast each cell grows during differentiation.

**Figure S13.**
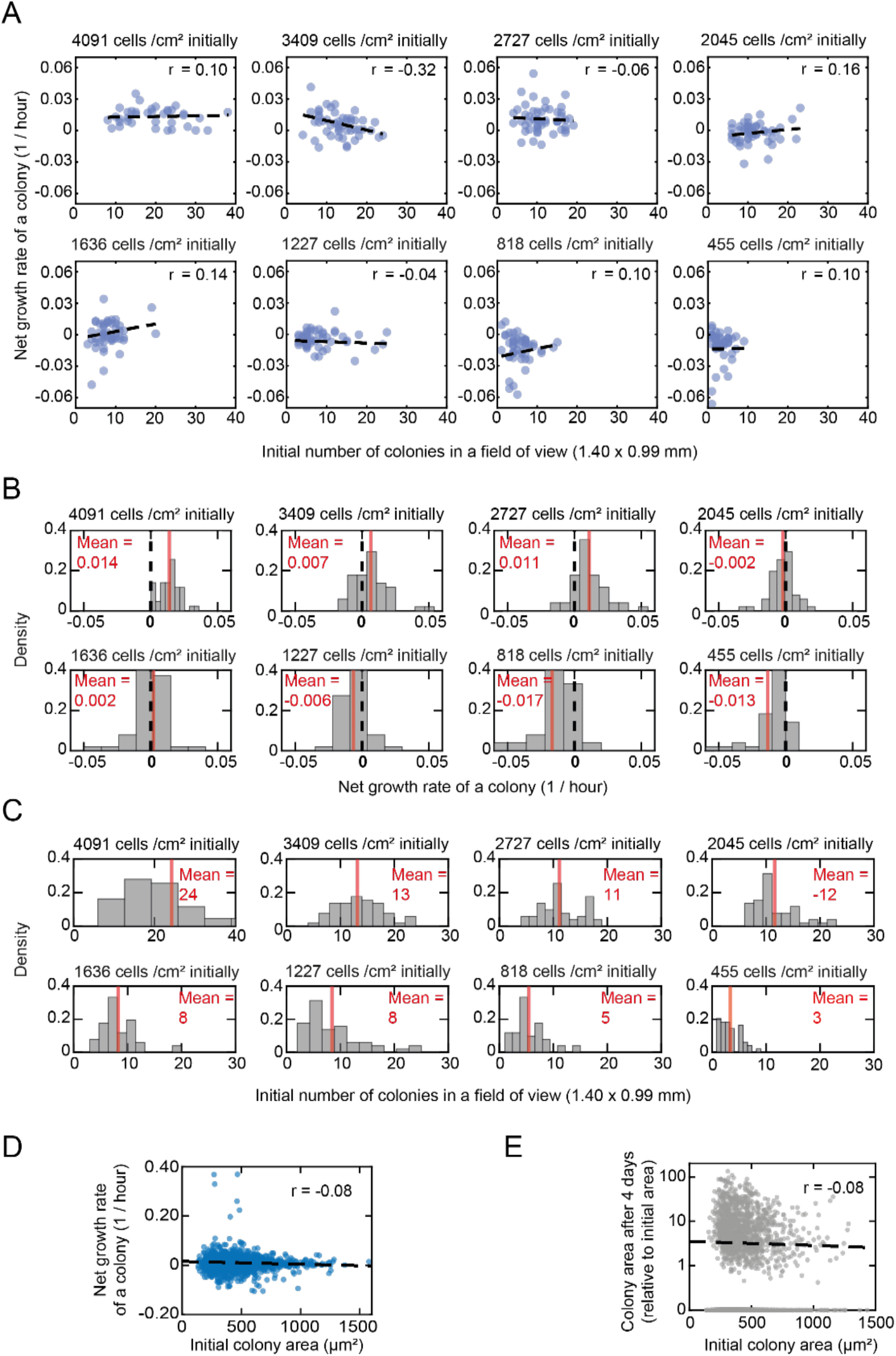
Net growth rate of a colony does not depend on how many cells are in the colony (i.e., colony area) for any population density (related to Fig. 4). Data for E14 cells during unguided differentiation in N2B27 (without any inducers such as RA) that were previously self-renewing in serum+LIF (see STAR Methods). Same protocol as in Fig. S10. **(A, B, C)** We fitted an exponential function, Area(t) = Aoexp(μt), to each grey trace shown in Fig. S10 (i.e., we estimated colony area as exponentially growing as a function of time). From this fit, we determined net growth rate μ for each colony (i.e., for every grey trace shown in Fig. S10). For a given colony, we determined its net growth rate and the initial number of colonies that resided in the same field of view. Each blue point shows the data for a single colony. The net growth rate is positive (μ > 0) if the colony grew over the four days of differentiation whereas it is negative (μ < 0) or zero if the colony died (i.e., shrank or did not grow after which it often detached from the plate and thus disappeared from the field of view). Each box shows a colony’s net growth rate (vertical axis) as a function of the initial number of colonies that resided in the same field of view (horizontal axis) for a specific population density. We analyzed 17 fields of view for each starting population density. Dashed lines in each box shows the Pearson correlation with the correlation coefficient *r* denoted in each box. Here we see that how fast a colony grows - and whether it survives or dies - is virtually uncorrelated with how many other colonies there are in its ~1 mm x 1 mm neighborhood. **(D)** There is virtually no correlation between a colony’s net growth rate and its initial area, as shown here. Data points come from many different, initial population densities and pooled together here. Dashed line shows the Pearson correlation with the correlation coefficient *r* = −0.08. **(E)** There is virtually no correlation between a colony’s final area relative to its initial area and its initial area, as shown here. Data points come from many different, initial population densities and pooled together here. Dashed line shows the Pearson correlation with the correlation coefficient *r* = −0.08.

**Figure S14.**
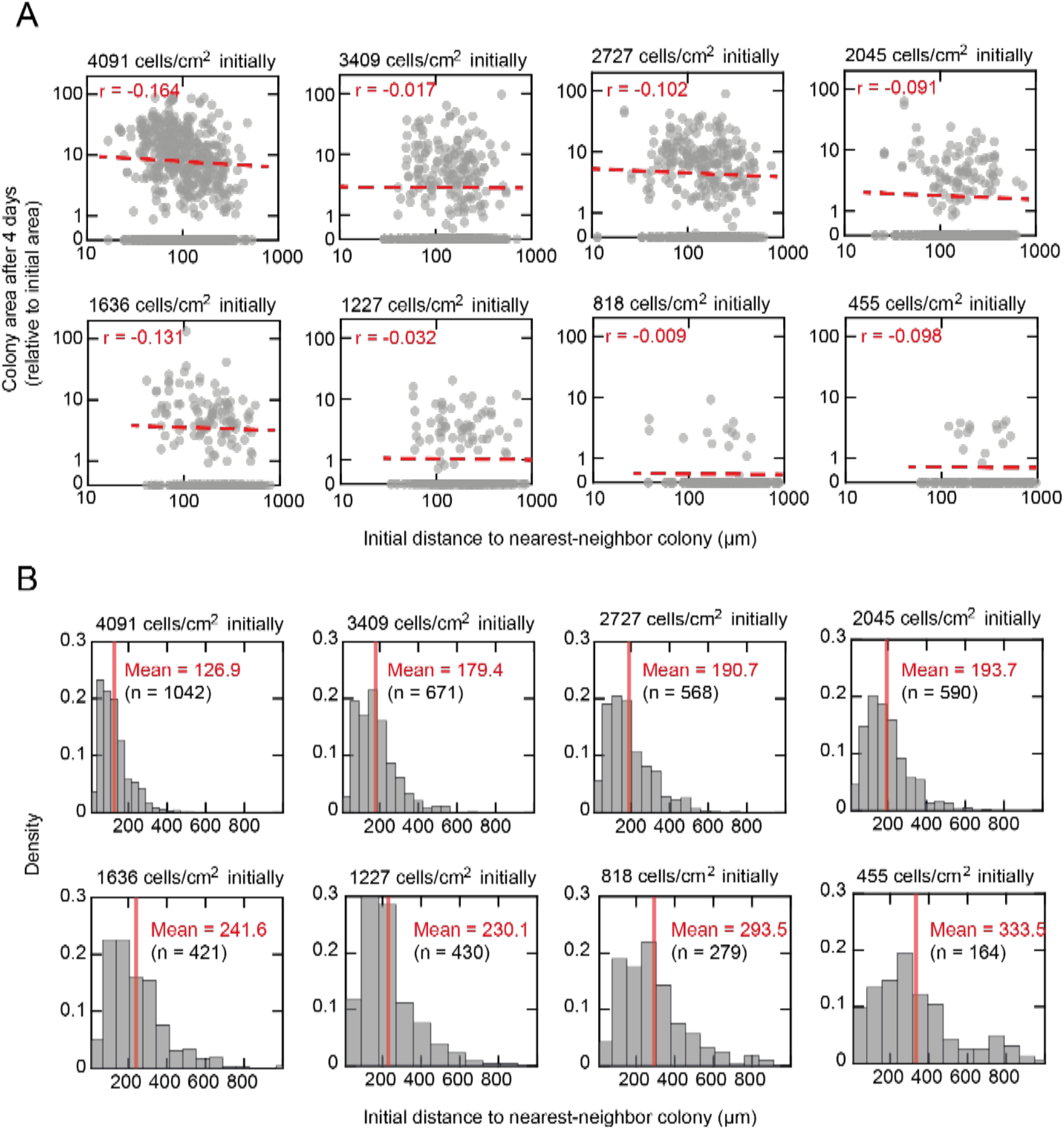
Growth and survival of differentiating cells do not depend on the distance between nearest-neighboring colonies for any initial population density (related to Fig. 4). **(A, B)** Data for E14 cells during unguided differentiation in N2B27 (without any inducers such as RA) that were previously self-renewing in serum+LIF (see STAR Methods). Same protocol as in Fig. S10. We scattered a desired number of cells across a 6-cm diameter dish containing N2B27 to initiate exit from pluripotency. Then, we used a wide-field microscope to locate and image the microcolonies in 17 fields of view (see STAR Methods). Each field of view has a dimension of 1.40 mm x 0.99 mm. From these images, we determined the distance between every pair of colonies that resided in the same field of view. From these measurements, we found the smallest distance (i.e., distance from one colony to its nearest-neighboring colony). We did this for each colony in a field of view. The resulting, nearest-neighboring distance for each colony is plotted here as a function of the initial population density. Each grey point represents the final area (after 96 hours) of a colony relative to its initial area as a function its distance to its nearest neighbor (at the start of the time-lapse movie). If a colony died during the time-lapse, then we assigned it a value of zero as the foldchange in its area. Each box shows the fold-change in the colony area (vertical axis) as a function of the distance from colony to its nearest neighbor (horizontal axis) for 17 fields of view and a specific population density. Dashed curves in each box denote the Pearson correlation with the correlation coefficient *r* in each box. There is virtually no correlation between a colony’s final area (and whether it dies or not) and its initial, nearest-neighbor distance, for any starting population density. These results strongly indicate that the presence of a nearby colony does not predict whether a colony survives or not and also does not predict how fast each colony grows during differentiation.

**Figure S15.**
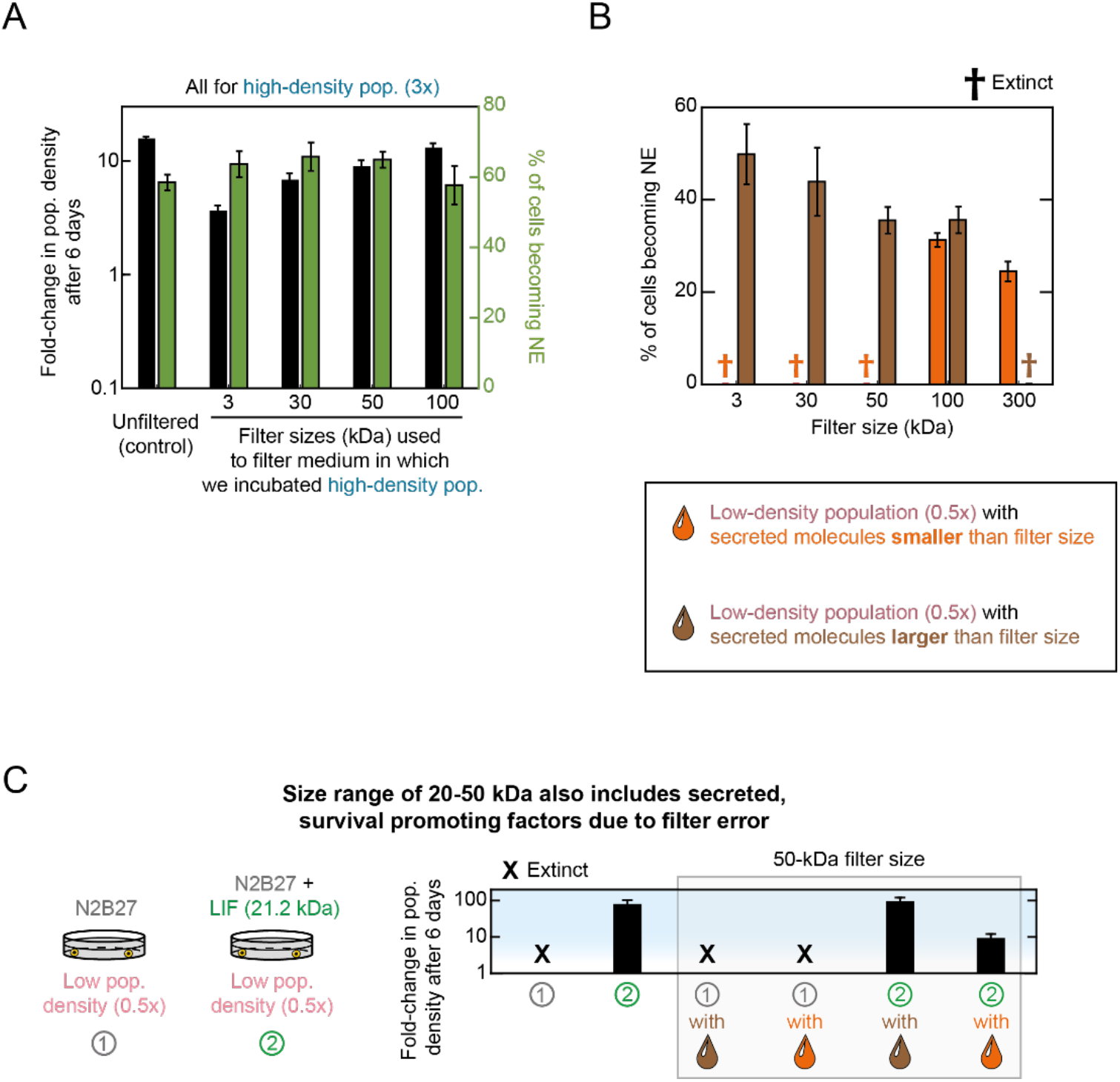
Filtering culture medium does not eliminate media components that are essential for cell growth (related to Fig. 5). We used commercial, membrane-based filters of various sizes (Fig. 5A-B, see STAR Methods). Data in (A) and (B) for 46C cells (which have Sox1 promoter driving GFP expression) differentiating towards NE lineage in N2B27 + Retinoic Acid (RA) that were previously self-renewing in serum+LIF (see STAR Methods). **(A)** To check that the filters do not remove any essential media components for cell growth, we filtered the medium of a high-density population (5172 cells/cm^2^) with filters of various sizes, after two days of differentiation. We then took the filtered medium (medium that passed through the filter and thus containing all molecules that are smaller than the filter size), and gave it back to the same high-density population. We incubated the population for four days in this medium. On the last day, we measured the fold-change in population density (black bars) and the percentage of cells becoming NE (green bars). As a control, we also measured the fold-change in population density after six days of growth in unfiltered medium (first black and green bars). Since all black bars have nearly the same height as do all the green bars, we can conclude that none of the filters catch any ingredients in the cell-culture medium that are essential for cell growth (e.g., vitamins and other components which are already present in the medium from the beginning of cell culture, rather than secreted by cells). **(B)** We cultured a low-density population (862 cells/cm^2^) in a medium from a high-density population (5172 cells/cm^2^) that went through the filter (orange bars). This medium has all molecules that are smaller than the filter size. We also cultured the low-density population in a medium that contained all the molecules that are larger (heavier) than the filter size (brown bars), which we captured by flowing the high-density population’s medium through the filter. We used a wide range of filter sizes (horizontal axis). The bars show the percentages of cells entering the NE after being cultured in the filtered medium. Crosses indicate that the population became extinct in the filtered medium. The corresponding fold-changes in population density are shown in Fig. 5B. **(C)** To check the accuracy of the filters in capturing a molecule of known size and effect, we filtered N2B27 with a 50-kDa size filter (we show in (B) that this procedure does not catch any ingredients in N2B27 that are essential for cell growth) that was supplemented with recombinant LIF (1000 U/mL). We then measured the fold-change in population density (black bars). Same color scheme is used as shown in (B). Recombinant LIF has a molecular weight of ~20 kDa and we confirmed that unfiltered N2B27+LIF results in a ~100 fold expansion of a low-density population (46C cells, previously self-renewing in serum+LIF) that otherwise becomes extinct during differentiation (here, N2B27 without any inducer such as RA). Both fractions of filtered N2B27+LIF resulted in the rescue and expansion of the low-density population, but the fraction containing molecules larger (heavier) than 50 kDa resulted in the largest possible expansion which was virtually identical to the expansion achieved with unfiltered N2B27+LIF. In light of the main results shown in Fig. 5A-B, we concluded that secreted (survival-promoting) molecules can have molecular weights as low as ~20 kDa.

**Figure S16.**
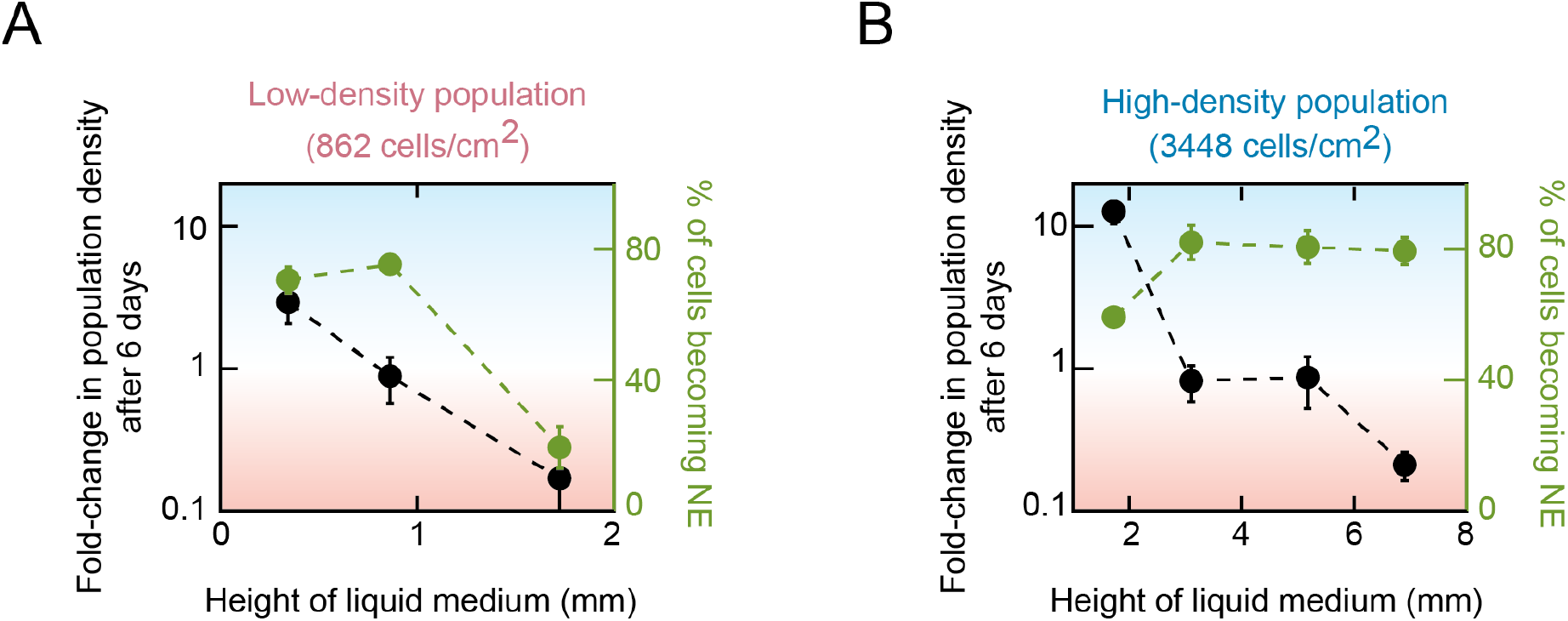
Changing the height of the cell-culture medium over several millimeters alters the survival-versus-extinction fate of the population on the bottom of the cellculture dish (related to Fig. 5). Data for 46C cells (which have Sox1 promoter driving GFP expression) differentiating towards NE lineage in N2B27 + Retinoic Acid (RA) that were previously self-renewing in serum+LIF (see STAR Methods). Black data points are duplicates of the data shown in Fig. 5F and they indicate the fold-change in the population density (relative to the initial population density) as a function of the liquid-medium height above the cells. Green data points indicate the percentage of the cells that expressed GFP (i.e., Sox1 - a marker for NE lineage), which we measured with a flow cytometer, as a function of the liquid height above the cells. In our study, we used a 10-mL liquid medium (e.g., in Fig. 2) unless we explicitly state that we used a different volume (e.g., in Fig. 5F and here). A 10-mL liquid has a height that is just below 2 mm in a 10-cm diameter dish. **(A)** Data for a differentiating population that starts with a low density (862 cells / cm^2^). In a 10-mL liquid, this population becomes extinct (last data point, at ~2-mm liquid height). From the smallest to the largest liquid height, the data correspond to 2 mL, 5 mL, and 10 mL liquid media. **(B)** Data for a differentiating population that starts with a high density (3448 cells / cm^2^). In a 10-mL liquid, this population survives and grows towards the carrying capacity (first data point, at ~2-mm height). From the smallest to the largest liquid height, the data correspond to 10 mL, 20 mL, 30 mL, and 40 mL liquid media. (A) shows that we can rescue a low-density population - it survives - if we decrease the liquid height by 50% or more from the usual, ~2-mm. (B) shows that we can drive a high-density population to barely survive or become extinct if we increase the liquid height by a 2-fold or more. Altogether, these results exclude local communication. They suggest that the key secreted molecules diffuse over at least several millimeters to control a population’s survival-vs-extinction fate.

**Figure S17.**
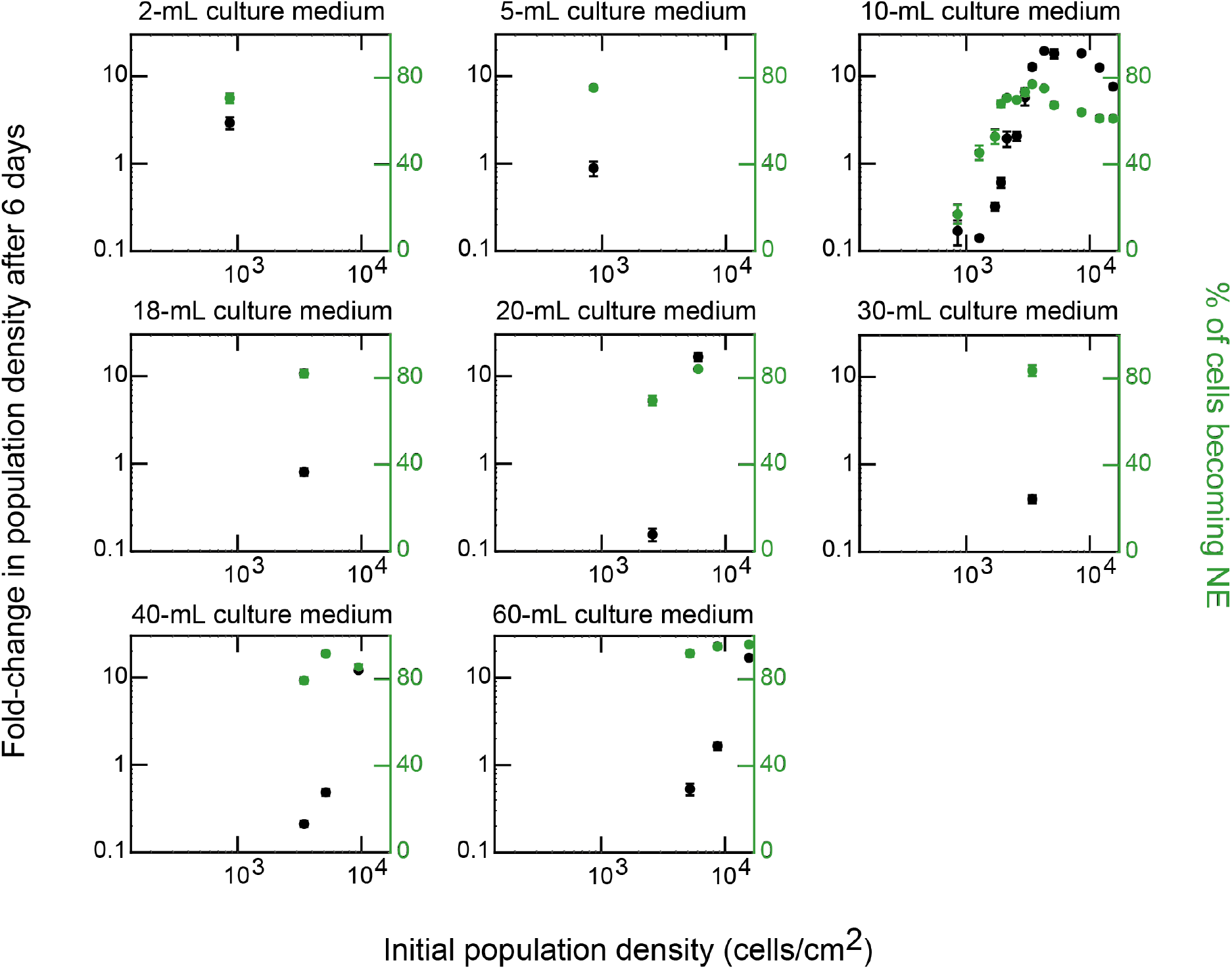
Volume of liquid medium at the start of differentiation determines survival-vs-extinction fate of differentiating populations (related to Fig. 5)). Data for 46C cells (which have Sox1 promoter driving GFP expression) differentiating towards NE lineage in N2B27 + Retinoic Acid (RA) that were previously self-renewing in serum+LIF (see STAR Methods). To experimentally test the model’s prediction (i.e., to verify that the model-produced phase diagram in Fig. 5H is correct as is the model’s prediction in Fig. 5G), we incubated 46C populations of different starting densities in different volumes of liquid medium (and thus in different heights of liquid medium). Each box shows a different volume of liquid medium. Horizontal axis of each box is the initial population density. Black points are the fold changes in the population density (relative to the initial density) after six days. Green points are the percentages of cells that entered the NE lineage (Sox1-GFP positive cells measured with flow cytometer). *n* = 3; Error bars are s.e.m.; *n* = 3. As seen in Fig. 5H, these results match the phase diagram produced by the model (i.e., data points of the right type fall on the right side (above or below) the model-produced phase boundary in Fig. 5H).

**Figure S18.**
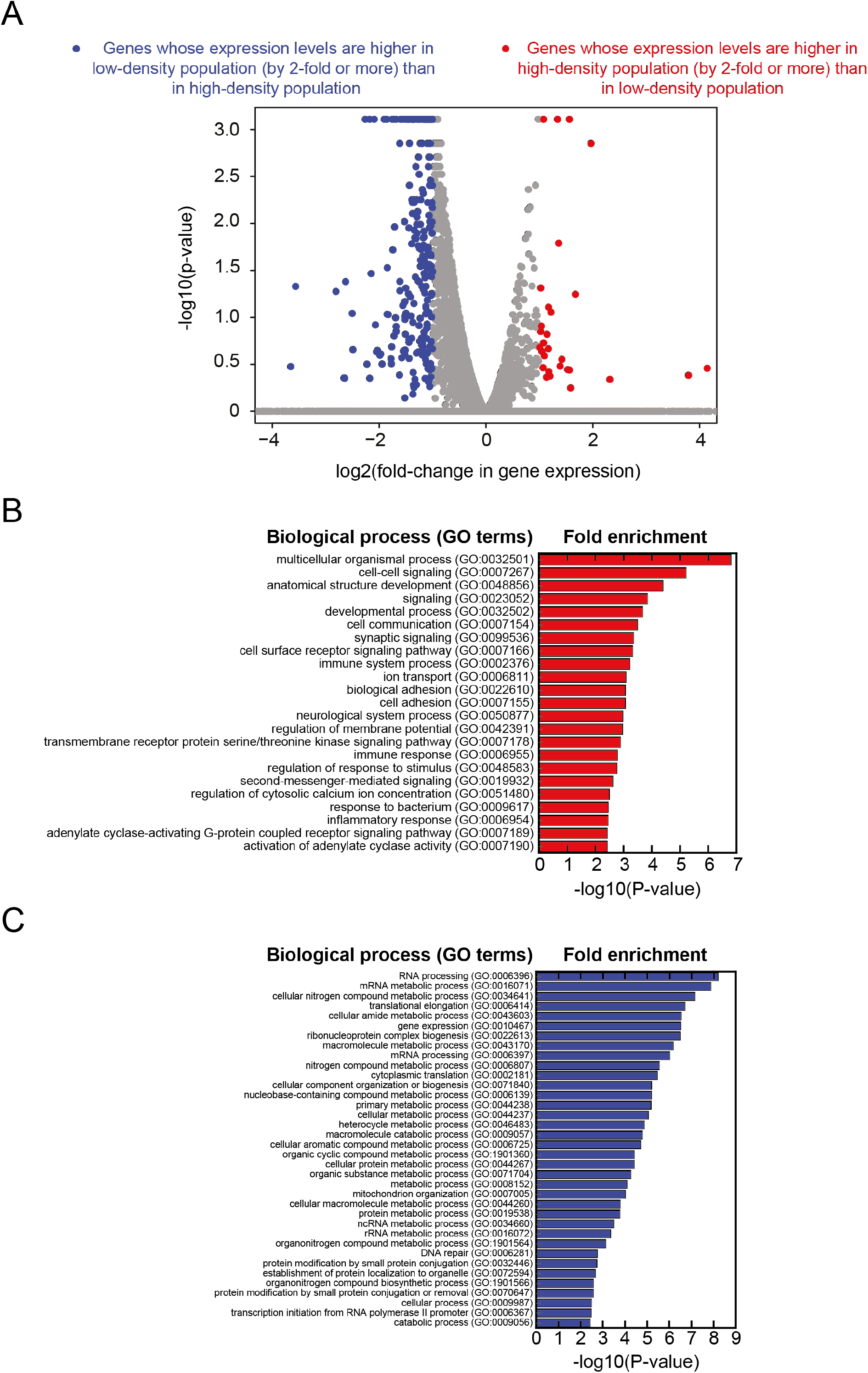
Enrichment analysis of RNA-Seq data reveals that high-density populations, compared to low-density populations, have higher levels of processes (GO terms) such as multicellular organismal processes, cell-cell signaling, neurological system processes, and cell adhesion (related to Fig. 6). Data for 46C cells (which have Sox1 promoter driving GFP expression) differentiating towards NE lineage in N2B27 (without any inducers such as RA) that were previously self-renewing in serum+LIF (see STAR Methods). We identified possible, intracellular pathways that the secreted molecules control by performing a transcriptome-level profiling (RNA-Seq) to examine differentially expressed genes. We performed RNA-Seq on populations of three different starting densities: (1) a low-density (862 cells/cm^2^) population that becomes extinct; (2) a high-density (5172 cells/cm^2^) population that grows towards the carrying capacity; and (3) a medium-density (1931 cells/cm^2^) population that is near the threshold density. For RNA-Seq, we collected all cells from these populations on the first and second days after triggering differentiation. We also collected cells that were kept pluripotent in a serum-based pluripotency medium (FBS with LIF) as a comparison. We analyzed the resulting transcriptome expression levels (FPKMs) for each gene by focusing on gene-expression levels that differed by more than 2-folds between the high- and low-density populations. **(A)** Volcano plot. Each grey dot represents a single gene. The horizontal axis shows the relative expression level: the expression level of the high-density population divided by the expression level of the low-density population. The vertical axis shows the p-value of the statistical test performed in Cufflinks (see STAR Methods). Genes that are more highly expressed by the high-density population than the low-density population, by 2-folds or more, are shown as red points. Genes that are more highly expressed by the low-density population than the high-density population, by 2-folds or more, are shown as blue points. **(B)** Enrichment analysis for genes that are more highly expressed by the high-density population than the low-density population by 2-folds or more (red data points in (A)). We used PANTHER and a custom MATLAB script to examine which GO terms (biological processes) are enriched and determine the corresponding significance (p-value) of the enrichment. We list here the enriched GO terms with their GO-term numbers. This plot shows that GO terms such as “multicellular organismal processes”, “cell-cell signaling”, “neurological system processes” and “cell adhesion” have some of the highest fold enrichments. **(C)** Enrichment analysis for genes that are more highly expressed by the low-density population than the high-density population by 2-folds or more (blue data points in (A)). We list here the enriched GO terms with their GO-term numbers. This plot shows that GO terms such as “RNA processing” and “macromolecule catabolic processes” have some of the highest fold enrichments. See also Fig. S19 for further analyses of the RNA-Seq dataset.

**Figure S19.**
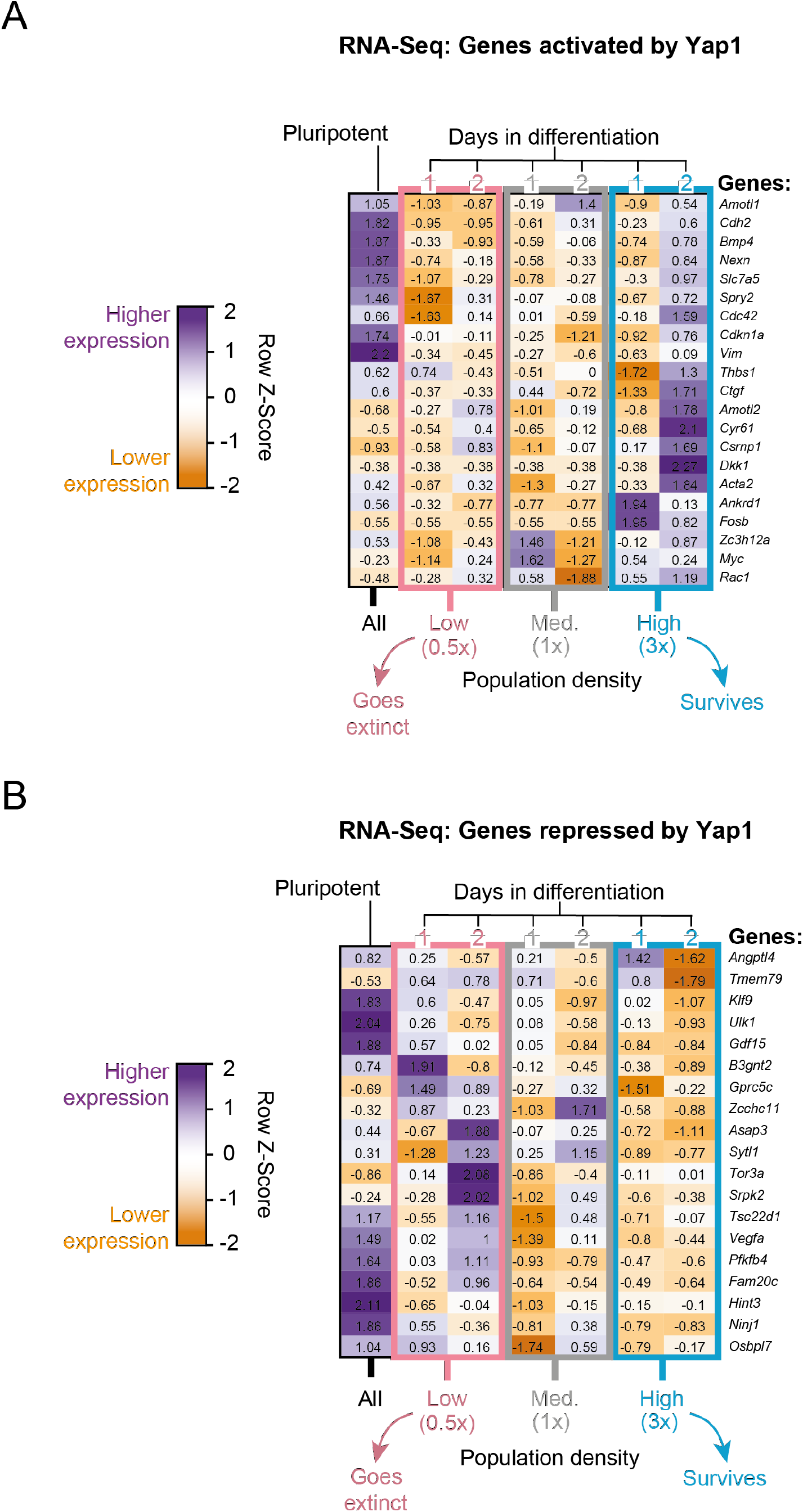
RNA-Seq analysis reveals that YAP1 becomes more active in populations that start with higher densities (related to Fig. 6). Data for 46C cells (which have Sox1 promoter driving GFP expression) differentiating towards NE lineage in N2B27 (without any inducers such as RA) that were previously self-renewing in serum+LIF (see STAR Methods). Same RNA-Seq dataset as in Fig. S18. YAP1 is a key component of the Hippo signaling pathway that is important for cell proliferation and apoptosis. We zoomed into YAP1-related genes in our RNA-Seq dataset because the analysis of enriched GO terms (Fig. S18) revealed several YAP1-related genes prominently participating in the “cell adhesion” processes (one of the top enriched GO terms). Details of the RNA-Seq are in the caption for Fig. S18. In brief, we performed RNA-Seq on populations of three different starting densities: (1) a low-density (862 cells/cm^2^) population that becomes extinct; (2) a high-density (5172 cells/cm^2^) population that grows towards the carrying capacity; and (3) a medium-density (1931 cells/cm^2^) population that is near the threshold density. For RNA-Seq, we collected all cells from these populations on the first and second days after triggering differentiation. We also collected cells that were kept pluripotent in a serum-based pluripotency medium (FBS with LIF) as a comparison. We analyzed the expression levels (FPKMs) by classifying genes into two groups: (1) genes that are activated by YAP1; and (2) genes that are repressed by YAP1 (*60–67*). For each gene, we compute its mean expression level μ by averaging the its expression level across all experimental conditions (i.e., across all densities and days). Afterwards, we determined the row Z-score for each gene and experimental condition, which is a measure of by how much a gene’s expression level in a given experimental condition deviates from the mean expression level (μ) for that gene. A gene that is more highly expressed has a high row Z-score (close to ~ 2) and is indicated as a shade of purple in the heat maps here. A gene that is more lowly expressed has a low row Z-score (close to −2) and is indicated as a shade of orange in the heat maps here. **(A)** Heat map that shows the row Z-score for each gene (each row) and each experimental condition (each column). These genes are known to be either directly or indirectly activated by YAP1. We observed that these genes, including *Cyr61* and *Amotl2*, were more highly expressed (purple color) by the higher-density populations than the lower-density populations, suggesting that YAP1 is more active in higher-density populations. **(B)** Heat map that shows the row Z-scores for each gene (each row) and each experimental condition (each column). These genes are known to be either directly or indirectly repressed by YAP1. We observed that these genes, including *Angptl4* and *Tmem79*, were more highly expressed (purple color) by lower-density populations than the higher-density populations. Taken together, the results here (A-B) are consistent with YAP1 becoming more active in higher-density populations.

**Figure S20.**
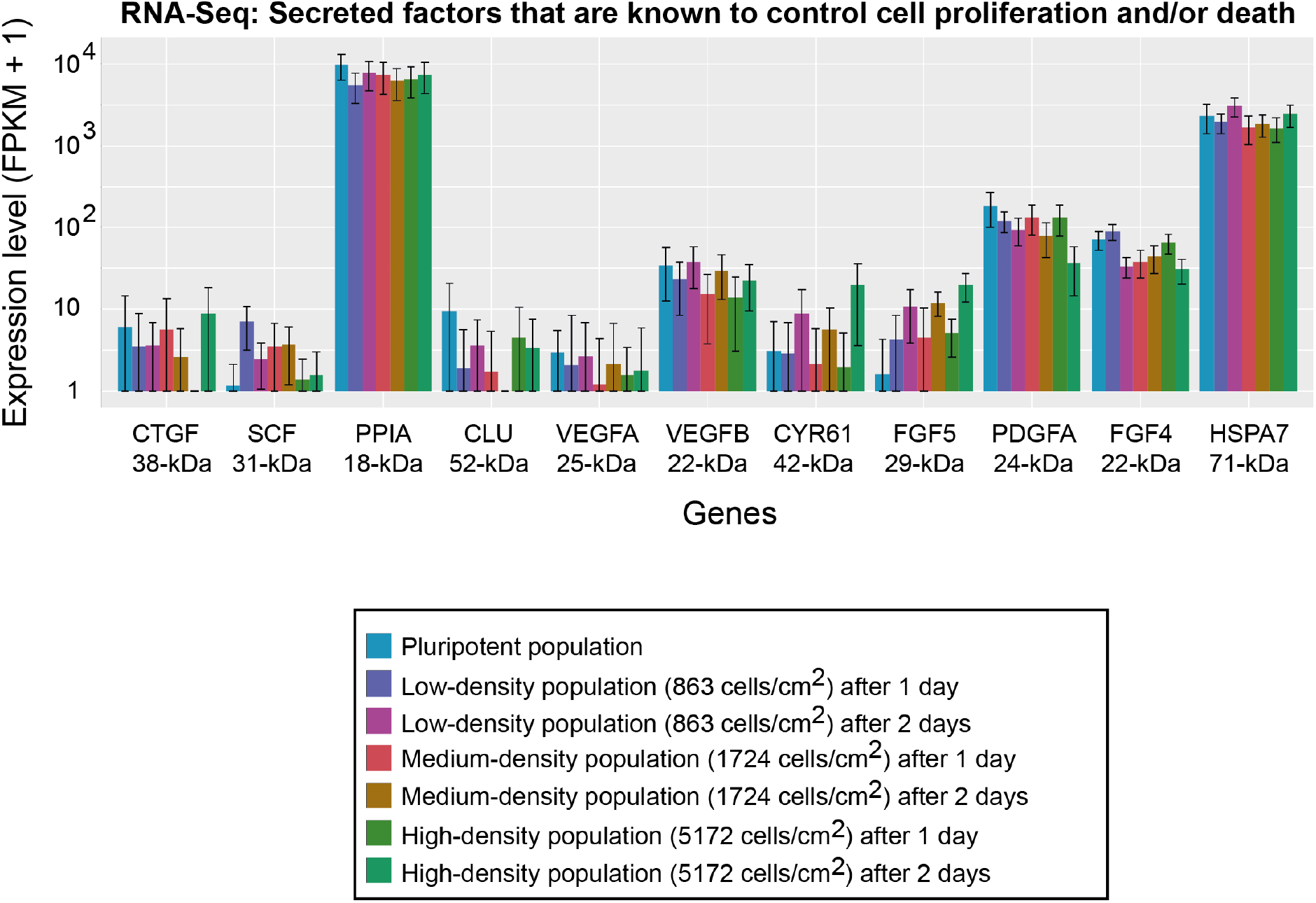
RNA-Seq: Expression levels of secreted factors that are known to control cell proliferation (related to Fig. 6). Data for 46C cells (which have Sox1 promoter driving GFP expression) differentiating towards NE lineage in N2B27 (without any inducers such as RA) that were previously self-renewing in serum+LIF (see STAR Methods). Same RNA-Seq dataset as in Fig. S18. To identify secreted factors other than FGF4 that might contribute to determining a population survival, we performed RNA-Seq to detect expression of any secreted factors that are known to control cell proliferation and/or death. We performed RNA-Seq on four populations: (1) pluripotent population prior to differentiation; (2) low-density (862 cells/cm^2^) population; (3) high-density (5172 cells/cm^2^) population; and (4) medium-density (1931 cells/cm^2^) population that is near the threshold density. For the three differentiating populations, we collected their cells on the first and second day after triggering differentiation. Expression levels (FPKMs) of secreted factors that are known to control proliferation and/or apoptosis in ES cells and that fall within the range of molecular weights that the membrane-filter experiments identified (50 – 300 kDa with +/-50% error) (Fig. 5A-B). Shown are the following genes: *Ctgf, Scf, Ppia, Clu, Vegfa, Vegfb, Cyr61, Fgf5, Pdgfa, Fgf4* and *Hspa8*. Below each gene name is the molecule’s weight (kDa) according to two online resources: Uniprot and ExPASy. *n* = 3 for all plots; Error bars are s.e.m.

**Figure S21.**
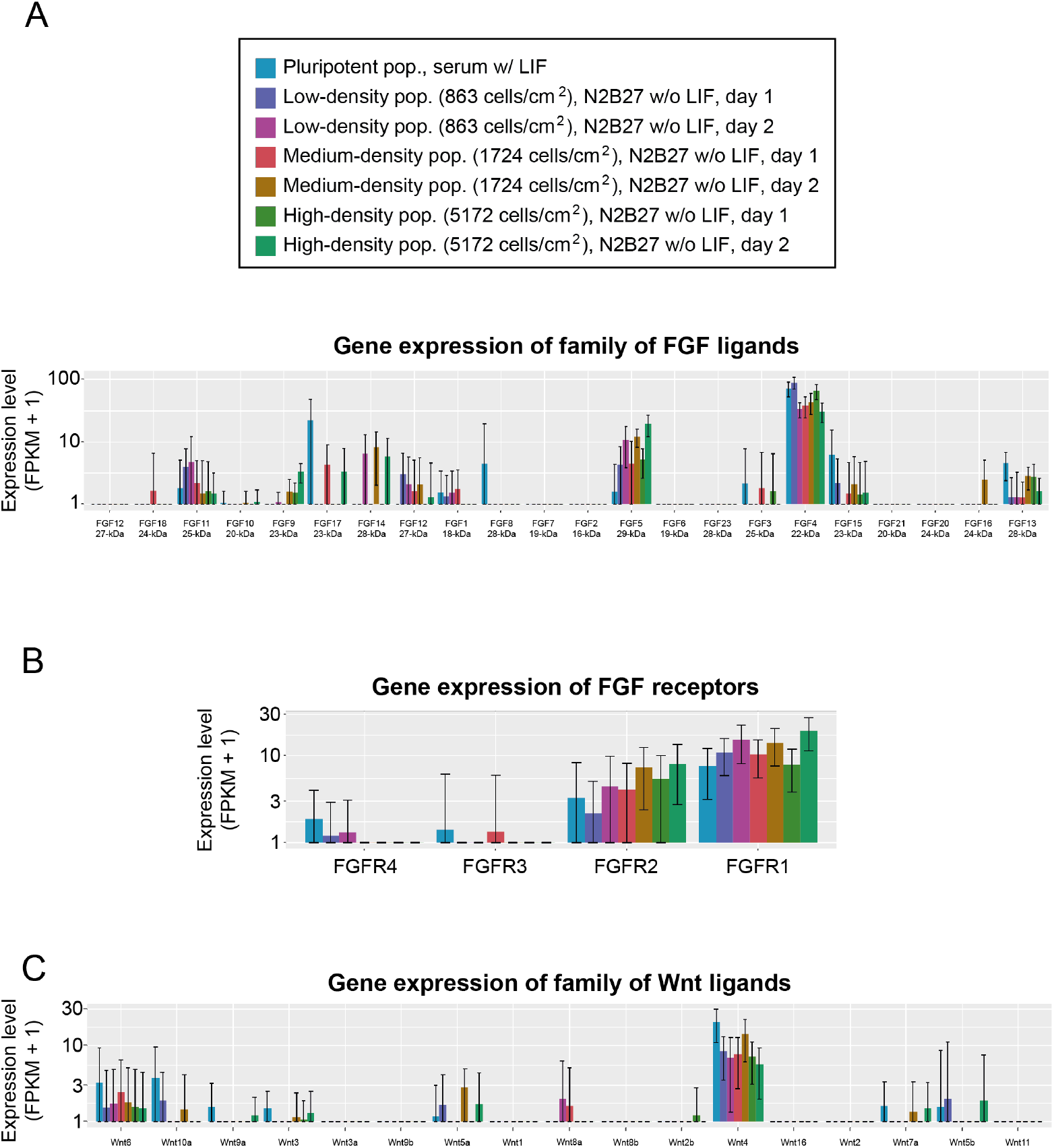
RNA-Seq: Expression levels of all 22 FGFs and their receptors (related to Fig. 6). Data for 46C cells (which have Sox1 promoter driving GFP expression) differentiating towards NE lineage in N2B27 (without any inducers such as RA) that were previously self-renewing in serum+LIF (see STAR Methods). Same RNA-Seq dataset as in Fig. S18. We performed RNA-Seq to measure the expression levels of all 22 FGF ligands (A) and their receptors (FGFRs) (B). We performed RNA-Seq on four populations: (1) pluripotent population prior to differentiation; (2) low-density (862 cells/cm^2^) population; (3) high-density (5172 cells/cm^2^) population; and (4) medium-density (1931 cells/cm^2^) population that is near the threshold density. For the three differentiating populations, we collected their cells on the first and second day after triggering differentiation. **(A)** Expression levels of all FGF ligands. Shown are the following genes: *Fgf1-8, Fgf20-21* and *Fgf23*. Below each gene name is the corresponding molecular weight in kDa. Note that FGF4 expression prominently stands out among all the FGFs. *n* = 3 for all plots; Error bars are s.e.m. **(B)** Expression levels of all FGF receptors (FGFRs). Shown are the following genes: *Fgfr1-4*. Below each gene name is the corresponding molecular weight in kDa, according to two online resources: Uniprot and ExPASy. *n* = 3 for all plots; Error bars are s.e.m. **(C)** Expression levels of Wnt ligands. Shown are the following genes: *Wnt6, Wnt10a, Wnt9a, Wnt3, Wnt3a, Wnt9b, Wnt5a, Wnt1, Wnt8a, Wnt8b, Wnt2b, Wnt4, Wnt16, Wnt7a, Wnt5b* and *Wnt11*. None of the Wnt genes prominently stand out, except for *Wnt4* which still has an order of magnitude lower expression relative to *FGF4* expression. *n* = 3 for all plots; Error bars are s.e.m.

**Figure S22.**
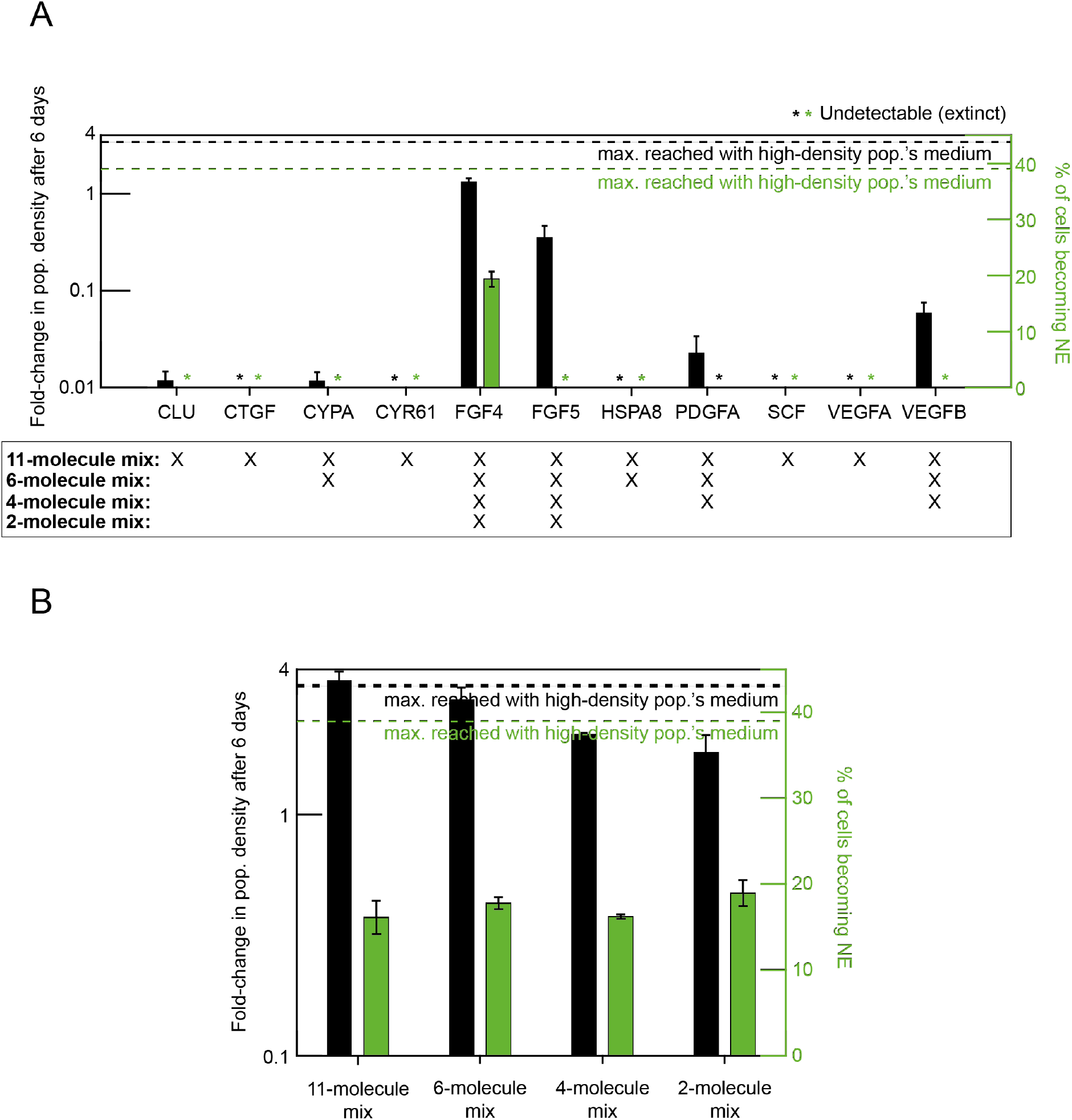
Out of all the extracellular factors that we added one-by-one into differentiation medium, only FGF4 rescues low-density populations from extinction (related to Fig. 6). The RNA-Seq (Fig. S20) revealed that 11 secreted factors that are known to control cell proliferation/and or death were highly expressed in differentiating, high-density populations. We thus reasoned that one or combinations of these factors may be the secreted molecule(s) that determine the survival-versus-extinction fate of a population. Data in (A) and (B) for 46C cells (which have Sox1 promoter driving GFP expression) differentiating towards NE lineage in N2B27+RA that were previously self-renewing in serum+LIF (see STAR Methods). **(A)** We tested these molecules by adding them one-by-one into the medium of a low-density population (862 cells/cm^2^) that would ordinarily become extinct. We added the following molecules individually, each at a saturating concentration (also see STAR Methods): version of recombinant mouse FGF4 used in Fig. S24C (200 ng/mL), recombinant human FGF5 (200 ng/mL), recombinant mouse PDGFA (100 ng/mL), recombinant mouse VEGFB 186 (100 ng/mL), recombinant mouse VEGFA (100 ng/mL), recombinant human CYR61 (500 ng/mL), recombinant human CTGF (500 ng/mL), recombinant mouse CLU (200 ng/mL), recombinant human HSPA8 (500 ng/mL), recombinant human CYPA (1000 ng/mL), and recombinant mouse SCF (2000 ng/mL). After 6 days in a medium containing one of these molecules, we measured the fold-change in density (black bars) and differentiation efficiency (green bars) of the low-density population. *n* = 3; error bars are s.e.m. These results show that only the recombinant mouse FGF4 causes the fold-change in population density to be higher than one. All the other factors resulted in the low-density population either approaching extinction (fold change much less than 1) or becoming extinct (indicated with an asterisk). The black dashed line marks the maximum fold-change in population density achieved when the low-density population grows in the medium of a high-density population. The green dashed line marks the maximum differentiation efficiency achieved when the low-density population grows in the medium of a high-density population. The box beneath the plot shows which signaling factors were mixed together and then given to the low-density population in (B). **(B)** Results obtained by giving combinations of the 11 factors together to the low-density population, with the ingredients of the mixture indicated in the box below (A). Giving all 11 factors together at once yielded the highest growth (~4-fold increase in population density; black bar), which was virtually identical to the growth obtained with a high-density population’s (5172 cells/cm^2^) medium (black dashed line). But, with the 11 molecules added together at once, the differentiation efficiency (green) remained rather low at ~20% compared to the ~40% (green dashed line) that we get from incubating the low-density population in the medium of a high-density population. As we progressively reduced the number of signaling factors in the mixture from 11 to 2, we observed only a modest decrease in population growth, down to about ~2 fold. Importantly, recombinant FGF4 was included in all these mixtures.

**Figure S23.**
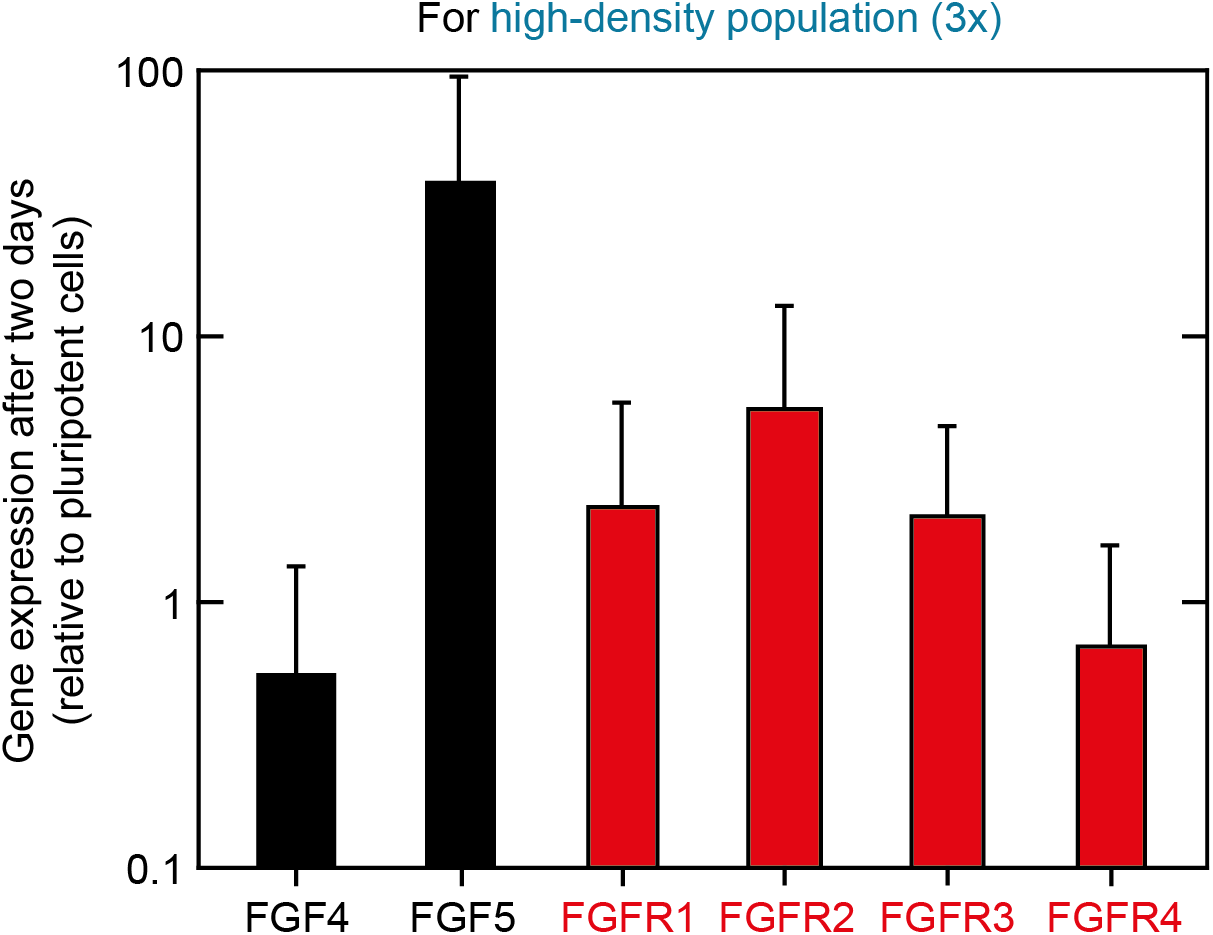
Differentiating populations express *FGF4, FGF5* and all four FGF receptor genes during the first 2 days of differentiation (related to Fig. 6). Data for 46C cells differentiating towards NE lineage in N2B27 (without any inducers such as RA) that were previously self-renewing in serum+LIF (see STAR Methods). We used real-time quantitative PCR (RT-qPCR) to measure the expression levels of all four receptors (*FGFR1-4*) of Fibroblast Growth Factors (FGFs) and the expression levels of the two FGFs, *FGF4* and *FGF5* (primers in Table S1). We examined a high-density population (5172 cells/cm^2^) after two days of differentiation. We normalized the resulting expressions of a gene relative to that of the housekeeping gene, *GAPDH* of the same population, and then further normalized the resulting value to the pluripotent population’s normalized expression level (similar to the procedure described in the caption for Fig. S30). Thus, a given gene’s expression level is compared to the pluripotent population’s expression level for that gene. Normalized expression levels of *FGF4* and *FGF5* (in black) and *FGFR1-2* (in red). *n* = 3; Error bars are s.e.m. Altogether, these results show that *FGF4, FGF5* and *FGFR1-2* are expressed - and some more so than the pluripotent population (i.e., expression value greater than 1) - during the first 2 days in which ES cells exit pluripotency.

**Figure S24.**
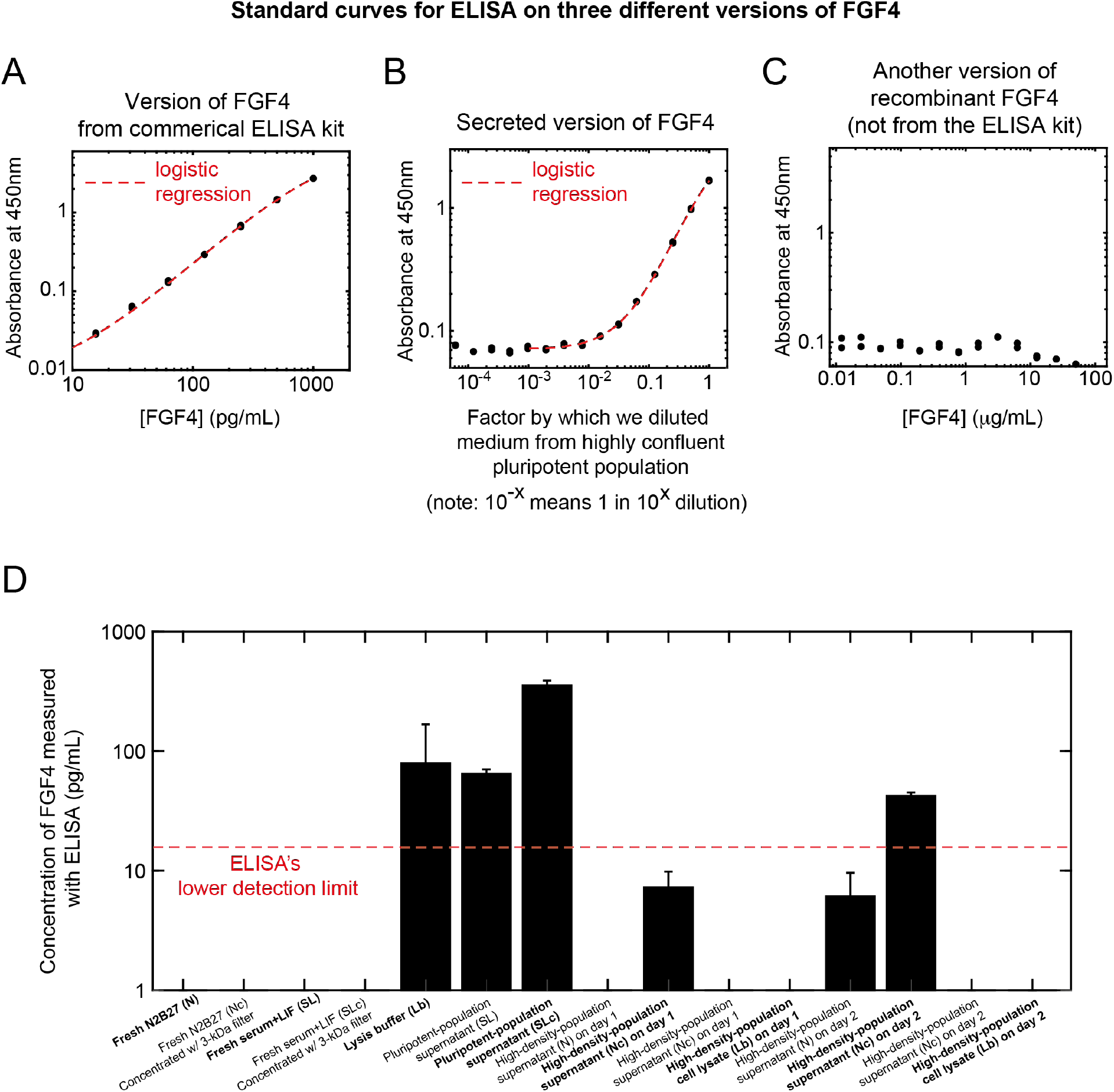
Cells secrete and extracellularly accumulate appreciable amounts of FGF4 during the first 2 days of differentiation (related to Fig. 6). We performed ELISA that detects mouse FGF4 (see STAR Methods). Data for 46C cells differentiating towards NE lineage in N2B27 (without any inducers such as RA) that were previously self-renewing in serum+LIF (see STAR Methods). **(A)** Standard curve based on a recombinant mouse FGF4 that came with the commercial ELISA kit. Note that this recombinant FGF4 is not necessarily the same version as the FGF4 that our cells secrete. Each measurement (absorbance at 450 nm) was done in duplicate (black data points). Then, we performed a logistic regression on the data by fitting a 4-parameter logistic function (red curve): 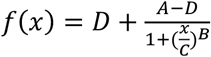, where *A, B, C* and *D* are constant coefficients and *x* is the known concentration of the recombinant mouse FGF4 that we added. We found: A = 0.0084, B = 1.313, C = 1312 and D = 6.59. **(B)** Standard curve based on the version of FGF4 that pluripotent cells secrete into their medium. We first concentrated the medium taken from a highly confluent (~80% confluent) pluripotent population with a 3-kDa filter and then performed ELISA on serially diluted fractions of this concentrated medium. Each measurement (absorbance at 450 nm) was done in duplicate (black points). Then, we performed a logistic regression on the black data points by fitting a 4-parameter logistic function (red curve): 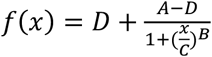, where *A, B, C* and *D* as constant coefficients and *x* as amount of lysed, pluripotent cells (day 0). We found: A = 0.07123, B = 1.184, C = 1.407 and D = 4.069. The standard curve shows that pluripotent ES cells secrete a version of FGF4 that our ELISA can detect. Moreover, it also shows a limitation of our ELISA: the assay can only detect sufficiently high concentration of FGF4 as seen by the fact that it cannot detect any FGF4 in a 1:100 dilution of a concentrated medium taken from a highly confluent ES cells. **(C)** Standard curve based on recombinant mouse FGF4 from a different manufacturer (not from the ELISA kit) that we could add to the cell-culture medium to rescue low-density populations (also see STAR Methods). Each measurement (absorbance at 450 nm) was done in duplicates (black points). As seen here, the ELISA cannot detect any amounts of this version of FGF4, even when its concentration is 100-folds higher than the highest concentration - of the version supplied by the ELISA kit - that we used in (A). The three standard curves (A-C) show that ELISA is highly sensitive to the form of FGF4 - we used three different forms in each of (A-C). The two versions of FGF4 that are not supplied by the ELISA kit (B-C) are detected with lower efficiency than the version supplied by the kit (A). **(D)** ELISA measurements of secreted FGF4 (in pg/mL) in various conditions (indicated with labels on the horizontal axis). We detected abundant FGF4 in the pluripotency medium (~500 pg/mL). In the medium of the high-density population (8620 cells/cm^2^) after two days of differentiation, we detected ~50 pg/mL of FGF4. After 1 day of differentiation, the medium of the high-density population did not contain any detectable amounts of FGF4. Hence, high-density populations take 2 days to accumulate appreciable (detectable) amounts of FGF4.

**Figure S25.**
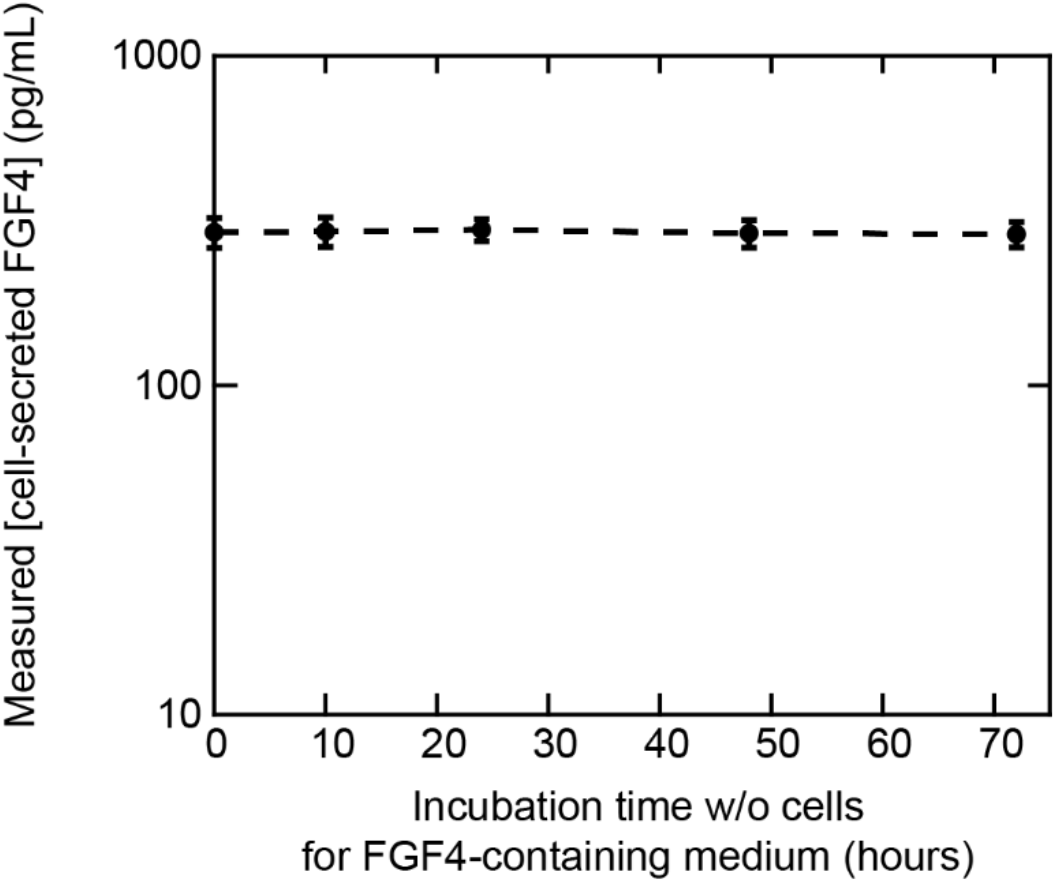
Secreted FGF4 shows no appreciable degradation for 3 days at 37°C in liquid medium (related to Fig. 6). Determining the degradation rate of FGF4 that cells secrete tells us whether FGF4 can diffuse by millimeters or not, through the Stokes-Einstein equation (see STAR Methods). We performed ELISA that targets FGF4 (see STAR Methods) to determine the concentrations of secreted FGF4 when incubated without cells in liquid medium. For this, we used a 3-kDa filter to concentration the pluripotency medium (serum+LIF) taken from a confluent population of 46C cells. Then we incubated the medium without any cells in a 37°C incubator for the hours indicated on the horizontal axis. We then took it out of the incubator and performed ELISA on it to measure the remaining [FGF4]. We observed that after incubating for 72 hours (3 days), the initial concentration of the secreted FGF4 was not appreciably degraded. As a control, we confirmed that no ingredient of a 3-kDa-concentrated pluripotency medium (without cells) interferes with our ELISA measurement to produce a non-zero concentration. Thus, this control showed no such signal (see Fig. S24D). Altogether, this result suggests that the concentration of secreted FGF4, by itself and in the absence of cells, is stable over at least 3 days and thus – according to the Stokes-Einstein equation (see STAR Methods) – can diffuse over millimeters (see STAR Methods).

**Figure S26.**
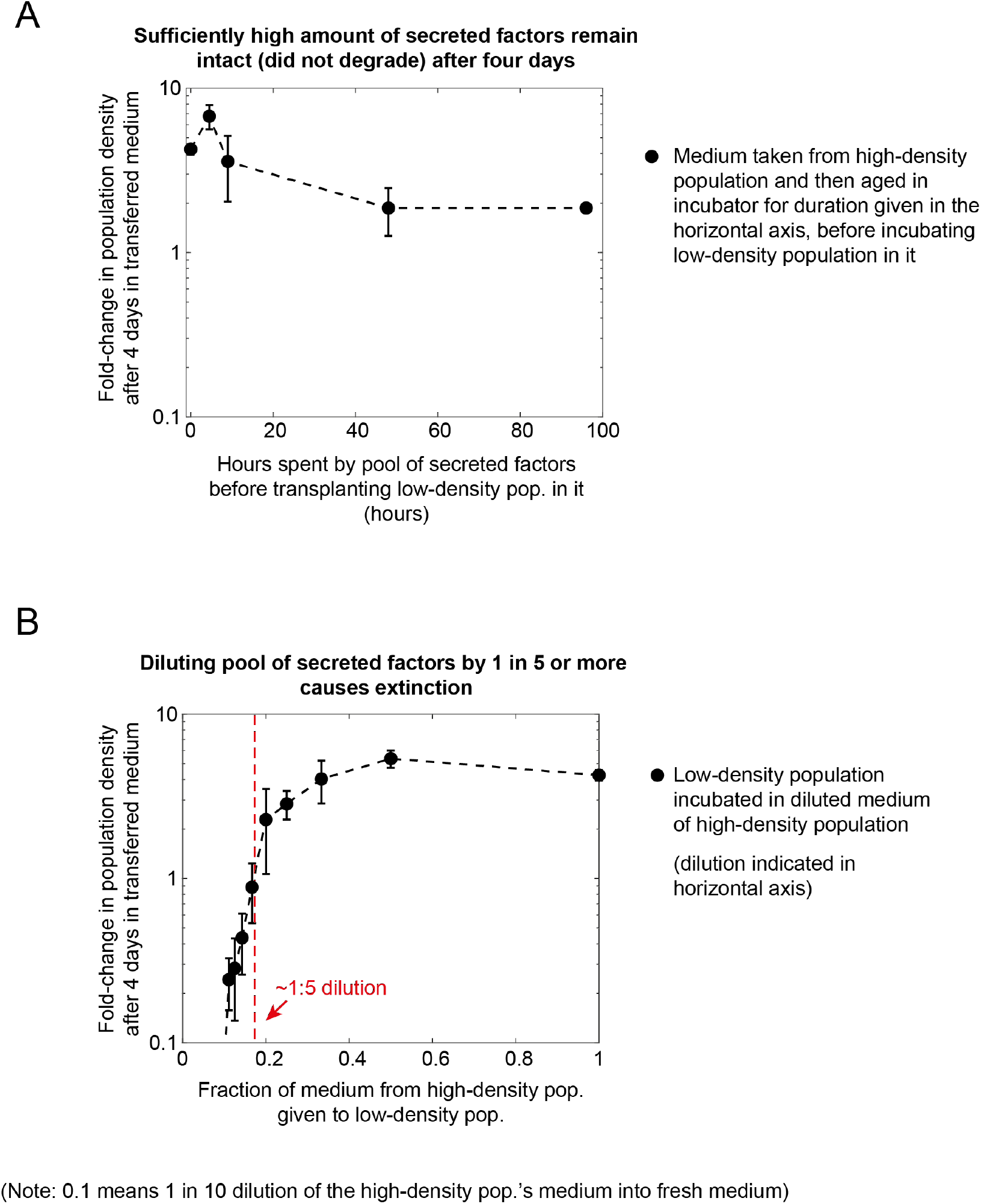
Secreted factors that control differentiating population’s survival-vs-extinction fate have a combined, effective half-life of at least 2 days (related to Fig. 6). Data for 46C cells differentiating towards NE lineage in N2B27 that were previously self-renewing in serum+LIF (see STAR Methods). According to the reaction-diffusion equation, molecules of ~100 kDa in an aqueous environment need to have a half-life of at least ~12 hours to have a diffusion length that is over 1 mm (see STAR Methods). We sought to infer the half-lives of all the secreted molecules that are important for determining the survival-versus-extinction fate of a population. What is important is actually not the individual half-life of each molecule but rather the effective half-life of all molecules combined. To determine the effect half-life of all molecules combined, we used two populations: (1) high-density population (5172 cells/cm^2^); and (2) low-density population (862 cells/cm^2^). Combining the results of (A) and (B) enabled us to infer the lower bound on the effective half-life, as we now explain. **(A)** After 2 days of differentiation with N2B27, we transferred the high-density population’s medium to an empty plate that had no cells. We incubated the medium without any cells in a 37°C incubator for various amounts of time before transferring it to a low-density population that was just ending its second day of differentiation. After 4 days of incubation in the transferred medium (so a total of 6 days of differentiation), we measured the fold-change in density of the low-density population (black points). We plotted the results here as a function of the amount of time the medium spent in the incubator without any cells before we transferred it to the low-density population. The result shows that ageing the medium for 96 hours in 37°C before transferring it to the low-density population still results in rescuing of the low-density population (fold change in population density > 1). The fold change achievable does decrease as the medium’s age increases, from ~4-fold (for unaged medium) to ~2-fold (for medium aged for ~96 hours). *n* = 3; error bars are s.e.m. **(B)** In a parallel experiment, we took the medium of the high-density population after two days of differentiation. Then, we diluted it by different amounts into a fresh differentiation medium (one that never harbored any cells). We incubated a 2-days-old low-density population into the diluted medium and then measured the foldchange in its density after four days (so a total of 6 days of differentiation). Plotted here is the fold-change in the population density as a function of how much of the medium from the high-density population was mixed with the fresh medium (e.g., 0.2 on the horizontal axis means a dilution by 1 in 5). The red line shows the maximum dilution that is allowed for still rescuing the low-density population. Any higher dilution causes the low-density population to have a fold-change in density that is less than one. *n* = 3; error bars are s.e.m. Combining the results of (A) and (B), we can conclude that more than 1/5 of the secreted molecules remain in the medium after four days of ageing in (A) since, for otherwise, the results in (B) tell us that the fold-change in (A) for medium that was aged for 96 hours should be less than 1, which is not the case. In fact, using the same reasoning, we can say that the effective, combined half-life of the secreted molecules is at least two days. To see, this, note that a half-life of one day would mean that after four days, we would have 1/16 of the molecules degraded after four days since 1/2^4^ = 1/16. But 1/16 < 1/5, which would mean that the low-density population should have become extinct in the medium that we aged for 96 hours in (A). This is not the case. Repeating the calculation by assuming that the effective, combined half-life is two days leads to: 1/2^2^ = 1/4 > 1/5, which is consistent with the data in (A). In summary, the effective, combined half-life of the secreted molecules is at least two days.

**Figure S27.**
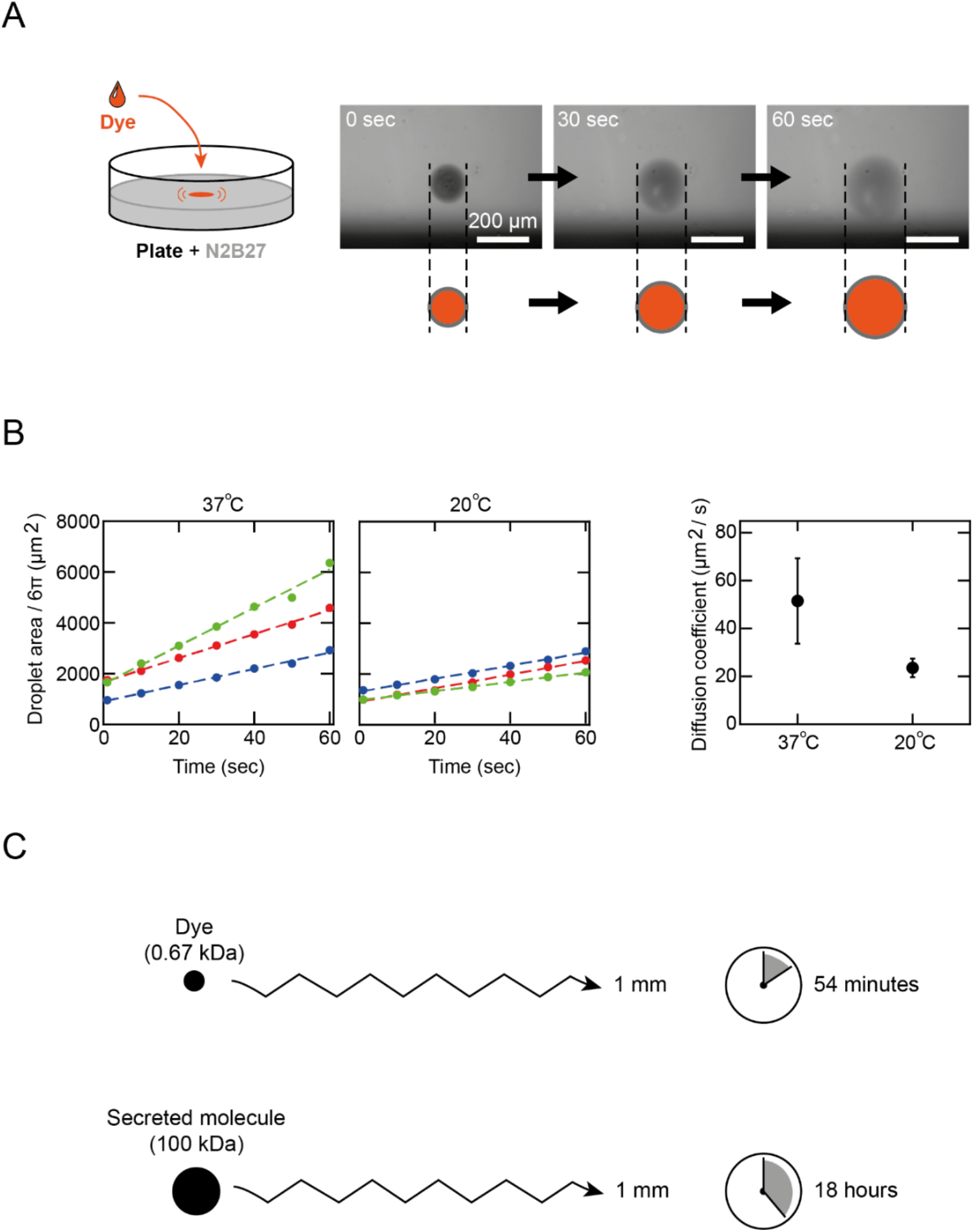
Diffusion alone, without any other mechanism of transport, explains the long-range (millimeters-scale) spreading of secreted survival-promoting factors (related to Fig. 6). To further support the idea that diffusion alone spreads the cell-secreted factors in our experiments, we determined how fast a droplet of a dye molecule of a known weight spreads in a differentiation medium without any cells, under the same incubation conditions as our cell cultures. **(A)** We used a gel loading dye (DNA Gel Loading Dye 6X, Thermo Scientific, #R0611) which consists of two molecules: bromophenol blue (669.96 Da) and xylene cyanol (538.61 Da). For simplicity, our calculations below will assume that the dye consists of only the heavier molecule, bromophenol blue. We injected a single, 0.5-μL droplet of the dye at the center of a 6-cm diameter plate that contained 5-mL of transparent N2B27 medium, either at room temperature (20°C) or pre-warmed at 37°C. We used a wide-field microscope to make a time-lapse movie with a bird’s eye view and snapshots every 10-seconds. Three snapshots (at 0, 30 and 60 seconds; all done at 37°C) of a single droplet of the dye shows the droplet expanding. Scale bar = 200 μm. **(B)** We determined the diffusion constant *D* of the dye in two ways: using the time-lapse movie and from theory. In the plots, three different colors represent three independent experiments. To determine *D* from the movies, we tracked the visible droplet boundary over time in a movie to plot the droplet area over time (shown in the two plots here at 37°C and 20°C). As shown, the droplet area linearly increased over time, which is consistent with pure diffusion (pure Brownian motion) since the area of a droplet is proportional to the mean squared displacement of a particle. Specifically, for a particle that undergoes a pure three-dimensional diffusion (Brownian motion), its mean squared displacement 〈*R*^2^〉 at time *t* is: 〈*R*^2^〉 = 6*Dt*. Let *A* be the 2-dimensionally projected area of the droplet. Then, 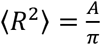 and hence, 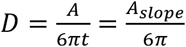, where *A_slope_* is the slope of the linear fits to the droplet area as shown in the two plots here. From these fits, the experimentally determined diffusion constants *D_exp_* at 37°C and 20°C are 51.5 ± 17.8 μm^2^/s and 23.5 ± 3.9 μm^2^/s respectively (*n* = 3; error bars are s.e.m.). As a comparison, we determined the diffusion constant *D* from theory - via the Stokes-Einstein equation which states, 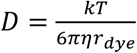 where *k* is the Boltzmann constant, *T* is temperature, *η* is the medium’s dynamic viscosity, and *r_dye_* is the radius of the dye molecule. For water, *η* = 0.000692 kg/m·s at 37°C and *η* = 0.001003 kg/m·s at 20°C (from BioNumbers (74)). We conservatively estimated *r_dye_* by noting that bromophenol blue consists of ~10 carbon-carbon bonds which would mean that the dye molecule’s diameter is 10 x 0.126 nm. For simplicity, we assume that *r_dye_* = 1*nm*. The Stokes-Einstein equation then states that the dye’s diffusion constants *D_theory_* at 37°C and 20°C are 328.1 μm^2^/s and 214.0 μm^2^/s respectively. Hence, *D_exp_* < *D_theory_*. The fact that our analysis relies on the visible (by eye) boundary of the expanding droplet would underestimate the *D_exp_* since the dye must be spreading at least as fast as the boundary does. More importantly, if there were significant convection currents in the liquid medium, then *D_exp_* would be much larger than the measured value. This argues against there being any significant liquid convection in our cell-culture media. In other words, the dye-based experiment strongly indicates that cell-secreted survival-promoting factors spread out by pure diffusion rather than by convection currents which, according to the dye, are negligible in our cell-culture conditions. Furthermore, note that 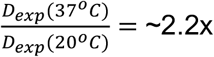 and 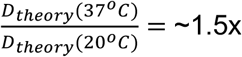; the experimental and theoretical values for pure diffusion closely match (proportional to a factor on the order of one). **(C)** Based on Brownian motion in 3 dimensions with the experimentally determined diffusion constant, the dye molecule has a mean squared displacement of 〈*R*^2^〉 = 1 mm^2^ after 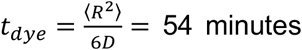 (with *D* = *D_exp_*(*dye*, 37° C) = 51.5 μm^2^/s). The same calculation, but now based on the Stokes-Einstein estimate of the diffusion constant would yield *t_dye_* = 8.5 minutes for a 1 mm^2^ mean squared displacement (with *D_theory_*(*dye*, 37°C) = 328.1 μm^2^/s). Hence the theory predicts a faster spreading of dye than experimentally observed - again, arguing against liquid convection or any other mechanism besides diffusion helping to spread the dye. A secreted molecule of 100 kDa would have a mean squared displacement of 1 mm^2^ after time, 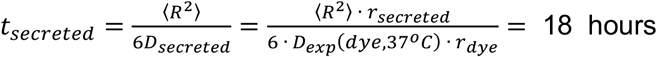, where we estimate the radius of the molecule to be *r_secreted_* = 20 nm (see STAR Methods). Hence the two days taken to observe appreciable amount of FGF4 and other survival-promoting factor(s) traveling millimeters and accumulating is consistent with the *t_secreted_* calculated here (i.e., if *t_secreted_* were much larger than two days, then we should not be observing the survival-factors travelling by millimeters within two days).

**Figure S28.**
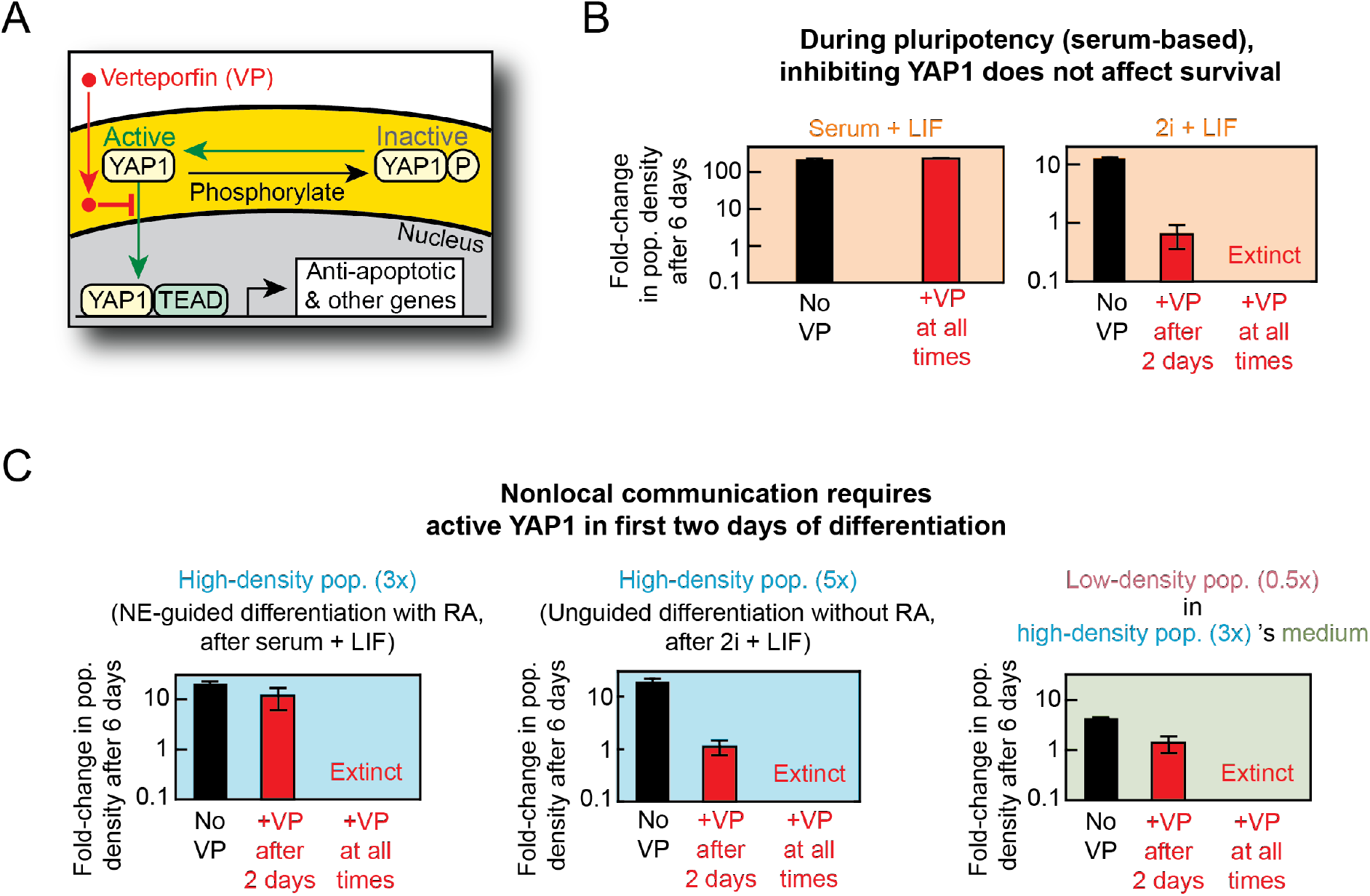
Nonlocal communication requires active YAP1 in first two days of differentiation, but during serum-based pluripotency inhibiting YAP1 does not affect survival (related to Fig. 6). **(A)** Cartoon shows YAP1 which exists as either phosphorylated (labeled “P”) or dephosphorylated. Verteporfin (VP) is a well-characterized inhibitor of active YAP1 (*60,72*) and thus inhibits active (dephosphorylated) YAP1 from entering the nucleus and regulating target gene expression and thus anti-apoptotic processes. **(B)** Data for 46C cells self-renewing in either serum+LIF (left graph) or 2i+LIF (right graph) (see STAR Methods). (Left graph) Fold-change in population density for low-density population (862 cells / cm2 initially) after 6 days of self-renewal in serum+LIF in the absence of VP (“No VP”, black bar) or presence of 1 μM VP at day 0 (“+VP at all times”, red bar). The result shows that addition of VP, and hence the inhibition of any active YAP1, does not affect survival and growth of serum-grown cells. (Right graph) Fold-change in population density for low-density population (862 cells / cm2 initially) after 6 days of self-renewal in 2i+LIF in the absence of VP (“No VP”, black bar) or presence of 1 μM VP at day 0 (“+VP at all times”, red bar) or day 2 (“+VP after 2 days”, red bar). Error bars are s.e.m.; n = 3. The result shows that addition of VP, and hence the inhibition of any active YAP1, does not affect survival and growth of only serum-grown pluripotent cells. **(C)** Data for 46C cells differentiating towards NE lineage in N2B27 with or without RA that were previously self-renewing in either serum+LIF or 2i+LIF (see STAR Methods). Left most and right most graphs are identical to the graphs shown in Fig. 6E and involved 46C cells previously self-renewing in serum+LIF and then differentiating in N2B27+RA. (Middle graph) Fold-change in population density for high-density population (8621 cells / cm2 initially, previously self-renewing in 2i+LIF) after 6 days of unguided differentiation in N2B27 without RA in the absence of VP (“No VP”, black bar) or presence of 1 μM VP at day 0 (“+VP at all times”, red bar) or day 2 (“+VP after 2 days”, red bar). Error bars are s.e.m.; n = 3. The result shows that – regardless of the use of serum during self-renewal and regardless of the use of RA during differentiation – nonlocal communication requires active YAP1 during the first two days of differentiation.

**Figure S29.**
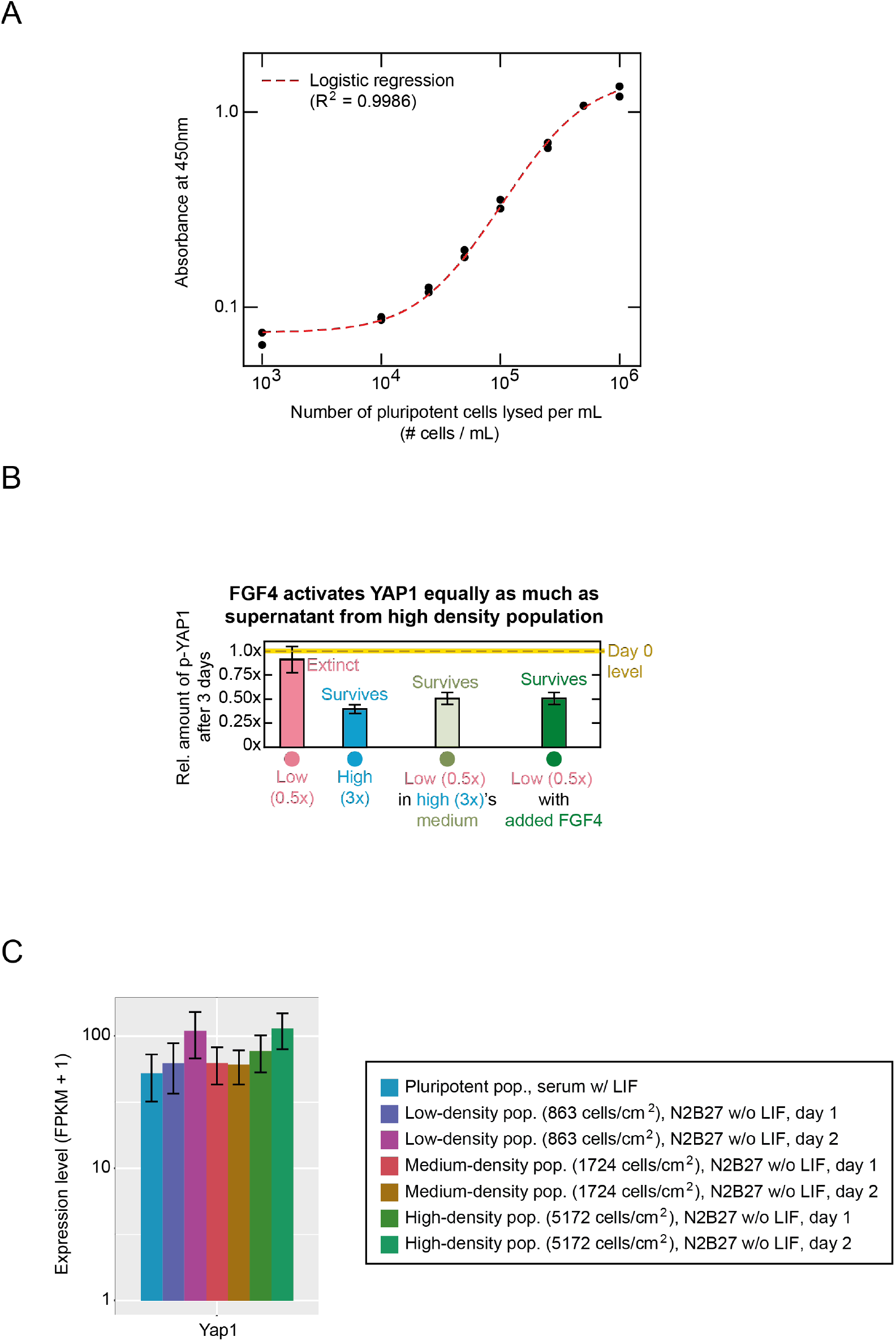
Populations that survive differentiation have more active (dephosphorylated) YAP1 compared to populations that become extinct during differentiation (related to Fig. 6). Data in (A) and (B) for 46C cells differentiating towards NE lineage in N2B27+RA that were previously self-renewing in serum+LIF (see STAR Methods). To determine how the YAP1 activity may be determined by the population density, we performed ELISA that specifically detects inactive (phosphorylated) YAP1 - YAP1 phosphorylated at Ser397, which is a primary phosphorylation site (*68,69*) (see STAR Methods). **(A)** Standard curve for the ELISA. Here we lysed pluripotent populations of various densities and then measured the amount of phosphorylated YAP1 in each lysate - this yields an optical absorbance value at 450 nm (black points). Duplicates for each lysate are shown. Red curve is the logistic fit function: 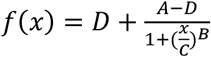, with *A, B, C* and *D* are fit constants and *x* is the number of cells lysed per mL. Results: A = 0.074, B = 1.418, C = 2.95 x 10^5^ and D = 1.502 with an R^2^ = 0.9986. **(B)** We examined three populations: (1) high-density population (5172 cells/cm^2^); (2) low-density population (862 cells/cm^2^); and (3) low-density population that we rescued from extinction by transplanting it, after two days, into the high-density population’s medium. For each population, we measured its level of phosphorylated (inactive) YAP1 three days after starting differentiation and then normalized this level to the level present in pluripotent cells of the same density (shown in (A)). This yielded a “relative abundance” for each of the three populations. Compared to the pluripotent cells of equivalent density, cells of the low-density population (pink bar) had ~10% fewer inactive YAP1 whereas cells of the high-density population (blue bar) had ~60% fewer inactive YAP1 than the pluripotent population of the same density. Cells of the rescued low-density population (green bar) had ~50% (green bar) less inactive YAP1 than pluripotent populations of the same density. Together, these results establish that, after exiting pluripotency, cells of surviving populations have more active (dephosphorylated) YAP1 than cells that head towards extinction. **(C)** Data for 46C cells differentiating towards NE lineage in N2B27 that were previously self-renewing in serum+LIF (see STAR Methods). Expression level of YAP1 from RNA-Seq dataset. Legend shows different conditions. Note that on each day, the low-density (862 cells/cm^2^ initially) and the high-density (5172 cells/cm^2^ initially) populations have virtually the same YAP1 expression level. Thus, we can compare the amounts of inactive (phosphorylated) YAP1 between the low- and high-density populations in Fig. 3E (i.e., since both populations have (nearly) the same total level of YAP1, we would be subtracting the amount of inactive YAP1 from the same value for both populations to get the amount of active Yap1). Medium-density (1931 cells/cm^2^ initially) population starts with the near-threshold density. *n* = 3 for all plots; Error bars are s.e.m.

**Figure S30.**
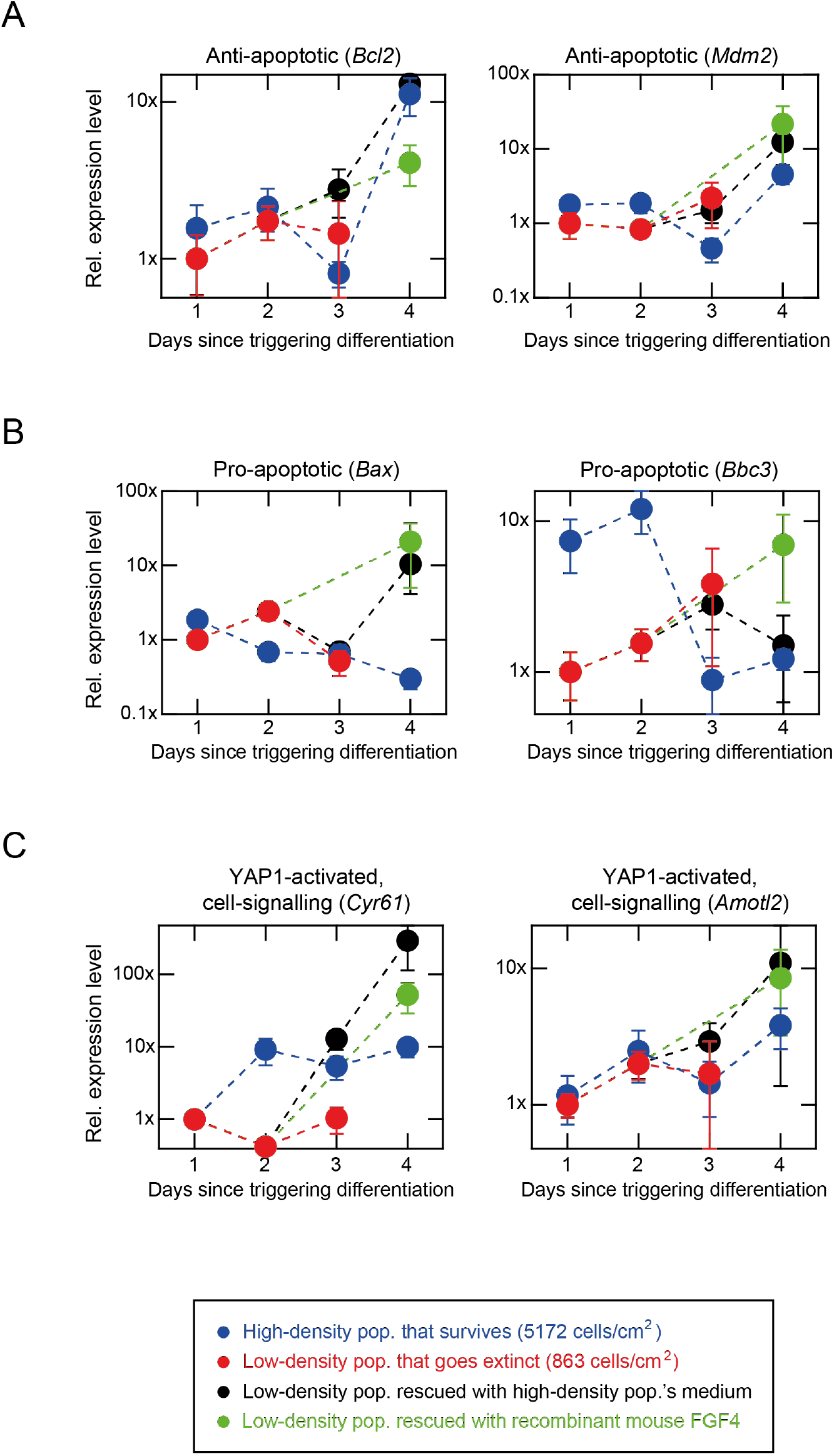
Low-density populations that are rescued from extinction with recombinant mouse FGF4 or high-density population’s medium activate anti-apoptotic pathways (e.g., *Bcl2* and *Mdm2* genes) and YAP1-controlled targets (e.g., *Cyr61* and *Amotl2* genes) (related to Fig. 6). Data for 46C cells differentiating towards NE lineage in N2B27+RA that were previously self-renewing in serum+LIF (see STAR Methods). With realtime quantitative PCR (primers in Table S1), we measured anti-apoptotic, pro-apoptotic, and YAP1-mediated cell-signaling genes over the course of differentiation. Normalization of expression values: for each gene *g*, we first divided its expression level by the expression level of *Gapdh*, resulting in a value *N_g_*. For each population, we divided its *N_g_* by the low-density population’s *N_g_* on day 1 to get the final, normalized expression level *μ* which is plotted in here in all graphs. Thus, “1x” is the expression level of the low-density population on the first day after starting differentiation. We examined four populations: (1) high-density population (5172 cells/cm^2^); (2) low-density population (862 cells/cm^2^); (3) low-density population that we rescued from extinction by transplanting it, after two days, into the high-density population’s medium; and (4) low-density population that we rescued after adding 200 ng/mL recombinant mouse FGF4 to its differentiation medium on day 0. **(A)** Expression levels of two anti-apoptotic genes, *Bcl2* (left graph) and *Mdm2* (right graph). Data for *Bcl2* is a replicate of the data shown in Fig. 6G which we show here for comparison with *Mdm2*. Both *Bcl2* and *Mdm2* show increased expressions (more anti-apoptotic) for high-density population (blue) and low-density population that was rescued by the medium of the high-density population after the 2nd day (black) or with FGF4 (green). Low-density population that goes extinct (red) shows nearly constant, low expression level of both genes. No data for 4th day is shown for the low-density population because it becomes extinct after the 3rd day (there were already barely any cells left for the 3rd day data shown here). **(B)** Expression levels of two pro-apoptotic genes - *Bax* (left graph) and *Bbc3* (right graph). Color scheme is the same as in (A). The high-density population initially has a higher Bbc3 expression than the low-density population but eventually down-regulates and has lower Bbc3 expression than the low-density population. The low-density population, in turn, gradually increases its Bbc3 expression over time, up to the moment of extinction (~ day 3). Note that the rescued low-density population keeps its Bbc3 expression level low, past day 2 (which is when it receives the medium from a high-density population) and has nearly same low Bbc3 expression as the high-density population after being rescued. Note that differentiation is known to increase expression of apoptotic genes. **(C)** Expression levels of two cell-signaling genes that are upregulated by Yap1, *Cyr61* (left graph) and *Amotl2* (right graph). Data for *Cyr61* is a replicate of the data shown in Fig. 6G. Only the high-density and the rescued low-density populations gradually increase the expression levels of both genes whereas the low-density population that heads towards extinction (red) maintains a nearly constant, low expression of both genes (consistent with our findings in Fig. 3 that secreted factors that are abundant for high-density populations increase YAP1 activity (and thus upregulate expression of *Cyr61* and *Amotl2*). In all the plots, *n* = 3; Error bars are s.e.m.

**Figure S31.**
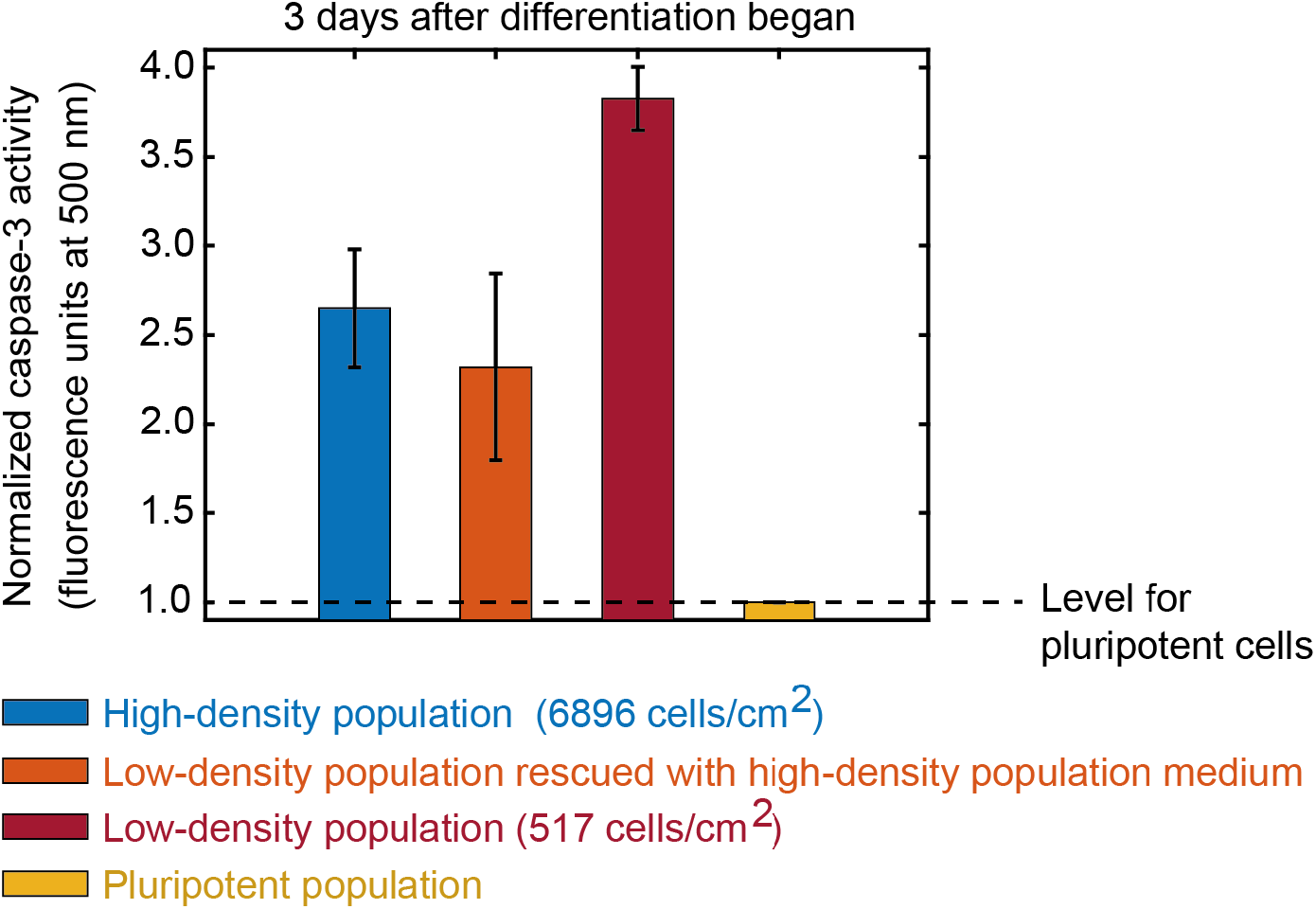
Pro-apoptotic caspase-3 activity is higher in low-density populations than in high-density and rescued low-density populations (related to Fig. 6). Data for E14 cells differentiating towards NE lineage in N2B27+RA that were previously self-renewing in serum+LIF (see STAR Methods). We sought to examine the activity of a well-known, pro-apoptotic marker (caspase 3) in populations that are either extinction-bound or surviving. We used a membrane-permeable, DNA-dye-based assay (NucView 488 Caspase-3 Assay Kit for Live Cells). This assay measured the amounts of active caspase 3/7 inside cells. We examined caspase-3 levels in four different populations: (1) high-density population (6896 cells/cm^2^ -blue bar); low-density population (517 cells/cm^2^ -red bar); (3) low-density population that was rescued from extinction by transplanting it, after two days, into the high-density population’s medium (orange bar); and (4) pluripotent E14 cells before we induced the differentiation (yellow bar). After 3 days of differentiation, we collected the cells from each of these populations, mixed them with the DNA-dye according to the manufacturer’s protocol, and then measured the resulting fluorescence at 500 nm in single cells with a flow cytometer. Plotted here are the geometric means of the fluorescence for each population. Higher fluorescence means more caspase-3 activity. We normalized the values to the fluorescence level of the pluripotent population (yellow bar), as indicated by the dashed line. *n* =3; Error bars are s.e.m. This plot shows that all differentiating populations upregulated levels of caspase-3 activity relative to the pluripotent populations. Thus, the nonlocal communication causes populations to have less caspase-3 activities than extinction-bound populations.

**Table S1.**
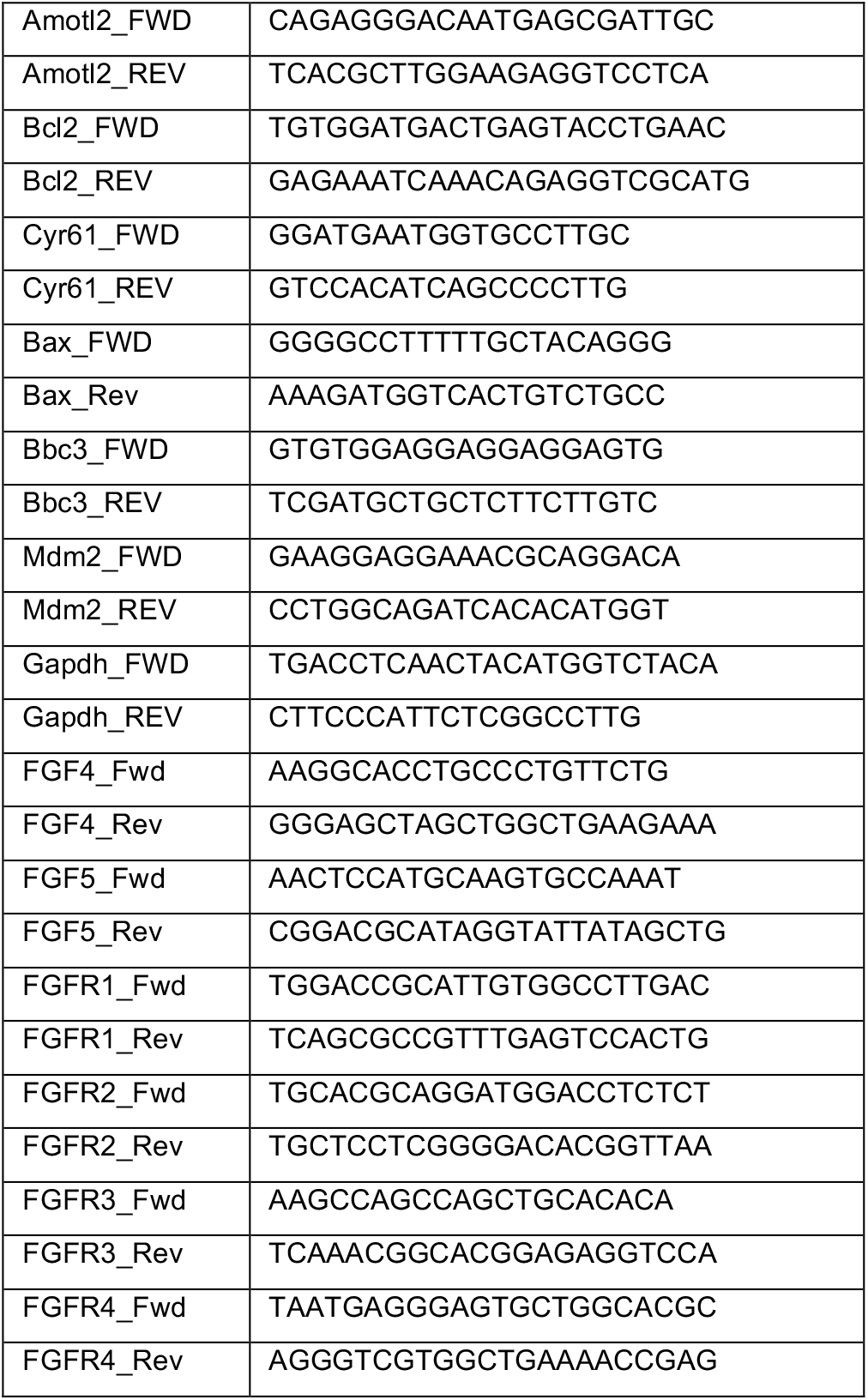
List of primers used for RT-qPCR (Related to Fig. 6) Below we summarized the forward (FWD) and reverse (REV) primers of the genes used for RT-qPCR. Also see STAR Methods for procedure.

**Table S2.** Ingredients for the differentiation medium (N2B27) and their molecular weights (Related to Fig. 5). Our recipe for making N2B27 (differentiation medium) is based on established protocols (Ying et al (*51*)) and is summarized in the STAR Methods. Below we list the ingredients of N2B27 together with their molecular weights (MW). This list is adapted from Mittal and Voldman (*16*) and based on information from ATCC, Ying et al. (*51*), and Brewer et al. (*78*). Most components of differentiation medium (N2B27) are smaller than the smallest filter size that we used (3 kDa), and if larger then not vital for growth of ES cells (see Mittal and Voldman (*16*)). We show experimentally that filtering N2B27 does not catch any ingredients vital for ES cell growth (see Fig. S15) Abbreviations: MW = molecular weight, Da = Daltons, N/A = not applicable (i.e., not included in mixture), CONF = confidential, propriety information (Invitrogen).

| Compound Name | MW (Da) | Conc. in DMEM/F-12 (μM) | Conc. in Neurobasal (μM) | Conc. In N2 supplement (μM) | Conc. In B-27 minus vitamin-A supplement (μM) |
| --- | --- | --- | --- | --- | --- |
| *Inorganic salts* |  |  |  |  |  |
| CaCl<sub>2</sub> (anhydrous) | 111 | 1000 | 1800 | N/A | N/A |
| CuSO<sub>4</sub> (anhydrous) | 160 | 0.01 | N/A | N/A | N/A |
| Fe(NO<sub>3</sub>)<sub>3</sub>·9H<sub>2</sub>O | 404 | 0.13 | 0.25 | N/A | N/A |
| FeSO<sub>4</sub>·7H<sub>2</sub>O | 278 | 1.5 | N/A | N/A | N/A |
| MgSO<sub>4</sub> (anhydrous) | 120 | 700 | 812 | N/A | N/A |
| KCl | 75 | 4000 | 5333 | N/A | N/A |
| NaHCO<sub>3</sub> | 84 | 14,300 | 26,000 | N/A | N/A |
| NaCl | 58 | 120,000 | 51,300 | N/A | N/A |
| Na<sub>2</sub>HPO<sub>4</sub> (anhydrous) | 142 | 500 | N/A | N/A | N/A |
| NaH<sub>2</sub>PO<sub>4</sub> ·H<sub>2</sub>O | 120 | 500 | 1,000 | N/A | N/A |
| ZnSO<sub>4</sub>·7H<sub>2</sub>O | 288 | 1.5 | N/A | N/A | N/A |
| *Amino Acids* |  |  |  |  |  |
| L-Alanine | 89 | 50 | 22.5 | N/A | N/A |
| L-Arginine·HCl | 210 | 700 | 400 | N/A | N/A |
| L-Asparagine·H<sub>2</sub>O | 150 | 50 | 5 | N/A | N/A |
| L-Aspartic Acid | 133 | 50 | N/A | N/A | N/A |
| L-Cystine·HCl·H<sub>2</sub>O | 294 | 60 | N/A | N/A | N/A |
| L-Cystine·2HCl | 312 | 100 | 10 | N/A | N/A |
| L-Glutamic Acid | 147 | 50 | N/A | N/A | N/A |
| L-Glutamine | 146 | 2,500 | 500 | N/A | N/A |
| Glycine | 75 | 250 | 400 | N/A | N/A |
| L-Histidine·HCl·H<sub>2</sub>O | 209 | 150 | 200 | N/A | N/A |
| L-Isoleucine | 131 | 415 | 800 | N/A | N/A |
| L-Leucine | 131 | 450 | 800 | N/A | N/A |
| L-Lysine·HCl | 146 | 625 | 1,000 | N/A | N/A |
| L-Methionine | 149 | 115 | 200 | N/A | N/A |
| L-Phenylalanine | 165 | 215 | 400 | N/A | N/A |
| L-Proline | 115 | 150 | 67 | N/A | N/A |
| L-Serine | 105 | 250 | 400 | N/A | N/A |
| L-Threonine | 119 | 450 | 800 | N/A | N/A |
| L-Tryptophan | 204 | 45 | 80 | N/A | N/A |
| *Vitamins* |  |  |  |  |  |
| D-Biotin (B7) | 244 | 0.01 | CONF | N/A | N/A |
| Choline Chloride | 140 | 64 | 28 | N/A | N/A |
| Folic Acid (B9) | 441 | 6 | 9 | N/A | N/A |
| myo-Inositol (B8) | 180 | 70 | 40 | N/A | N/A |
| Niacinamide (B3) | 122 | 16.5 | 33 | N/A | N/A |
| D-Pantothenic Acid (B5) | 219 | 10 | 8 | N/A | N/A |
| Pyridoxine-HCl (B6) | 202 | 10 | 20 | N/A | N/A |
| Riboflavin (B2) | 376 | 0.6 | 1 | N/A | N/A |
| Thiamine-HCl (B1) | 373.5 | 6 | 10 | N/A | N/A |
| Vitamin B-12 | 1,355 | 0.5 | 0.25 | N/A | N/A |
| *Proteins* |  |  |  |  |  |
| Insulin | 5,800 | N/A | N/A | 4.3 | CONF |
| Serotransferrin | 77,000 | N/A | N/A | 1.33 | CONF |
| BSA | 66,000 | N/A | N/A | 0.75 | CONF |
| SOD-1 (Dimer) | 32,000 | N/A | N/A | N/A | CONF |
| Catalase (Tetramer) | 240,000 | N/A | N/A | N/A | CONF |
| *Other* |  |  |  |  |  |
| D-Glucose | 180 | 17,500 | 25,000 | N/A | N/A |
| HEPES | 238 | 15,000 | 10,000 | N/A | N/A |
| Hypoxanthine | 136 | 17.5 | N/A | N/A | N/A |
| Linoleic Acid | 280 | 0.16 | N/A | N/A | N/A |
| Phenol Red, Sodium Salt | 354 | 23 | 23 | N/A | N/A |
| Putrescine-2HCl | 161 | 0.5 | N/A | N/A | N/A |
| Pyruvic Acid-Na | 111 | 500 | 225 | N/A | N/A |
| DL-Thioctic Acid | 206 | 0.5 | N/A | N/A | N/A |
| Thymidine | 242 | 1.5 | N/A | N/A | N/A |
| Progesterone | 314 | N/A | N/A | 0.02 | CONF |
| Sodium Selenite | 173 | N/A | N/A | 0.02 | CONF |
| Putrescine | 88 | N/A | N/A | 180 | CONF |
| Lecithin | 770 | N/A | N/A | N/A | CONF |
| Linolenic Acid | 278 | N/A | N/A | N/A | CONF |
| Phosphatidylcholine | 776 | N/A | N/A | N/A | CONF |
| Retinol | 286 | N/A | N/A | N/A | CONF |
| Retinyl Acetate | 328 | N/A | N/A | N/A | CONF |
| Sodium Selenite | 173 | N/A | N/A | N/A | CONF |
| T3 (Triodo-L-Thyronine) | 650 | N/A | N/A | N/A | CONF |
| DL-α-Tocopherol (Vitamin E) | 431 | N/A | N/A | N/A | CONF |
| DL-α-Trocopherol<br>Acetate | 473 | N/A | N/A | N/A | CONF |
| L-Tyrosine·2Na·2H<sub>2</sub>O | 263 | 200 | 400 | N/A | N/A |
| L-Valine | 117 | 450 | 800 | N/A | N/A |

